# Tracing the origin of a new organ by inferring the genetic basis of rumen evolution

**DOI:** 10.1101/2020.02.19.955872

**Authors:** Xiangyu Pan, Yu Wang, Zongjun Li, Xianqing Chen, Rasmus Heller, Nini Wang, Chen Zhao, Yudong Cai, Han Xu, Songhai Li, Ming Li, Cunyuan Li, Shengwei Hu, Hui Li, Kun Wang, Lei Chen, Bin Wei, Zhuqing Zheng, Weiwei Fu, Yue Yang, Tingting Zhang, Zhuoting Hou, Yueyang Yan, Xiaoyang Lv, Wei Sun, Xinyu Li, Shisheng Huang, Lixiang Liu, Shengyong Mao, Wenqing Liu, Jinlian Hua, Zhipeng Li, Guojie Zhang, Yulin Chen, Xihong Wang, Qiang Qiu, Brian P. Dalrymple, Wen Wang, Yu Jiang

**Affiliations:** Key Laboratory of Animal Genetics, Breeding and Reproduction of Shaanxi Province, College of Animal Science and Technology, Northwest A&F University, Yangling 712100, China; School of Ecology and Environment, Northwestern Polytechnical University, Xi’an 710072, China; Section for Computational and RNA Biology, Department of Biology, University of Copenhagen, DK-2100 Copenhagen, Denmark; Marine Mammal and Marine Bioacoustics Laboratory, Institute of Deep-sea Science and Engineering, Chinese Academy of Sciences, Sanya 572000, China.; College of Life Sciences, Shihezi University, Shihezi, Xinjiang 832003, China; College of Animal Science and Technology, Yangzhou University, Yangzhou 225009, China; Joint International Research Laboratory of Agriculture and Agri-Product Safety of Ministry of Education of China, Yangzhou University, Yangzhou 225009, China; Key Laboratory for Major Obstetric Diseases of Guangdong Province, The Third Affiliated Hospital of Guangzhou Medical University, Guangzhou 510150, China; School of Life Science and Technology, Shanghai Tech University, Shanghai 201210, China; College of Animal Science and Technology, Nanjing Agricultural University, Nanjing 210095, China; College of Veterinary Medicine, Shaanxi Centre of Stem Cells Engineering & Technology, Northwest A&F University, Yangling, Shaanxi 712100, China; Department of Special Economic Animal Nutrition and Feed Science, Institute of Special Animal and Plant Sciences, Chinese Academy of Agricultural Sciences, Changchun 130112, China; Section for Ecology and Evolution, Department of Biology, University of Copenhagen, DK-2100 Copenhagen, Denmark; China National GeneBank, BGI-Shenzhen, Shenzhen 518083, China; State Key Laboratory of Genetic Resources and Evolution, Kunming Institute of Zoology, Chinese Academy of Sciences, Kunming 650223, China; Center for Excellence in Animal Evolution and Genetics, Chinese Academy of Sciences, Kunming 650223, China; State Key Laboratory of Grassland Agro-Ecosystem, College of Life Sciences, Lanzhou University, Lanzhou 730000, China; School of Animal Biology and Institute of Agriculture, The University of Western Australia, 35 Stirling Highway, Crawley WA 6009, Australia

## Abstract

The rumen is the hallmark organ of ruminants and hosts a diverse ecosystem of microorganisms that facilitates efficient digestion of plant fibers. We used 897 transcriptomes from three Cetartiodactyla lineages: ruminants, camels and cetaceans, as well as data from ruminant comparative genomics and functional assays to explore the genetic basis of rumen origin and evolution. Comparative analyses reveal that the rumen and the first-chamber stomachs of camels and cetaceans shared a common tissue origin from the esophagus. The rumen recruited genes from other tissues/organs and up-regulated many esophagus genes to aquire functional innovations involving epithelium absorption, improvement of the ketone body metabolism and regulation of microbial community. These innovations involve such genetic changes as ruminant-specific conserved elements, newly evolved genes and positively selected genes. Our *in vitro* experiements validate the functions of one enhancer, one positively selected gene and two newly evolved antibacterial genes. Our study provides novel insights into the origin and evolution of a complex organ.

---

Evolutionary biology has a long history of trying to understand how complex organs evolve^1^. The origin of some notable organs has been central to animal evolution, e.g. the eyes of animals^2,3^, electric organs of fishes^4^, mammalian placenta^5,6^ and ruminant headgear^7^. Another remarkable organ innovation found in mammals are the multi-chambered stomachs found in the Cetartiodactyla lineages, including Tylopoda (e.g. camels), Tayassuidae (e.g. peccaries), Hippopotamidae (e.g. hippos), Cetacea (e.g. whales) and Ruminantia (Fig. 1). Among these, ruminants have the most complex digestive system in herbivores, allowing efficient uptake of nutrients from plant material by providing a microbial fermentation ecosystem in the highly specialized rumen^8^. Camels (Tylopoda) have three-chambered stomachs and are also sometimes called "pseudo-ruminants" due to their similar ruminating behavior and microbial fermentation taking place in their first-chamber (FC) stomach^9^. The whales (Cetacea) form the sister group of the Ruminantia^10^, however the FC of their four-chambered stomach is mainly used as a temporary storage chamber for ingested food and for mechanical grinding of food items^11^. With the rumen, ruminants obtained a unique evolutionary advantage through superior utilization of short chain fatty acids (SCFAs) from microbial fermentation, which significantly promoted the expansion and diversification of ruminant taxa^12^. The evolutionary innovation of the rumen is therefore interesting not only in its functional complexity and uniqueness, but also because it has greatly benefited humans by providing high-quality nutrition in the shape of highly productive ruminant livestock species^13,14^.

**Fig. 1.**
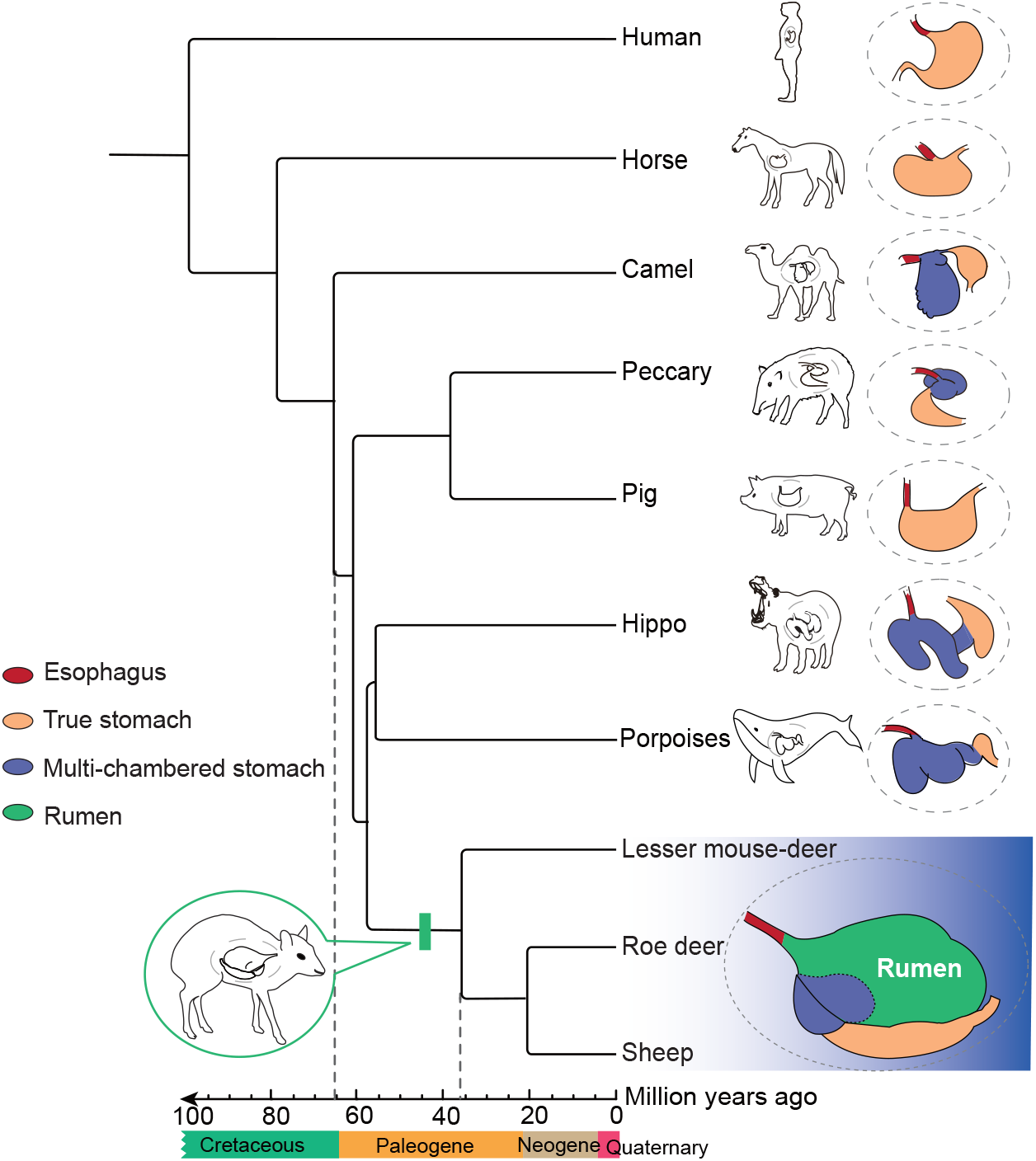
Origin of the rumen. Maximum-likelihood (ML) tree generated using 3,316,385 four-fold degenerate sites with 11,567 single-copy orthologous genes. Dates for major events are taken from the TimeTree Database^46^ and Chen *et al*.,^30^. The green rectangular block indicates the Ruminantia. Dotted lines link to the detailed divergence times of the two taxa. The esophagus is colored red, the additional stomach chambers in the multi-stomach lineages purple, the rumen green, and the true stomach/abomasum orange.

The anatomical predecessor from which the rumen evolved has been proposed to be the esophagus^15^, yet the two organs are highly divergent in morphology and physiology. The stratified squamous epithelium of the esophagus is smooth and non-keratinized, and mainly serves a barrier function, but in contrast the rumen stratified squamous epithelium is keratinized and lined with papillae, which facilitates nutrient uptake and antibacterial peptide production^16,17^. These features allow the absorption of SCFAs and sustain the homeostasis of microorganisms. The origin and evolution of new organs involve structural and functional innovations that were proposed to be driven by several types of genetic reprogramming: recruitment of genes usually expressed in other organs, transformation of regulatory elements such as promoters and enhancers, mutations in protein-coding genes and post-transcriptional mechanisms^1,5^. Given the substantial structural and physiological changes involved in the transition from esophagus to rumen, significant genetic reprogramming must have occurred during the process.

Usually, it is challenging to obtain detailed insights into the genetic reprogramming associated with organ evolution due to the rarity of such occurrences and the lack of intermediate evolutionary states^5^. However, in the case of the rumen, we can take advantage of two important points allowing “triangulation” of the changes leading to the rumen: the availability of synapomorphic stomach chambers in Cetartiodactyla and the likely ancestral relation between the esophagus and the rumen. Here, we conducted a comprehensive comparison using 897 transcriptomes of different tissues from three Cetartiodactyla lineages and multiple genomes to investigate the genetic basis of gene programming evolution and functional innovations in rumen, together with validation of some cases using *in vitro* experiments.

## Results

### Gene expression features of the rumen

We sequenced transcriptomes of 33 samples across 14 adult tissues from Bactrian camels, eight adult tissues from one species in Mysticeti (Bryde’s whale) and one species in Odontoceti (Indo-Pacific Finless Porpoise) from Cetacea, 852 samples (210 sequenced in this study and 642 published in previous studies^7,18,19^) from 50 tissues of two representative ruminants (sheep and roe deer) within Ruminantia (**Supplementary Table 1**). The global gene expression patterns of all the FC stomachs are consistently most similar to the esophagus in all species (Fig. 2a, **Fig. S1**). To investigate the specifically expressed genes in the three types of FC stomachs, we defined those that the rank of expression is less than or equal to a E50 index threshold with type I error less than 0.05 (**Supplementary Note**) in the FC stomachs of ruminants, camels, and cetaceans compared to other conspecific tissues/organs. We identified 655, 593, and 375 such specifically expressed genes in the FC stomachs of ruminants, camels, and cetaceans, respectively (**Supplementary Table 2-4; Supplementary Note**).

**Fig. 2.**
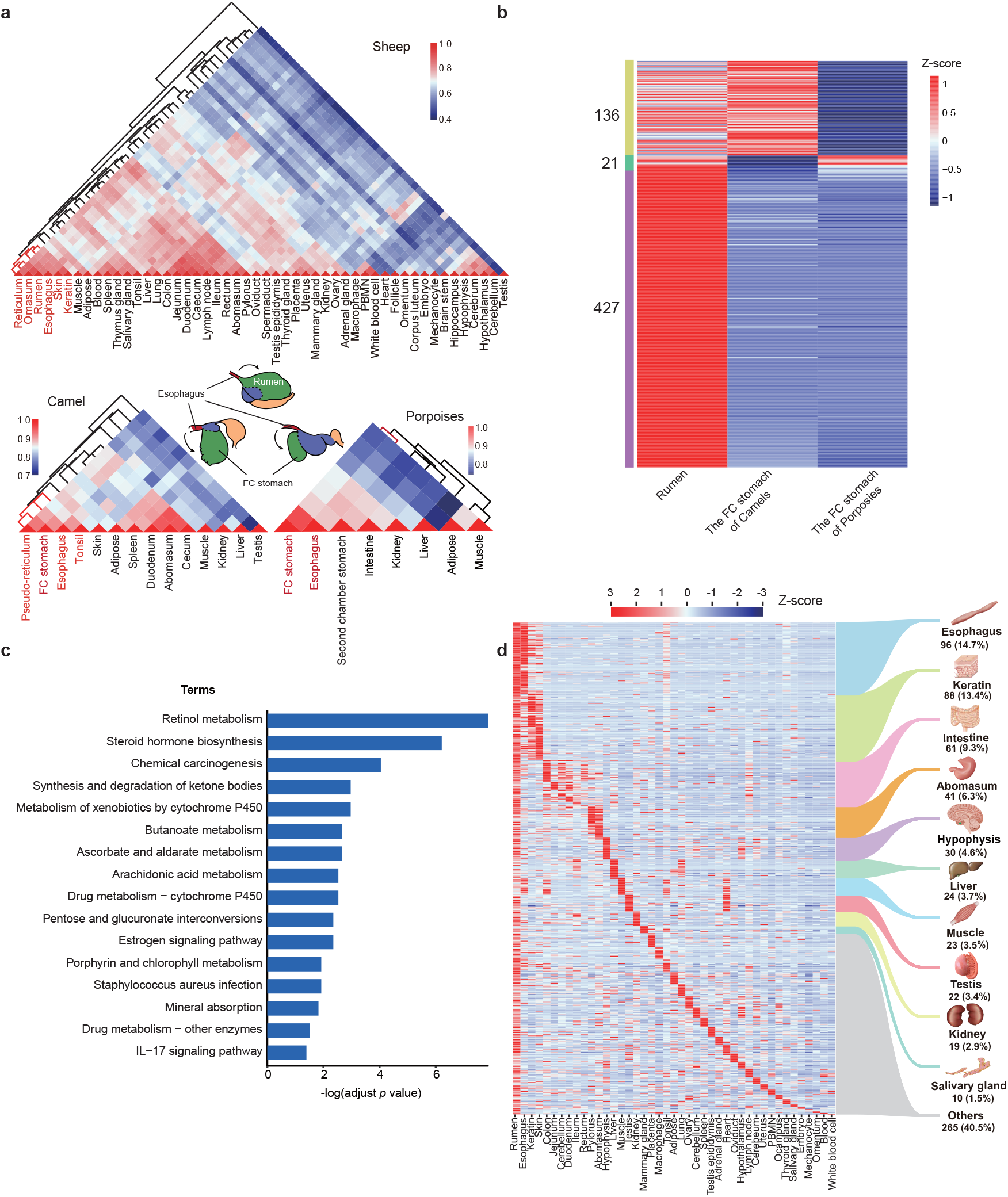
Comparisons of gene expression profile among rumen and other tissues. **a**, Hierarchical clustering results showing the relationships among 50 tissues of sheep and a heatmap showing the pairwise Spearman correlations between sheep tissues(the top triangle), between 14 tissues of camels (lower left triangle) and between eight tissues of two cetaceans (lower right triangle). **b**, Heatmap of differentially expressed rumen specifically expressed genes among the rumen and other FC stomachs. The color bars on the left present 136 DEGs of the rumen relative to the FC stomach of cetaceans (yellow), 21 DEGs relative to the FC stomach of camels (green), and 427 DEGs relative to the FC stomach of both species (purple). The expression levels were normalized by Z-scores. **c**, KEGG pathway analysis of 427 rumen up-regulated DEGs relative to both the FC stomach of camels and cetaceans. **d**, Heatmap showing the gene expression profiles of all 655 rumen specifically expressed genes across 43 tissues of sheep. Different colored lines represent the tissues from which the rumen specifically expressed genes were recruited. Number of genes from each tissue is shown below the tissue name with the percentage of total genes recruited in parentheses.

#### Comparisons of gene expression profiles between rumen and the first-chamber stomach of camels and cetaceans

Among these FC-specific genes, the three FC stomachs shared 18 genes which are co-expressed in the esophagus in all species (**Supplementary Table 5**). The 18 genes were significantly enriched in keratinocyte differentiation (**Supplementary Table 6**, Fisher’s exact test, adjusted *P* value = 9.85×10^−3^). This is consistent with the fact that the FC stomachs all share a basic stratified squamous epithelium with the esophagus^20–22^, which is markedly different from other stomach chambers (e.g. the abomasum of the ruminants, the third-chamber stomachs of camels and cetaceans). Notably, *PAX9*^23^, a known key transcription factor during esophagus differentiation, is highly expressed in all three FC stomachs and may play a role in the origin of the FC stomachs from their anatomic origin (**Supplementary Table 5**). Our results therefore indicate that the FC stomachs in Cetartiodactyla share a common developmental origin from the esophagus, and that changes in epidermis development may be an ancestral feature in this proto-rumen.

Despite the shared features of epithelial histology found in all Cetartiodactyla FC stomachs, the rumen also has a series of unique structural and functional innovations. Among the 655 rumen specifically expressed genes, we identified 448 up-regulated and 79 down-regulated genes when compared to the FC stomachs of camels (Fig. 2b; **Supplementary Table 7**), and 563 up-regulated and 29 down-regulated genes when compared to the FC stomachs of cetaceans (Fig. 2b; **Supplementary Table 8; Supplementary Note**). Among these, the majority (427, 65.2%) are up-regulated in rumen relative to both the FC stomach of camels and cetaceans (Fig. 2b; **Supplementary Table 9**). These exclusively rumen-specific (i.e., not specifically expressed in other FC stomachs) genes are significantly associated with the synthesis and degradation of ketone bodies (Fisher’s exact test, adjusted *P* value = 1.21×10^−3^) (Fig. 2c; **Supplementary Table 10**). Unlike monogastric animals, in which ketogenesis mainly occurs in the liver and the intestinal tract^24,25^, the rumen is the main site of ketogenesis in adult ruminants, and the occurrence of ketogenesis is regarded as a diagnostic feature of rumen maturity^26^. In addition to ketogenesis genes, seven genes from the KEGG pathway *Staphylococcus aureus* infection were also highly expressed in the rumen compared to the FC stomachs of camels and cetaceans (Fisher’s exact test for KEGG pathway enrichment, adjusted *P* value = 1.35×10^−2^) (**Supplementary Table 10**). These results indicate that improved ketone body metabolism and microbial regulation were important features in the evolution of the rumen from a proto-rumen origin shared with other Cetartiodactyls.

#### Gene recruitment by the rumen

Among the 655 rumen specifically expressed genes, the rumen co-expressed 96 (14.7%) genes with the esophagus(Fig. 2d; **Supplementary Table 2**). The 96 genes were enriched in the cornified envelope (adjusted *P* = 4.11×10^−14^) and epidermal cell differentiation processes (adjusted *P* = 3.77×10^−25^) (**Supplementary Table 11**). Meanwhile, we also found that the rumen recruited genes from a range of other tissues and biological pathways (Fig. 2d), e.g. keratinocyte differentiation (**Supplementary Table 12**, 88 genes co-expressed with keratinization-associated tissues), urea cycle (**Supplementary Table 13**, 24 genes co-expressed with liver), monocarboxylic acid transport (**Supplementary Table 14**, 61 genes co-expressed with intestine), skeletal muscle contraction (**Supplementary Table 15**, 23 genes co-expressed with muscle), urea transport (**Supplementary Table 16**, 19 genes co-expressed with kidney) and saliva secretion (**Supplementary Table 17**, 10 genes co-expressed with salivary gland). These pathways are all strongly associated with known rumen functions. For instance, enhanced urea recycling is an important characteristic of the rumen leading to increased nitrogen utilization for ruminants^27^. Collectively, these results suggest that the rumen—in addition to up-regulating genes expressed in the esophagus—recruited genes from different tissues to evolve its unique structure and complex functions.

#### Identification of genes functioning in early rumen development

The above rumen specifically expressed genes are identified in postnatal rumen, but the development of the rumen structure mainly occurs during early embryo stages^28,29^. In order to identify genes functioning in this critical stage, we performed five RNA sequencing from the ruminal and esophageal epithelium cells of four 60 days’ sheep embryos, the stage at which the ruminal epithelium starts to differentiate^28,29^ (**Supplementary Table 1**). We identified 285 rumen up-regulated differentially expressed genes (DEGs) compared to the esophagus (**Supplementary Table 18**). These are enriched in cell-cell junction (adjusted *P* value = 8.33×10^−3^) and desmosome organization (adjusted *P* value = 1.47×10^−3^) (**Supplementary Table 19**). We also found 1,840 rumen down-regulated DEGs which are enriched in anatomical structure morphogenesis (adjusted *P* value = 1.39×10^−15^) (**Supplementary Table 18, 20**). These results indicate that the specific epithelial histology of the rumen wall constitutes the most significant developmental genetic reprogramming as the organ forms and grows in the embryo. After filtering redundancy, we combined the 655 rumen specifically expressed genes with the 285 rumen up-regulated DEGs compared to the esophagus at the key development stage and eventually obtain 846 rumen key genes which we consider crucial for rumen development and evolution.

### Evolutionary analyses on the rumen key genes

Based on the data from ruminant comparative genomics^30^, we employed evolutionary genomic analyses on the 846 rumen key genes in the evolutionary context of 51 ruminants and 12 other mammals, by identifying ruminant-specific conserved nonexonic elements (RSCNEs) (≥ 20 bp), newly evolved genes and positively selected genes (PSGs) to systematically investigate the genetic changes associated with these rumen key genes. In the common ancestor of Ruminantia, we identified 657 genes with RSCNEs (**Supplementary Table 21**), two newly evolved genes and 28 PSGs (**Supplementary Table 22**) among the 846 rumen key genes. They are mainly involved in keratin filament binding, serine-type peptidase activity, ketone body metabolism and detection of bacterium.

#### Improved ketone body synthesis in rumen

In the pathway of synthesis and degradation of ketone bodies, *HMGCS2* and *SLC16A1* were under positive selection in the common ancestor of ruminants (Fig. 2c, 3a; **Supplementary Table 9, 10, 22**), and had ruminant-specific mutations when compared to non-ruminant mammals (Fig. 3b). Of the five ruminant-specific amino acid changes in the HMGCS2 protein, four are located in the HMG-CoA synthase domain (PF01154) (Fig. 3b). To further examine the effects of these mutations on the enzyme structure, we conducted three-dimensional (3D) structure simulations, and found that mutations in HMG-CoA synthase domain could induce a change of the protein 3D structure when compared to the human HMGCS2 protein (Fig. 3c). We also noted that the *SLC16A1* gene, which participates in the transportation of ketone bodies into the blood^24^, exhibited seven ruminant-specific mutations, six of which are located in the MFS_1 domain (PF07690), resulting in a domain structure change as revealed by protein structure homology-modeling (**Fig. S2, S3**). We therefore hypothesized that the changes in *HMGCS2* and *SLC16A1* may result in a more efficient ketone body metabolism in ruminants. This is supported by *HMGCS2* being the key rate-limiting enzyme in the ketogenesis pathway^24^. To explore the functional relevance of these mutations, we synthesized sheep and human *HMGCS2* orthologs *in vitro* and tested their enzyme synthetic activities by measuring the activities in a reconstituted system consisting of the enzyme and substrate (**Supplementary Note**). The sheep HMGCS2 (S) protein variant exhibites significantly higher metabolic efficiency than human proteins (H) (~2-fold increase, t-test, *P* < 0.001) (Fig. 3d). The enzyme activity of human HMGCS2 containing the five ruminant-specific amino acids replacements (H-5R) is also significantly higher than the regular human protein (~1.5-fold increase, *P* <0.01), while sheep HMGCS2 with the corresponding five human amino acid replacements (S-5H) exhibites significantly lower enzymatic activities than the sheep protein (~2-fold decrease, *P* < 0.001) (Fig. 3d). These results confirm that ruminants have evolved a more efficient ketogenesis than that of other mammals.

**Fig. 3.**
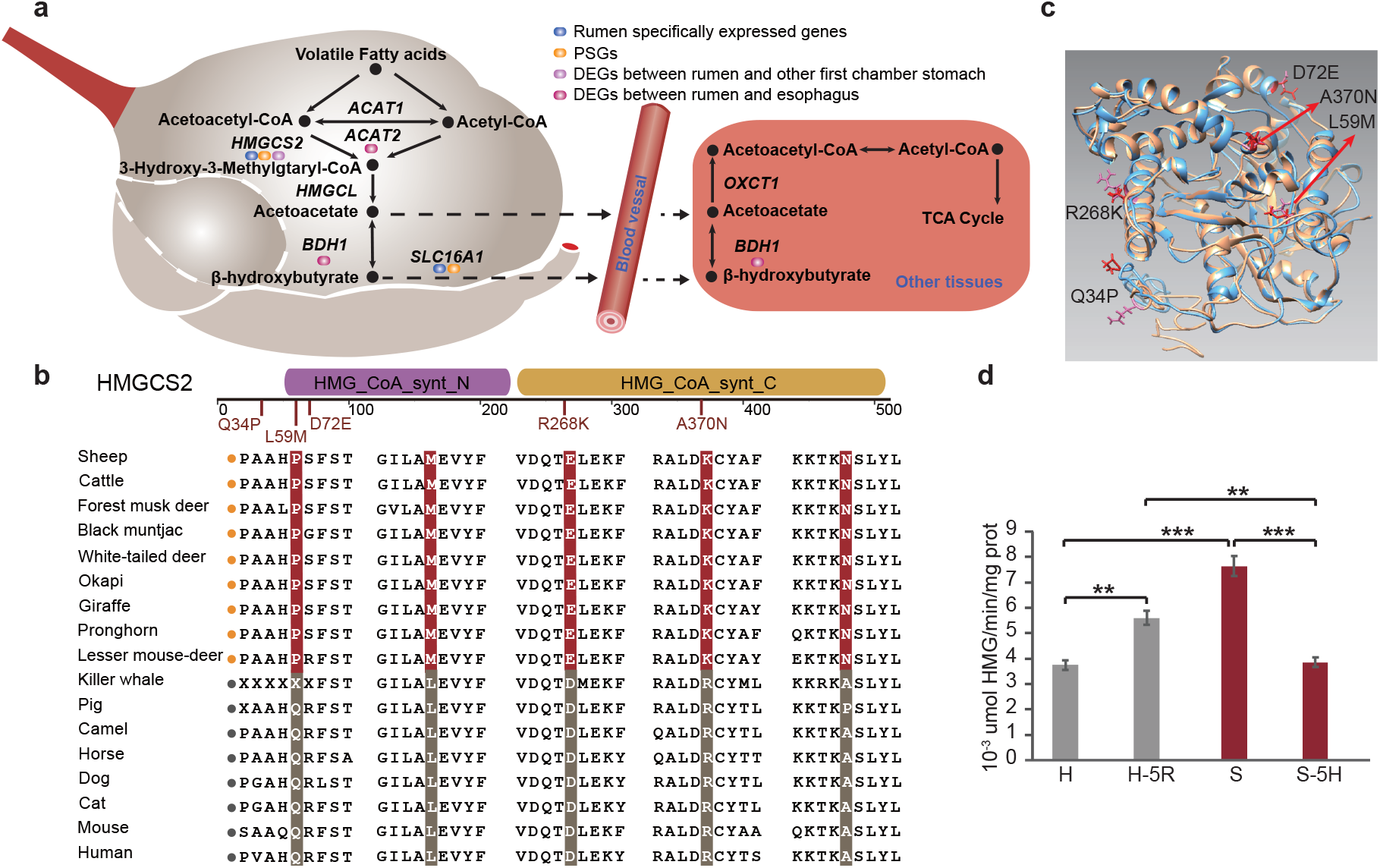
Genetic changes in the rumen ketone body metabolism genes and pathways. **a**, Genes annotated in the ketone body metabolism are labeled with different color to indicate rumen specifically expressed genes (blue), positively selected genes in ruminant (orange) and differentially expressed genes between rumen and other FC stomachs (purple). The solid arrows represent ketone body metabolism pathways. The dashed arrows indicate the process of material transport from rumen to other tissues. **b**, Top panels: Structural domains of the HMGCS2 protein and the location of the ruminant specific mutations. Lower panel: Peptide sequence alignment of HMGCS2. The species is followed a yellow circle belonging to the ruminant. The red highlighting indicates ruminant-specific amino acid mutations. **c**, Predicted tertiary structures of the HMGCS2 of ruminant (blue) and other mammals (orange), respectively. **d**, Enzyme activities of HMGCS2 compared with those of sheep and human in vitro. H: human, H-5R: human HMGCS2 with five ruminant aa replacements, S: sheep, S-5H: sheep HMGCS2 with five human aa replacements. ** *p* value < 0.01, *** *p* value< 0.001 calculated from the t test. Data are shown as mean±s.d.

#### Immune system and microbial regulation

We identified one PSG (*NOD2*) (**Supplementary Table 22**) and two newly evolved genes (*DEFB1* and *LYZ1*) in the rumen key gene list that are involved in immune functions. Among these, our transcriptomic data show that *NOD2* was co-expressed with the macrophage cells, and highly expressed in the rumen compared to both the FC stomachs of camels and cetaceans (**Supplementary Table 2, 9**). We detected 11 ruminant-unique amino acid changes in NOD2, resulting in domain structure changes as revealed by protein structure homology-modeling (**Fig. S4, S5**). This gene functions in the upstream part of IL17 signaling pathway, activating the Th17 cells to produce IL17F as part of the gastrointestinal immune system^31^ (Fig. 4a). The IL17 signaling pathway protects the host against extracellular pathogens via activating downstream pathways to induce the expression of antimicrobial peptides^32^.

**Fig. 4.**
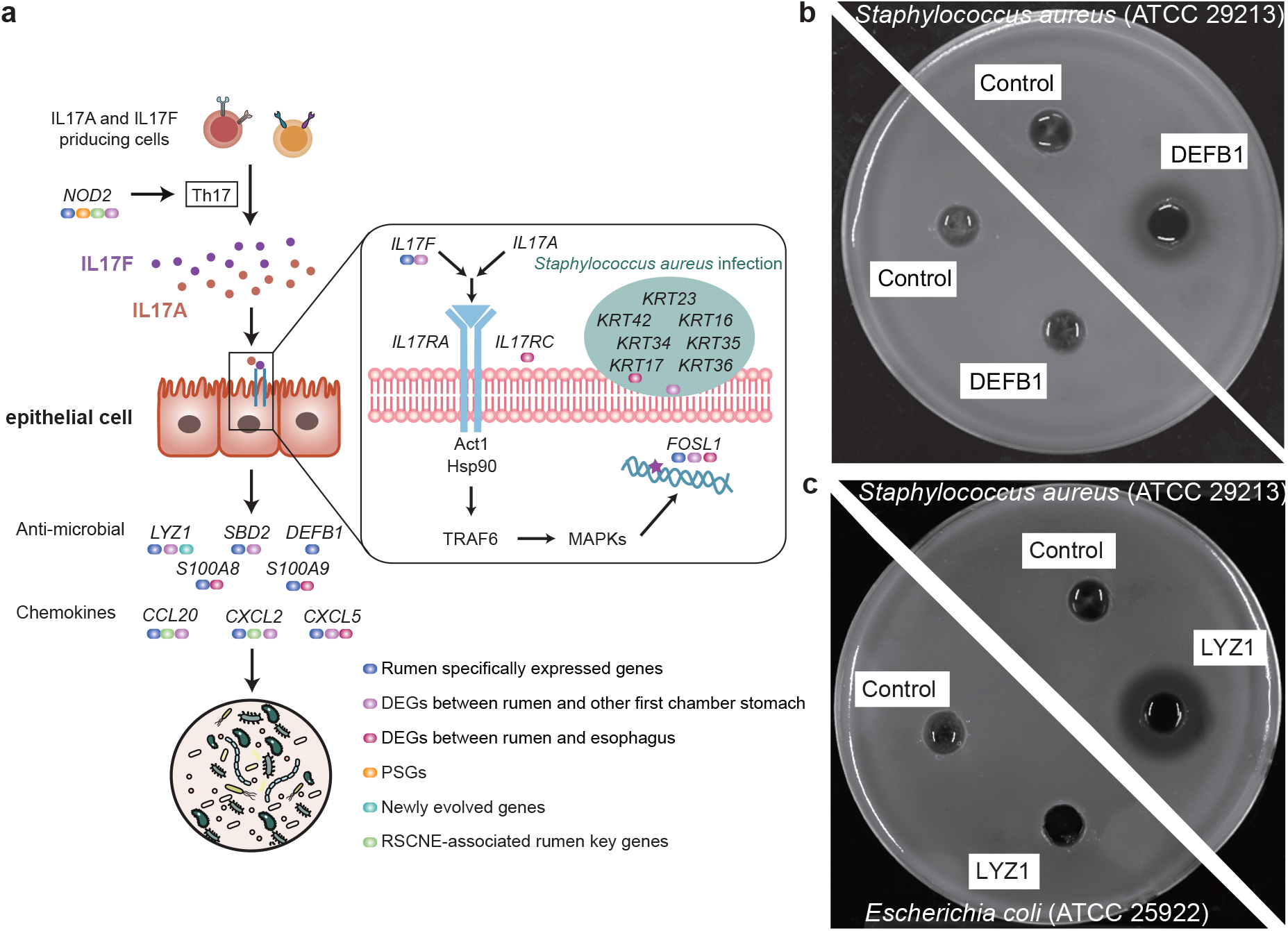
Microbial management of the rumen. **a**, Rumen specifically expressed genes (blue), differentially expressed genes between rumen and other FC stomachs (purple), positively selected genes in ruminant (orange), differentially expressed genes between rumen and esophagus (red), newly evolved genes (cyan) and RSCNE-associated rumen key genes (green) involved in IL17 signaling pathway and *Staphylococcus aureus* infection. The antibacterial ability of **(b)**, DEFB1 and **(c)**, LYZ1. Inhibition zone assays on agarose plates with *Escherichia coli* (ATCC 25922) and *Staphylococcus aureus* (ATCC 29213).

Among the newly evolved genes in the ancestor of ruminants, we identified a rumen key gene, *DEFB1*, which belongs to the beta-defensin family that have important roles as antimicrobial peptides in the resistance of epithelial surfaces to microbial colonization (**Supplementary Table 2**). In addition, we identified one newly evolved rumen key gene *LYZ1* in the lysozyme *c* family (**Supplementary Table 2**), which may protect the rumen epithelium from the activity of pathogenic bacteria^18^. We predicted that the LYZ1 contains a ruminant-specific 20 amino-acid-chain that encodes a probable transmembrane anchor (**Fig. S6, S7**), suggesting that the *LYZ1* gene encodes a secreted membrane-anchored protein, which may act on the rumen environment.

To validate the functions of these two newly evolved genes, we synthesized DEFB1 and LYZ1 *in vitro* and tested their antibacterial ability by performing an inhibition zone assay on agarose plates with *Escherichia coli* (American Type Culture Collection, ATCC 25922) and *Staphylococcus aureus* (ATCC 29213) as representative of Gram-negative and -positive bacteria (**Supplementary Note**). The DEFB1 (Fig. 4b) and LYZ1 (Fig. 4c) protein both showed antibacterial activity to *S. aureus*, but not *E. coli*. This characteristic of selective inhibition of Gram-positive bacteria is similar to that of monensin, which is commonly used as an antibiotic drug that regulates the microbiome and increases ruminant feed conversion efficiency^33,34^. Taken together, these results highlight that several important antibacterial functions are uniquely evolved in the rumen relative to other similar organs, and that some of these may work by specifically managing the microbiome composition.

#### New regulatory elements related to rumen epithelium absorbtion function

We searched among 221,166 RSCNEs to identify candidate regulatory regions in the vicinity of rumen key genes. We found that 657 of the 846 rumen key genes have nearby RSCNEs (**Supplementary Table 21**). To assess the regulatory role of these RSCNEs in the recruitment of increased gene expression in the rumen, we performed eight ATAC-seq libraries of the ruminal and esophageal epithelium cells from four 60 days’ sheep embryos (**Supplementary Table 23; Supplementary Note**). Our analysis indicates that 243 rumen key genes have nearby RSCNEs overlapping with identified open accessible peaks (**Supplementary Table 24**), and these genes are enriched in epidermal cell differentiation (adjusted *P* value = 4.82×10^−19^) (**Supplementary Table 25**). In the comparison of ATAC-seq between the rumen and esophagus, we identified 3,904 rumen-specific and 5,531 esophagus-specific open differentially accessible peaks (DAPs) (**Fig. S8; Supplementary Table 26**). Interestingly, we found 267 and 478 RSCNEs (≥ 20 bp) overlapping with rumen-specific and esophagus-specific DAPs, which is highly statistically signficant (Fisher’s exact test, both *P* value = 0.00). Rumen-specific DAP-associated RSCNEs are physically near 22 rumen key genes (**Supplementary Table 27**). Among these genes, *CRNN* is one of the genes in the epidermal differentiation complex (EDC) locus, which is essential for the cornified cell envelope in rumen^15^, and is implicated in several epithelial malignancies in human^35^. A rumen-specific DAP-associated RSCNE with six ruminant-specific mutations was found at the 5’ upstream of *CRNN* of ruminants, which might play a role in regulating its expression in rumen. Concordantly, *DMRT2* is a key transcriptional factor in the dermomyotome organization and *DMRT2*-deficient mice have epithelial morphology abnormalities^36^. We observed that *DMRT2* has five rumen-specific DAP-associated RSCNEs in its 3’ downstream region, potentially causing high *DMRT2* expression in rumen.

Interestingly, *WDR66* is not only highly expressed in the rumen compared with both the FC stomachs of camels and cetaceans but also under positive selection in the common ancestor of Ruminantia (Fig. 5a; **Supplementary Table 9, 22**). It regulates the expression of occludin, which tightens the intercellular space and enables epithelial permeability^37^. We observed 10 ruminant-specific non-synonymous mutations and one rumen-specific DAP-associated RSCNE in the intronic region of *WDR66* (Fig. 5b; **Fig. S9; Supplementary Table 27**). In order to assess the regulatory activity of this particular RSCNE, we cloned it into a luciferase reporter vector (pGL3-Promoter) and transfected it into both sheep and goat fibroblasts *in vitro*. The RSCNE showed significantly higher luciferase transcriptional activation compared to the pGL3-Promoter control (t-test, *P* < 0.05) (Fig. 5c), confirming that it acts as an enhancer. Therefore, these DAP-associated RSCNEs might plausibly have exerted novel *cis*-regulation of the rumen key genes, thus providing a mechanistic explanation of how the rumen might have recruited these genes from other tissues. Hence, we propose a central role of such regulatory elements in the development and evolution of rumen structure and function.

**Fig. 5.**
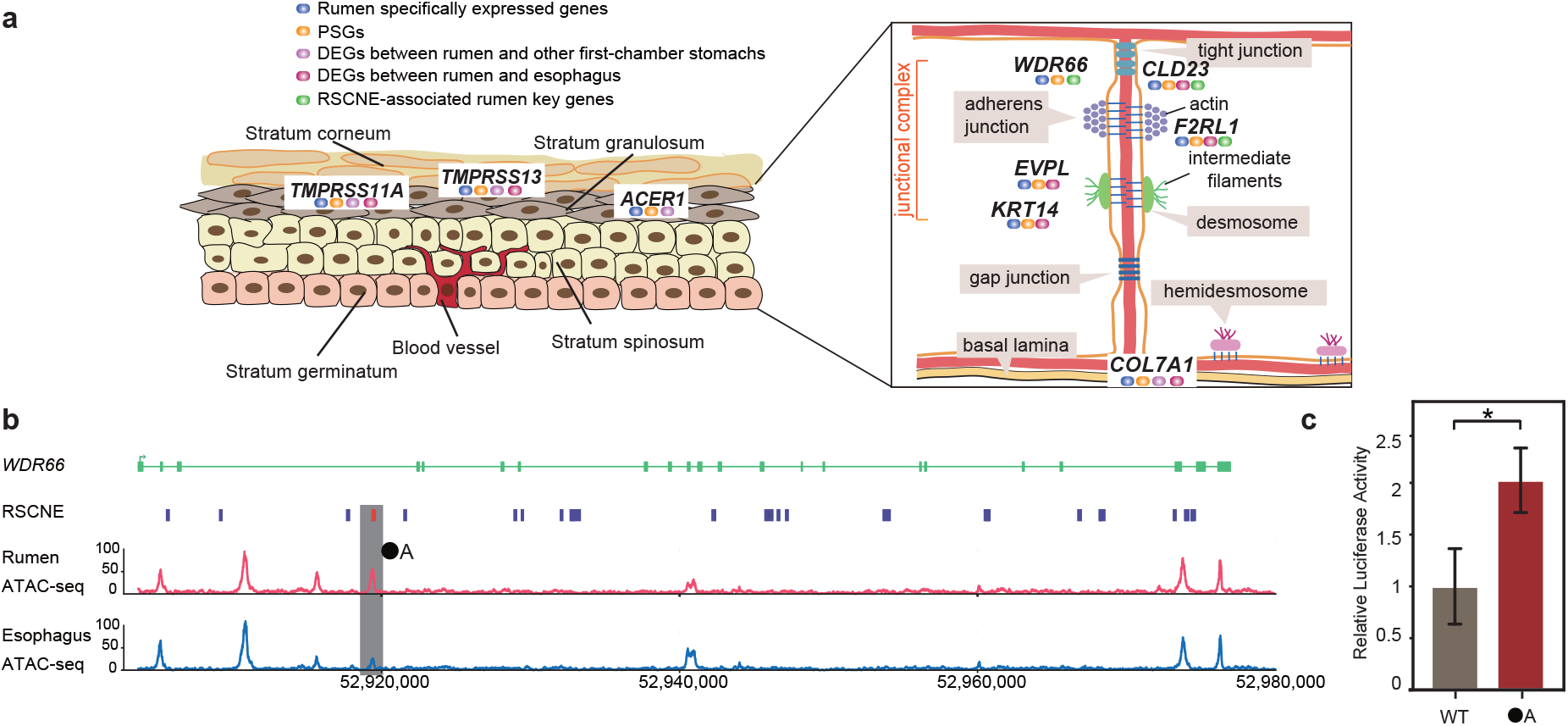
Genetic changes related to rumen epithelium transportation and absorption. **a**, Diagram of rumen epithelial cell proteins involved in epithelium permeability identified in the common ancestor of the ruminants. Rumen specifically expressed genes (blue), positively selected genes in ruminant (orange), differentially expressed genes between rumen and other FC stomachs (purple), differentially expressed genes between rumen and esophagus (red), and RSCNE-associated rumen key genes (green). Note the junction structure (desmosome) between keratinocytes of the ruminal epithelium has been degraded, instead the enlarged intercellular space with copious blood supply enables metabolites absorption in the ruminal epithelium^47^. **b**, Gene structure of *WDR66* based on the NCBI Oar_v4.0 annotation shown above. Green boxes represent exons. Purple bars indicate ruminant-specific conserved non-exonic elements (RSCNEs). Red and blue bars indicate ATAC-seq peaks of the ruminal and esophageal epithelium cell, respectively. The grey rectangle box is the overlapping element of RSCNE and ATAC-seq which is located in the intron region. **c**, The luciferase activity of the pGL3-Promoter (WT) and the pGL3-Promoter with the RSCNE (●A). * *p* value < 0.05 calculated from the t test. Data are shown as mean ± s.d.

#### Positively selected genes involved in rumen epithelium absorption

We observed that eight rumen key genes involved in the cell junction biological process (*WDR66*, *COL7A1*, *EVPL*, *KRT14*, *CLDN23*, *F2RL1*, *TMPRSS13* and *TMPRSS11A*) were under positive selection in ruminants (Fig. 5a; **Fig. S9-S16**; **Supplementary Table 22**). Non-synonymous changes in these genes may result in the change of cell junctions, which may break the epithelium barrier and increase the epithelium absorption properties^38–42^. *COL7A1* is highly expressed in the rumen of fetal sheep, but not in the esophagus (**Supplementary Table 18**). We detected 17 unique amino acid (aa) changes in COL7A1 in ruminants (**Fig. S10**). COL7A1 is an anchoring fibril between the external epithelia and the underlying basal lamina^39^. Amino acid mutations in this gene are associated with epidermolysis bullosa, a condition in which tissue fluid diffuses through the intercellular space into the epidermis^39^. In addition, *TMPRSS13*, a membrane-anchored serine protease gene^41^, is highly expressed in rumen compared to esophagus (**Supplementary Table 18**). Interestingly, we identified five ruminant-specific aa changes in TMPRSS13, four of which are located in the trypsin-like serine protease domain (**Fig. S15**). It is reported that the deficiency of *TMPRSS13* in mice impairs stratum corneum formation and epidermal barrier acquisition, accompanied by trans-epidermal fluid loss^41^. In normal epithelium cells (e.g., epithelium cells of skin), the epithelium barrier is produced by strong intracellular protein filaments crossing the cytoplasm and attaching to specialized junctions, which in turn ties the surfaces of adjacent cells either to each other or to the underlying basal lamina^43^ (Fig. 5a). Given that the epithelium transportation and absorption functions are affected by the epithelium barrier, mutations in these cell junction-related genes may be related to metabolite uptaking function of the rumen.

## Discussion

Our large quantity of transcriptomic data in adults and an early embryo rumen development stage provide a detailed comparative insight into the distinct gene expression profile of the rumen. Although there has been no consensus about the evolutionary relationship between the FC stomachs of camels, peccaries, cetaceans and ruminants^21,44^, it is unlikely that the multi-chambered stomach evolved independently four times in Cetartiodactyla exclusively. Therefore, the most parsimonious explanation is that they may have a single evolutionary origin, followed by specialization in the different lineages of the Cetartiodactyla due to their specific diets and niches. For instance, the FC stomachs of camels have evolved the ability to store water^21,45^, the FC stomachs of cetaceans has the capacity to mechanically grind food^11^, and the rumen provides efficient fermentation and metabolism of plant material. The gene expression profiles of the FC stomachs in ruminants, camels and cetaceans show that they are all highly similar to the esophagus, suggesting these organs share an anatomical origin from the esophagus (Fig. 2a; **Fig. S1**).

Based on our comparative genomic and functional data, we outline the genetic mechanisms underlying the origin, development and evolution of the rumen from the ancestral esophagus tissue. These genetic innovations are mainly related to epithelium absorption, ketone body metabolism and microbial regulation. Among the 846 rumen key genes (**Supplementary Table 2, 18**), we found that 657 (77.7%) genes have nearby RSCNEs (**Supplementary Table 21**), 28 genes are under positive selection (**Supplementary Table 22**) and two genes newly evolved in the common ancestor of ruminants, suggesting these three types of genetic reprogramming all contributed to the structural and functional evolution of rumen. Notably, the majority of rumen key genes have RSCNEs nearby and our ATAC-seq validated that 243 rumen key genes had nearby RSCNEs overlapping with highly accessible chromatin (**Supplementary Table 24**), suggesting the RSCNEs as regulatory elements may play a crucial role in rumen gene recruitment. The highly significant association between RSCNEs, rumen key genes and open accessible peaks is a strong indication of this, although there were also many RSCNEs that did not overlap with open accessible peaks in our ATAC-seq analysis. While this suggests that RSCNEs play other roles besides being regulatory elements, it is also possible that some were false negatives due to the limitations of development stages sampled in this study, which might have omitted some associations between rumen key genes and regulatory RSCNEs. Hence, a denser sampling of different developmental time points might expand the rumen key gene list and reveal novel regulatory roles of RSCNEs. Nevertheless, our study has revealed the important genetic mechanisms underlying the key evolutionary innovations of the rumen. The identified rumen key genes and their specific mutations provide a starting point for future studies of rumen development, and for understanding the interactions between rumen and microbiota. This will be key to further improvement of ruminant livestock, e.g. by providing a framework for manipulating the rumen fermentation process.

## Data availability

The raw reads for all RNA-seq data, the ATAC-seq data from the rumen and the esophagus have been deposited at the Sequence Read Archive (SRA) under project number PRJNA485657.

## Acknowledgments

This project was supported by the National Natural Science Foundation of China (31822052, 31572381), the National Thousand Youth Talents Plan to Y.J., National Natural Science Foundation of China (31660644) to S.H, National Natural Science Foundation of China (41422604) to S.L. We thank the members of the FANNG project for sharing their transcriptome data. We thank Yongchuan Li, Zhengzhi Wei, Zixin Yang, and Haiyu Gao from Institute of Deep-sea Science and Engineering, Chinese Academy of Sciences, for helping to collect samples from the porpoise and whale. We thank High-Performance Computing (HPC) of Northwest A&F University (NWAFU) for providing computing resources.

## Author contributions

Y. J. and W.W. conceived the project and designed the research. X.P., Y.C., N.W., C. Z., and X.H. performed the majority of analysis with contributions from K.W., L.C., Z.L., Z.Z., B.W., S.H.; Q.Q., S.M., X.L., W.F., L.L., Y.L., W.S., W.L., T.Z., J.H., M.L., S.L., S.H., M.L., C.L., and Y.C. prepared the sheep, camels and cetaceans samples for transcriptomics and rumen and esophagus epithelium cells for ATAC-seq. H.L. performed the luciferase reporter assay. X.C., Y.Y. and Z.H. performed the inhibition zone assay and the enzyme synthetic activities assay. X.P., Z.L. and Y.C. drafted the manuscript with input from all authors, whereas Y.J., W.W., R.H., B.P.D., G.Z., X.W. and Y.W. revised the manuscript.

## Competing interests

Two provisional Chinese patent applications on potential application in the antimicrobial and antibiotic substitute by way of the *DEFB1* gene and *LYZ1* gene have been filed by Northwest A&F University (application number 202010100677.8 and 202010097562.8), where Y.J., X.P., X.C, and W.W. are listed as inventors. The authors declare no competing interests.

