## Supplementary Information for "Tracing the origin of a new organ by inferring the genetic basis of rumen evolution"

#### Table of contents

**Supplementary Notes**

**Supplementary Figures**

**Supplementary Tables**

**References**

### Supplementary Notes

#### Part 1. Phylogeny relationship and sample collection

##### 1.1 Construction of the phylogenetic tree

To present the evolutionary panorama of the multi-chambered stomach evolution in Cetartiodactyla, we constructed a phylogenetic tree using four-fold degenerate sites using single-copy orthologous genes from nine species (human, horse, camel, pig, hippo, killer whale, lesser mouse-deer, roe deer, and sheep) as representatives of the major taxonomy. A final maximum likelihood tree was generated using IQ-TREE multicore (version 1.6.6.a)[1] with the parameters "-bb 1000 -m TEST -o Human" (**Fig. S17**). Note that one of the families in the Suina, the Tayassuidae, has one stomach with three chambers. Although no genomes of species in the Tayassuidae are available, we chose the peccary as the representative of Tayassuidae and present the position of the peccary according to the results of mitochondrial genomes[2]. The newick format of tree result is as follows: (((((LesserMouse-deer:0.1383065387,(Roedeer:0.0549981637,Sheep:0.0511103239)100:0.0574393138)100:0.0488086408,(Killerwhale:0.0631824323,Hippo:0.0758665489)100:0.0082276945)100:0.0179186732,Pig:0.1394609521)100:0.0101113682,Camel:0.1196798498)100:0.0411710418,Horse:0.1192130573,Human:0.1795944986);

##### 1.2 RNA-seq analysis

###### 1.2.1 Collection and sources of samples for RNA-seq

We collected and sequenced a total of 323 tissue samples from 11 Texel ♂ × Kazakh ♀ hybrid sheep (*Ovis aries*) in Yili city (Xinjiang, China) (210 sequenced and 123 used in another paper at our laboratory)[3]. Other sequencing data was downloaded from 83 tissue samples from 4 Texel sheep included in the NCBI BioProject PRJEB6169[4] and 426 tissue and cell samples from 33 Texel ♂ × Scottish Blackface♀ hybrid sheep from ENA (accession number PRJEB19199)[5] collected in independent studies. These 832 samples covered 50 tissues of all major organs and tissues and allowed the classification of the rumen-specificity of the expression of detected genes.

We also collected 20 tissue samples from one roe deer (*Capreolus pygargus*)[3], which was used as the representative of the Cervidae.

Thirty-two samples from seven Bactrian camel (*Camelus bactrianus*) and eight samples from two cetaceans (*Neophocaena asiaeorientalis* and *Balaenoptera brydei*) were used as outgroups for comparative transcriptome analysis of the rumen. Information about all tissue samples is provided in **Supplementary Table 1**. Samples of camels are from Hualing Animal Husbandry Base in Urumqi (Xinjiang, China). Samples of cetaceans are from two naturally deceased individuals, provided by Institute of Deep-sea Science and Engineering, Chinese Academy of Sciences. All other animals were slaughtered under the guidelines of Northwest A&F University Animal Care Committee. All samples were frozen in liquid nitrogen until utilization. All tissues rinsed with PBS were dissected for RNA sequencing.

##### 1.2.2 RNA isolation, library construction, and sequencing

In all tissue samples collected for this study, total RNA was isolated from a frozen sample according to the Trizol protocol (Invitrogen, USA), using 1.5 μg RNA per sample as the input material for sample preparation. Sequencing libraries were generated using a NEBNext® Ultra RNA Library Prep Kit for Illumina® (NEB, USA) according to the manufacturer's recommendations, and index codes were added to attribute sequences to samples. Briefly, mRNA was purified from total RNA using poly-T oligo-attached magnetic beads and fragmented using divalent cations at elevated temperature in NEB Next First-Strand Synthesis Reaction Buffer (5X). First-strand cDNA was synthesized using random hexamer primers and M-MuLV Reverse Transcriptase (RNase H). Second-strand cDNA was subsequently synthesized using DNA Polymerase I and RNase H. Remaining overhangs were converted into blunt ends by exonuclease/polymerase activity. After adenylation of 3' ends of DNA fragments, NEB Next Adaptors with hairpin loop structures were ligated to prepare for hybridization. To select cDNA fragments with appropriate lengths, the library fragments were purified with an AMPure XP system (Beckman Coulter, Beverly,

USA). Then 3 μl of USER Enzyme buffer (NEB, USA) was incubated with size-selected, adaptor-ligated cDNA at 37 °C for 15 min followed by 5 min at 95 °C before PCR amplification, using Phusion High-Fidelity DNA polymerase, Universal PCR primers, and Index (X) Primer. Finally, PCR products were purified using the AMPure XP system, and library quality was assessed using an Agilent Bioanalyzer 2100 system. The index-coded samples were clustered with a cBot Cluster Generation System using a HiSeq 4000 PE Cluster Kit (Illumina) according to the manufacturer's instructions. After cluster generation, the library preparations were sequenced on an Illumina Hiseq X Ten platform, and 150 bp paired-end reads were generated. All these sequencing procedures were performed by Novogene Technology Co., Ltd., Beijing, China.

**1.2.3 Data quality control and quantification processing**

We obtained high-quality reads by removing adaptor sequences and filtering low-quality reads from raw reads using Trimmomatic (version 0.36)[6] with the following parameters: LEADING:3 TRAILING:3 SLIDINGWINDOW:4:15 MINLEN:40. High-quality reads from the 832 sheep samples were all aligned to the NCBI assembly Oar_v4.0 reference sheep genome[4]. High-quality reads from the 32 samples of Bactrian camel were aligned to the NCBI assembly Ca_bactrianus_MBC_1.0 reference genome[7]. High-quality reads from the 8 samples of cetaceans were aligned to the NCBI assembly Neophocaena_asiaeorientalis_v1 reference genome[8]. For this, we used STAR (Version 2.5.1)[9] with the following parameters: outFilterMultimapNmax 1, outFilterIntronMotifs RemoveNoncanonical Unannotated, outFilterMismatchNmax 10, outSAMstrandField intronMotif, outSJfilterReads Unique, outSAMtype BAM Unsorted, outReadsUnmapped Fastx, and outFileNamePrefix. The unmapped reads were extracted by SAMtools (Version 1.3)[10] for further mapping by HISAT2 (Version 2.0.3-beta)[11]. We assembled transcripts including novel splice variants by StringTie (Version1.3.4)[12] and computed Fragments Per Kilobase per Million mapped reads (FPKM) values for the transcripts and genes in each sample using Ballgown (Version 2.2.0)[13]. Finally, we removed transcripts with FPKM lower than one in more than 90% of samples of each tissue.

##### 1.3 Identification of the ancestral organ of the first-chamber stomach

We generated tissue expression profiles with samples from sheep (n=832), roe deer (n=20), Bactrian camel (n=32), cetaceans (including Bryde's whale (n=2) and Indo-Pacific Finless Porpoise (n=6)). FPKM values for all genes were used to calculate a correlation matrix based on Spearman's rank correlation coefficients for all pairwise combinations of sheep, roe deer, Bactrian camel, and cetaceans samples for further identification of the ancestral organ of the first-chamber stomach. The correlation matrix was used for unsupervised hierarchical clustering analyses of samples, and the results were visualized in a heatmap of all pairwise correlation coefficients between tissues (**Fig. 2A; Fig. S1**).

Based on the sheep tissue expression profile, we found that the rumen is more highly correlated with the esophagus, reticulum and omasum than with the stomach (Spearman correlation coefficients: 0.8996, 0.8982 and 0.9563, respectively). We excluded the reticulum and omasum as ancestral organs because they are both regarded as parts of the forestomach with the rumen. Moreover, lesser mouse-deer (*Tragulus javanicus*) (the so-called proto-ruminant family) has only a three-chamber stomach, that lacks an omasum[14]. Hence, we prefer that the reticulum and the omasum appeared later in the process of chamber-stomach evolution. We used the roe deer tissue expression profile to verify our conclusions regarding the rumen in Cervidae.

##### 1.4 Identification of first-chamber stomach specifically expressed genes in ruminants, camels and cetaceans

The maximum FPKM values for the genes in all samples of each tissue were used to estimate their expression levels. A cutoff value of one FPKM was used as the detection limit in rumen samples. We used the concept of genome N50 to define E (expression) 50, which is defined that the minimum number of tissues required to meet the half of the total log2-transformed expression (FPKM) in all tissues. This index is distributed in bimodal model in sheep, camels, and cetaceans, representing the specificity and broad-spectrum expression of genes. Firstly, Expectation Maximization (EM) Algorithm was used to perform parameter estimation on the bimodal model. Then, we selected the cutoff of E50 to meet the probability of the first type of error falling into the second peak which is less than 0.05. After using this cutoff to screen out the tissue-specific expression genes, we further screened out genes with the rumen expression rank which is less than or equal to E50 as the first-chamber stomach specifically expressed genes in ruminants, camels and cetaceans. Finally, all the FC stomach specifically expressed genes were verified by t-test to ensure that their expression was significantly higher than other tissues.

###### 1.4.1 Gene classification of FC stomach specifically expressed genes

Among the FC stomach specifically expressed genes, the tissues in which they had the second-highest expression level (second only to the rumen) were regarded as the source organs/tissues.

#### Part 2. ATAC-seq library preparation and sequencing

##### 2.1 Collection and sources of the rumen and the esophagus cell samples

We collected a total of eight samples of the rumen and esophagus epithelium tissues from four 60-day Hu sheep embryo (E60) from XiLaiYuan ecological agriculture co. LTD in Taizhou city (Jiangsu, China). All samples rinsed with PBS and were soaked in cold 1 x PBS added with penicillin-streptomycin. All animals were slaughtered under the guidelines of Northwest A&F University Animal Care Committee.

###### 2.1.1 Isolation of ruminal epithelial cells.

A piece of ruminal epithelial tissue was removed from PBS buffer (pH 7.4), placed on a watch glass, and brushed with sterile D-Hanks in all directions. The clipped tissue (approximately 500 mg) was placed in a small beaker and rinsed 2-3 times with a D-Hanks (pH 7.4) solution (4 times antibody, pre-warmed in a 37 °C water bath). Next, 0.25% trypsin (15 ml) was pre-warmed in a 37 °C water bath and added to a conical flask with the rumen epithelium sample, which was then digested in a 37 °C water bath for 30 min while shaking well every 5 min. The ruminal epithelial tissue was removed and the trypsin digestion solution was discarded. This step was repeated three times until the epithelium felt sticky. The treated epithelial tissue was then placed in a sterile beaker and rinsed three times with D-Hanks solution, and this step was repeated using a fresh beaker. Ten milliliters of trypsin were added, and the mixture was digested in a 37 °C water bath for 10-20 min until the cells detached. Cells from the first 3-4 detachments were not collected because these are generally necrotic or granular cells. Only the last two digested cell types (spinous and basal cells) were generally collected, after the cells in the digested sample were observed under a microscope. The cells were filtered through a cell strainer and added to a 10 ml centrifuge tube containing a drop of calf serum. The above digestion and collection steps were repeated 3 times. The digestate was collected following centrifugation at 1500 r/min for 5-10 min, and the supernatant was discarded. One milliliter of Dulbecco's Modified Eagle Medium (DMEM) solution was added to the precipitate, and the mixture shaken or blown to adjust the cell density to $10^6$ cells/ml. Trypan blue was added to verify that cell viability reached 95%.

The esophageal epithelial cells were obtained with the same pipeline with the ruminal epithelial cells as above.

**2.1.2 Preparation of nuclei**

To prepare nuclei, we spun 50,000 cells at 500 x*g* for 5 min and then washed the pellet using 50 μl of cold 1× PBS. The solution was then centrifuged at 500 x*g* for 5 min, and the cells were lysed using cold lysis buffer (10 mM Tris-HCI, pH 7.4, 10 mM NaCl, 3 mM $MgCl_2$ and 0.1 NP40). Immediately after lysis, the nuclei were spun at 500 x*g* for 10 min using a refrigerated centrifuge. To avoid losing cells during the nucleus preparation, we used a fixed angle centrifuge and carefully pipetted away from the pellet after centrifugation.

**2.1.3 Transposition and purification**

The pellet was immediately resuspended in transposase reaction mix (17.5 μl of

DEPC $H_2O$, 5μl of TTBL buffer, 2.5 μl of TTE mix buffer and all the nuclear DNA). The transposition reaction was carried out for 10 min at 55 ℃ in metal bath, and the sample was immediately purified using a Qiagen MinElute kit.

**2.1.4 Library construction**

PCR was performed to amplify the library for 14 cycles using the following PCR conditions: 72 ℃ for 3 min, 98 ℃ for 30 s, and thermocycling at 98 ℃ for 15 s, 60 ℃ for 30 s and 72 ℃ for 3 min.

**2.2 ATAC-seq analyses**

**2.2.1 Data quality control and short-read alignment**

Sequencing reads must undergo quality control and adapter trimming to optimize the alignment process. FastQC (version 0.11.5)[15] was used to assess overall quality. Reads were trimmed for quality as well as the presence of adapter sequences using the Trim Galore Wrapper script[16] with default parameters. Four rumen and four esophagus ATAC-seq reads of sheep were mapped to the sheep reference genome (NCBI assembly Oar_v4.0) using Bowtie2 (version 2.2.8)[17] with default parameters. Duplicated reads were removed using the default parameters in Picard (version 2.1.1)[18]. Reads mapping to mitochondrial DNA were excluded from the analysis together with low-quality reads (MAPQ < 20).

**2.2.2 Open accessible peak calling**

Accessible regions and peaks were identified using MACS[19] with parameters "-q 0.05 -shift 37 -extsize 73" for narrow peaks. The centers of identified peaks were used to define peak overlaps with genomic features according to the following criteria. If a center site was located in the promoter of a gene (2 kb upstream from the transcription start site (TSS)), or the gene body, the peaks would be assigned to that gene. Distal intergenic regions refer to regions > 3 kb from the TSS and > 1 kb from the transcription termination site (TTS).

We called 82,980, 106,412, 31,749, 128,111 peaks from four rumen ATAC-seq datasets and 121,564, 49,128, 153,495, 52,222 peaks from four esophagus ATAC-seq datasets, respectively. The plot and heatmap of regulatory region distribution (**Fig. S18-19**) were obtained using ChIPseeker[20]. The distribution of peaks throughout the sheep genome is shown in a pie chart (**Fig. S20**).

###### 2.2.3 Consensus peaks and differentially accessible peaks

Open accessible peaks were identified in four biological replicates of each tissue by using "bedtools intersect", and consensus peaks with openness values of each peak in each sample were built by merging these regions and calculated with R package "Diffbind" (version 2.10.0)[21]. Differentially accessible peaks were identified by R package "edgeR" (version 3.22.5)[22].

###### 2.2.4 Peak annotation

Peak annotation was performed using R packages "GenomicFeatures", "ChIPseeker", and "AnnotationHub".

##### 2.3 Corresponding RNA-seq analysis of ATAC-seq

###### 2.3.1 RNA-seq library preparation and sequencing

We prepared directional RNA-seq libraries from the cells of the same samples as used for ATAC-seq. Finally, we obtained three ruminal epithelium cell samples and two esophageal epithelium cell samples for RNA-seq. Each sample was added 1ml Trizol protocol (Invitrogen, USA), and frozen in -80 °C until utilization. The RNA-seq library preparation and sequencing procedures are as the same as above.

###### 2.3.2 Identification of differentially expressed genes between the rumen and the esophagus

We used R package "edgeR" (version 3.22.5)[22] to identify differentially expressed genes (DEGs) between the RNA-seq of the three ruminal epithelium cell samples and two esophageal epithelium cell samples. A regular threshold (false discovery rate value $< 0.05$ and $\log_2$(Fold Change) $> 2$ or $< -2$) was used to identify the DEGs.

###### 2.3.3 KEGG pathway and Gene Ontology (GO) enrichment and statistical analysis

All the GO enrichment analysis was applied by an in-house script. All the KEGG pathway enrichment analysis was applied by R package "clusterprofiler"[23]. All the fisher's exact test analysis was applied by python (version 3.6.0) module of *scipy.stats*.

**Part 3. Orthologous gene construction for differentially expressed analysis and evolution analysis**

**3.1 Orthologous gene construction for differentially expressed analysis**

**3.1.1 Construction of orthologous genes among sheep, camels and cetaceans**

To compared the differences among the rumen and the first-chamber stomach of camels and cetaceans, we firstly constructed the orthologous genes by using all-vs-all blastp[24] of the Bactrian camel proteins vs the sheep proteins and the Indo-Pacific Finless Porpoise proteins vs the sheep proteins. Finally, we identified 18,712 and 18,688 orthologous genes in Bactrian camel and cetaceans, respectively. Then we used rumen specifically expressed genes as the reference to match the corresponding genes in camels and cetaceans. If the rumen specifically expressed gene has no orthologous in camels or in cetaceans, then the expression of the gene will be filled with "0.001" in camels or in cetaceans.

**3.1.2 Differentially expressed genes among the rumen, the first-chamber stomach of camels and cetaceans**

A regular threshold (2-fold change or 0.5-fold change) was used to identify the DEGs between the rumen and the first-chamber stomach of camels or cetaceans.

**3.2 Evolution analysis**

**3.2.1 Identification of positively selected genes (PSGs)**

We cited the genomic features related to ruminant evolution results of Chen et al[25]. In this part, we introduced the methods briefly. Based the multiple genome alignments of all the 51 Ruminantia species and 12 outgroup mammals, the consensus coding sequences of each species were extracted based on the cattle annotation file (Bos_taurus.UMD3.1.87.gff3.check.gff). The codeml program in the PAML package (version 4.8)[26] with the free-ratio model (model=1) was run for each ortholog. A likelihood ratio test (LRT) was conducted to compare a model that allowed sites to be under positive selection on the foreground branch with the null model in which sites could evolve either neutrally and under purifying selection. The *p* values were computed based on *Chi*-square statistics, and genes with *p* value less than 0.05 were treated as candidates that underwent positive selection.

**3.2.2 Analysis of conserved non-exonic elements analysis**

This part is cited from Chen et al.,[25]. Twelve mammalian species were chosen as outgroups to identify the conserved elements of ruminants, namely, dolphin, minke whale, killer whale, sperm whale, camel, horse, rhinoceros, pig, cat, cheetah, dog and human. The non-conserved model was estimated by phyloFit (version 1.4, with the default parameters) in the PHAST package[27] with 4d sites in the ruminant alignments and the topology). Then, we ran phastCons[28] with the ruminant non-conserved model to estimate the ruminant conserved models (version 1.4, with parameters: phastCons --estimate-rho). Also, with phastCons, we predicted the highly conserved elements (with the parameters: phastCons --most-conserved --score) in ruminants and generated base wise conservation scores with the ruminant conserved and non-conserved models. Exon regions were excluded from the highly conserved elements to generate the conserved non-exonic elements (CNEs). We identified 2,346,281 CNEs with a total length of 241.8M longer than or equal to 20 bp. Next, we identified two types of ruminant-specific CNEs (RSCNEs). Type I RSCNEs were CNEs for which sequences were not found in either outgroup species. Type II RSCNEs were CNEs that had orthologous sequences in at least three outgroup species, but were only conserved in ruminants according to the phyloP (version 1.4, with the parameters: phyloP --method LRT --mode CONACC)[27] under the non-conserved model. For Type II RSCNEs, we separated the outgroups into three different sets (1) Cetacea: dolphin, minke whale, sperm whale and killer whale; (2) camel, horse, rhinoceros and pig; (3) Primates and Carnivora: cat, cheetah, dog, and human. Then, we filtered the results with phyloP ($p$ value $< 0.05$, false discovery rate (FDR) corrected) in all three sets of phyloP tests the ruminant with outgroups non-conserved model and. Additionally, to produce a higher quality set of RSCNEs, we removed elements of < 20 bp in length. Finally, we obtained 141,351 Type I RSCNEs and 79,815 Type II RSCNEs.

Genetic regions of RSCNEs were based on the coordinate of goat, then we turn the comparative genomes files in MAF format to chain format. Then we used liftOver tool in UCSC Genome Browser to correspond the RSCNEs to the coordinate of sheep.

**3.2.3 Identification of newly evolved genes in ruminants**

To identify newly evolved genes, we dated all cattle (UMD3.1) genes along the phylogenetic tree following the pipeline described by Zhang et al[29,30]. In brief, from genomes of goat (ASM1.1), black muntjac, pronghorn, giraffe, musk deer, killer whale (Oorc_1.1), pig (Sscrofa 11.1), horse (EquCab2.0), mouse (mm10) and human (hg38), we extracted their whole genome alignments. Based on the gene annotations of cattle, we investigated whether a reciprocal best-to-best match could be found for a coding sequence (CDS) between the two adjacent genomes. If 2/3 of the length of a CDS is covered with 50% identity, it suggests the presence of the CDS. By adopting a parsimony strategy, we classified genes into different age groups and chose the oldest gene to represent the age of the gene. We identified new genes at the ancestor node of ruminants.

**3.2.4 Membrane anchor and signal peptide prediction**

To identify the presence or absence of potential membrane anchor regions, amino acid sequences of proteins and signal peptide regions were submitted to the TMMM2.0[31] server (http://www.cbs.dtu.dk/services/TMHMM/) and the SignalIP4.1[32] server (http://www.cbs.dtu.dk/services/SignalP/).

**3.2.5 Protein 3D structure simulation**

The 3D structures of all the rumen key genes under positive selection in the common ancestor of Ruminantia were predicated using the I-TASSER website with the URL (https://zhanglab.ccmb.med.umich.edu/I-TASSER/)[33], and then visualized using UCSF Chimera[34].

##### 3.2.6 Protein structure domain analysis

The protein domain architectures were predicated using SMART website with the URL (http://smart.embl-heidelberg.de/)[35].

#### Part 4. Functional assays

##### 4.1 *In vitro* estimation of HMGCS2 enzyme activity

###### 4.1.1 Construction of expression plasmids

*HMGCS2* genes were synthesized by GeneCreate Biological Engineering Co., Ltd. (Wuhan, China), including *HMGCS2* of sheep and human, *HMGCS2* of sheep with five ruminants positively selected sites replaced by the human amino acids and *vice versa*. They were codon-optimized according to *E. coli* preference.

The genes were cloned into vector pET-28a, respectively, by Gibson assembling methods[36]. The plasmids were transformed into DH5α for application, and then they were extracted for further verification. Finally, the correct plasmids were transformed into BL21 (DE3) to be expressed, respectively.

###### 4.1.2 Expression and purification of enzymes in E. coli BL21 (DE3)

Recombinant *E. coli* BL21 cells were grown in 2YT-broth (yeast extract 10 g/L, tryptone 16 g/L and NaCl 5 g/L) containing 50 mg/L ampicillin at 37 °C under good aeration. Induction of 1a-hydroxylase cDNA transcription using the associated promoter was initiated by addition of isopropylthio-β-D-thiogalactoside (IPTG) at a final concentration of 1 nM for approximately 16 h. The cells were then collected by centrifugation and disrupted using a constant cell disruption system. The supernatant was separated from the disrupted cell walls by centrifugation. Thereafter, the protein was purified using $Ni^{2+}$ affinity columns. Subsequently, the enzymes were stored until quantification and activity measurement.

###### 4.1.3 Measurement of enzyme activity

To explore the functional relevance of the sheep, human, sheep with five human sites replaced and human with five sites of ruminants replaced *in vitro*, we synthesized the orthologs and then tested their enzyme catalytic activities by measuring each enzyme activity in a reconstituted system consisting of the enzyme, substrate. The quantification of the HMGCS2 activity were carried out using a spectrophotometry quantitative detection kit for 3-hydroxy-3-methylglutaryl coenzyme A (HMG-COA) synthase activity (GMS50264.6) purchased from GENEMED (Shanghai, China). The enzyme activity was calculated according to the OD 303 nm change of the reconstituted system.

###### 4.1.4 Statistics

The t-test in the SPSS22 software (SPSS, Chicago, IL, USA) was applied to calculate the significance for the HMGCS2 enzyme activity. Differences were considered to be statistically significant when $p < 0.05$.

##### 4.2 Detection of regulatory activity of candidate RSCNEs

###### 4.2.1 Cell lines and cell culture

Sheep and goat fibroblast cells were provided by Guangxi University and were cultured in Dulbecco's Modified Eagle Medium (DMEM) containing 10% FBS (Gibco, Grand Island, NY, USA). All cell lines used in this study were maintained in the specified medium supplemented with 1× Penicillin–Streptomycin (Gibco) and incubated in 5% $CO_2$ at 37 °C.

###### 4.2.2 Cloning and luciferase assays

All the reporter constructs were cloned into pGL-3 promoter plasmids (Promega, Madison, WI, USA). Fragments of the candidate RSCNEs were cloned into pGL3-promoter vector digested by *BamH* I and *Sal* I downstream of the luciferase gene. All constructs were confirmed by sequencing. Transfection of all reporter plasmids constructs was performed using TurboFect (R0531, Thermo Scientific, Waltham, USA). Renilla Luciferase pRL-TK-Rluc (Promega) served as a transfection control, and luciferase expression was subsequently monitored with the dual luciferase assay (Promega) 24 h after transfection. Each luciferase assay was monitored at least five times, independently.

**4.2.3 Statistics**

The t-test in the GraphPad Prism7.0 software (Prism, San Diego, CA, USA) was applied to calculate the significance for the regulatory activity. Differences were considered to be statistically significant when $p < 0.05$.

**4.3 Expression of LYZ1 and DEFB1 in *Pichia pastoris* expression system**

**4.3.1 Strains and plasmids**

*Escherichia coli* (ATCC 25922) and *Staphylococcus aureus* (ATCC 29213), used for lysozyme (LYZ1) and β-defensin (DEFB1) activity assay, were preserved in Northwestern Polytechnic University. pPIC9K plasmid (Invitrogen, Grand Island, NY, USA) were used for construction of the recombinant expression plasmid.

**4.3.2 Reagents, media and sample**

The preparation of LB, yeast extract peptone dextrose (YPD), MD, buffered minimum glycerol medium (BMGY) and buffered minimum methanol medium (BMMY) were performed according to the Multi-Copy Pichia Expression kit (Invitrogen). All other chemicals were of analytical grade.

**4.3.3 Construction of expression plasmids**

Gene was synthesized by GeneCreate Biological Engineering Co., Ltd. (Wuhan, China). *LYZ1* and *DEFB1* genes were codon-optimized according to *P. pastoris* preference. The *LYZ1* sequence was:

AAAAAGTTCGAAAGATGTGAATTGGCTAGAACCTTGAAGAGATTGGGTTTAGATGGTTACAGAGGTGTTTCTTTGGCTAACTGGATGTGTTTGGCTAGATGGGAATCTAACTATAACACTAGAGCTACCAACTACAACAGAGGTGATAAGTCTACTGATTACGGTATCTTCCAGATCAATTCTAGATGGTGGTGTAACGATGGTAAAACTCCAAGAGCTGTTAACGCTTGTAGAATTCCATGTTCTGCTTTGTTGAAGGATGATATTACTCAAGCTGTTGCTTGCGCTAAGAGAGTTGTTTCTGATCCACAAGGTATTAAGGCTTGGGTTGCTTGGAGAAACAAATGCCAAAACAAGGACTTGAGATCCTACGTTAAGGGTTGTAGAGTTCACCACCACCACCACCACTAA after removing the signal peptide sequences. The *DEFB1* sequence was

TCTGGTTTCACTCAAGGTATTAGATCTAGAAGATCCTGCCATAGAAACAA GGGTGTTTGTGCTTTGACTAGATGTCCAAGAAACATGAGACAAATCGGTA CTTGTTTTGGTCCACCAGTTAAGTGTTGTAGGAAGAAGCACCACCACCAC CACCACTAA.

Based on the complete DNA sequence of the LYZ1 gene (GenBank accession: XM_005680191.3), a pair of PCR primers were designed. Forward and reverse primers, synthesized by Sangon (China), were LYZ1-F: 5'-TACGTAGAATTCAA GAGAAAAAAGTTCGAAAGATGTGAATTGGCT-3' with a homologous arm with the vector pPIC9K (underlined) and LYZ1-R: 5'-ATGGTGATGATGATTCGCGGC CGCTTAGTGGTGGTGGTGGTGGTGAACTCTA-3' with a homologous arm with the vector pPIC9K (underlined), respectively. And based on the complete DNA sequence of the *DEFB1* gene (GenBank accession: XM_018042143.1), a pair of PCR primers was designed. Forward and reverse primers, synthesized by Sangon (China), were DEFB1-F: 5'- CGTAGAATTCAAGAGATCTGGTT TCACTCAAGGTAT TA GATCTAGAAGAT -3' with a homologous arm with the vector pPIC9K (underlined) and DEFB1-R: 5'- GCGAATTAATTCGCGGCCGCTTAG TGGTGGTGGTGGTGG TGC -3' with a homologous arm with the vector pPIC9K (underlined), respectively.

The genes were cloned into vector pPIC9K, respectively, by Gibson assembling methods[36]. The plasmids were transformed into DH5α for application, and then the plasmids were extracted for verification. Finally, the correct plasmids were transformed into *P. pastoris* GS115 (Invitrogen) to be expressed, separately.

**4.3.4 Transformation and recombinant plasmid expression**

The pPIC9K-LYZ1 was linearized with *Sac*I, and transformed into *P. pastoris* GS115 by eletroporation using a Gene Pulser apparatus (Bio-Rad, Hercules, CA, USA) according to the manufacturer's instructions. The screening of *P. pastoris* transformants was first carried out on an MD plate according to histidine auxotroph. The integration of LYZ1 into the genome of *P. pastoris* GS115 was confirmed by PCR analysis using primers 5'-AOX1: AATACTGCTGATAGCCTAACGTT and 3'-

AOX1: GCAAATGGCATTCTGACATCCTCT with yeast genomic DNA as a template extracted by the Yeast Genomic DNA Extraction kit. Expression of *LYZ1* in *P. pastoris* GS115 was performed according to the instructions of the Multi-Copy Pichia Expression kit (Invitrogen). Each single colony of transformants was inoculated in 50 ml of BMGY medium in a 500 ml conical flask, and grown at 30˚C on a rotary incubator (220 rpm) until the OD 600 reached 1.0. The cells were then harvested by centrifugation at 3,000 rpm and suspended in 150 ml of BMMY medium in a 500 ml conical flask. The recombinant LYZ1 expression was then induced at 30 ˚C for 6 days by adding 750 μL methanol to a final concentration of 0.5% at 24 h intervals.

All the procedures are the same with the *DEFB1* gene. And the empty vector with purity yeast culture was used as the blank control.

**4.3.5 Protein assay**

Protein concentration was determined by using the BCA-200 Protein Assay kit, with bovine serum albumin as the standard. Sodium dodecyl sulfate-polyacrylamide gel electrophoresis (SDS-PAGE) was performed on a 12.5% gel according to a standard protocol[37], and isolated proteins were visualized by staining with Coomassie Brilliant Blue R-250.

**4.3.6 The recombinant plasmid inhibition zone assay**

We tested the antibacterial ability of LYZ1 and DEFB1 by measuring inhibition zone assay on agarose plates with the Gram-negative *Escherichia coli* (ATCC 25922) bacteria and the Gram-positive *Staphylococcus aureus* (ATCC 29213) bacteria. In disc diffusion method (inhibition zone), 100 μL of the bacteria suspension was cultured at 37 °C for 24 h on agar plates. Each plate was prepared in two circular shapes with 1 cm diameter and then placed in contact with the agar plates. One circular shape was added with 1mL yeast extract with the recombinant plasmid (LYZ or DEFB1), and the other was added with the yeast extract with the empty vector. After incubation of specimens at 37 °C for 24 h, the inhibition zones formed as halo around the specimens with no bacteria growth were photographed.

**Supplementary Figures**

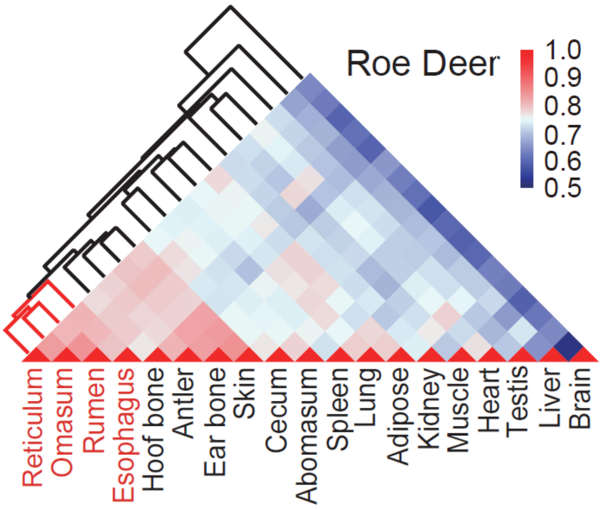

**Fig. S1 |** Hierarchical clustering results showing the relationships among 20 samples of roe deer and a heatmap showing the pairwise Spearman correlations.

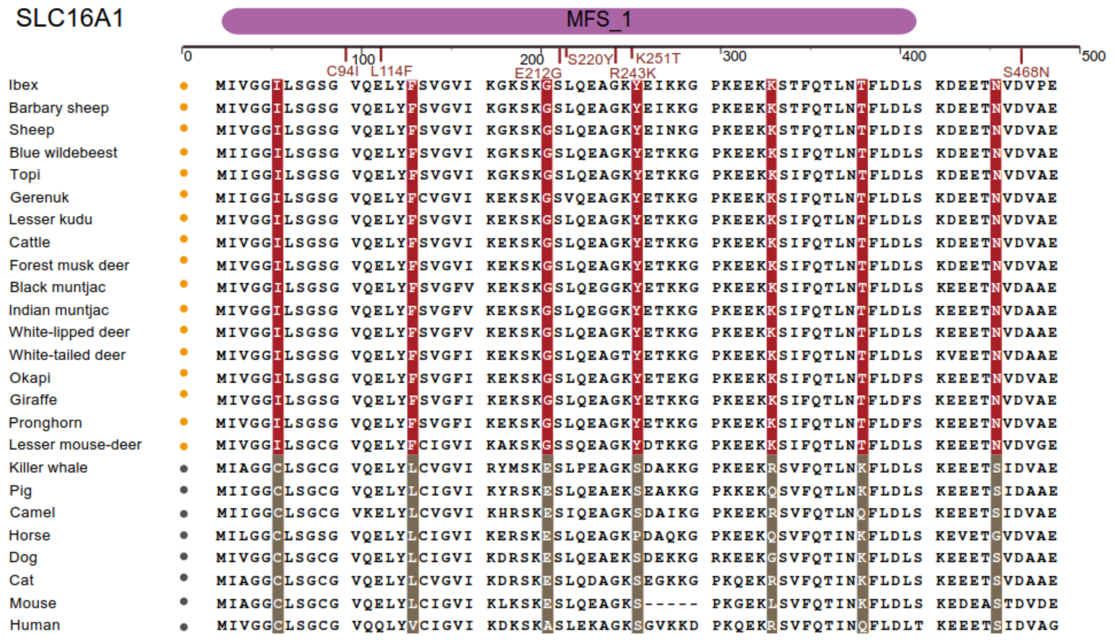

**Fig. S2** | Top panels: Structural domains of the SLC16A1 protein and the location of the ruminant specific mutations. Lower panel: Peptide sequence alignment of SLC16A1. The species followed a yellow circle are belonging to the ruminants and grey circle belonging to the outgroup mammals. The red highlighting indicates ruminant-specific amino acid mutations.

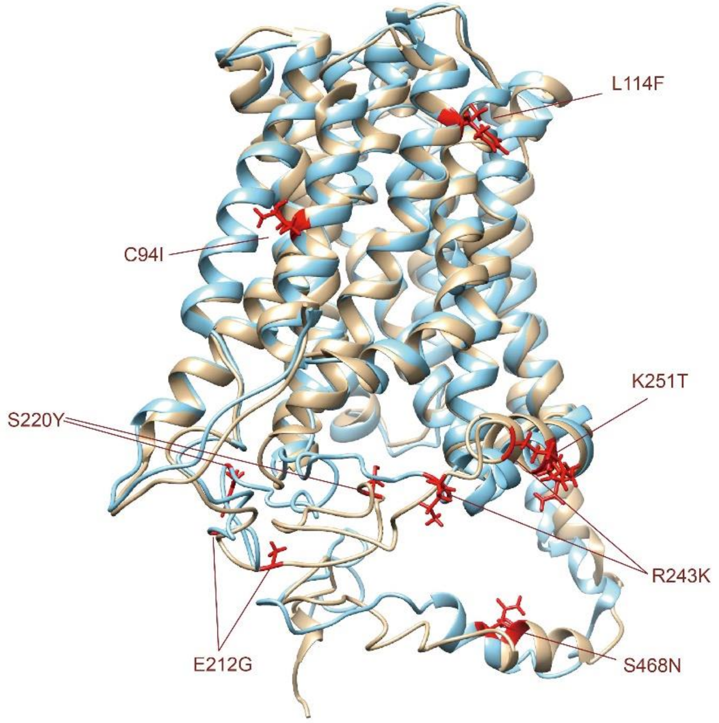

**Fig. S3 |** Predicted tertiary structures of the SLC16A1 of ruminant (blue) and other mammals (orange), respectively. The straight lines point out the positively selected sites with the direction from other mammals to ruminants.

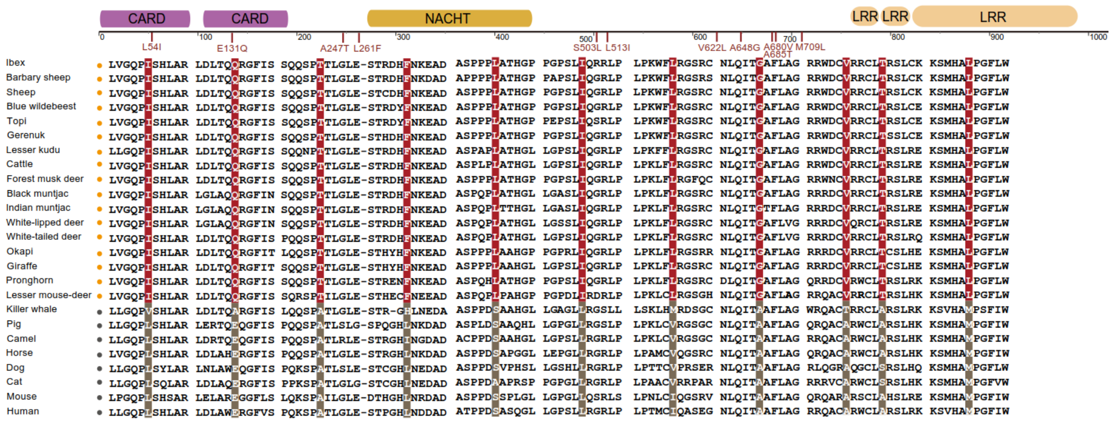

**Fig. S4** | Top panels: Structural domains of the NOD2 protein and the location of the ruminant specific mutations. Lower panel: Peptide sequence alignment of NOD2. The species followed a yellow circle are belonging to the ruminants and grey circle belonging to the outgroup mammals. The red highlighting indicates ruminant-specific amino acid mutations.

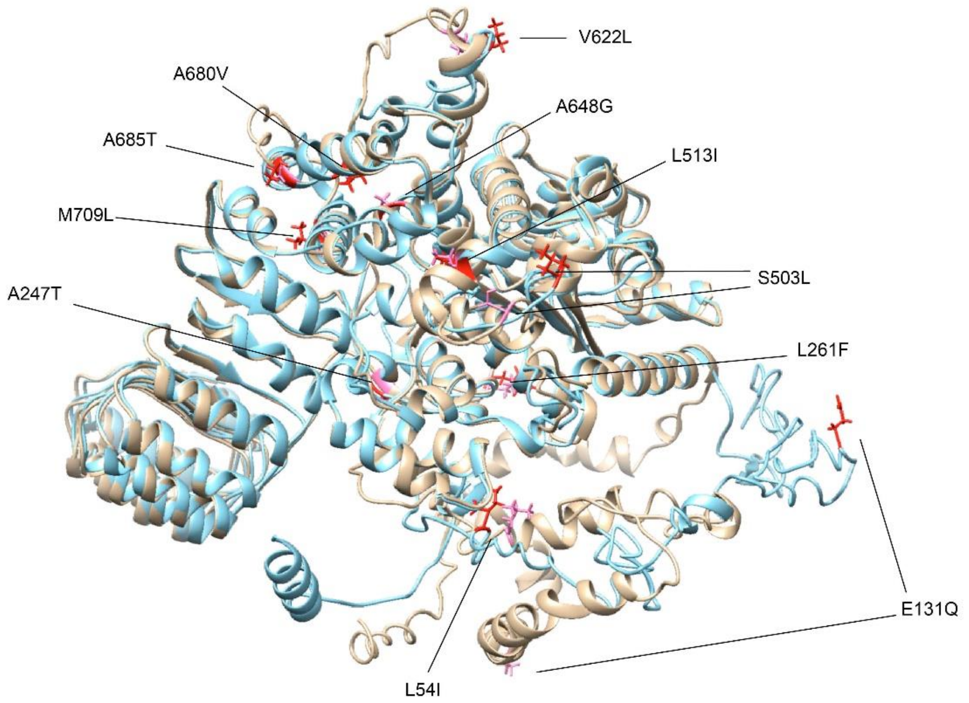

**Fig. S5 |** Predicted tertiary structures of the NOD2 of ruminant (blue) and other mammals (orange), respectively. The straight lines point out the positively selected sites with the direction from other mammals to ruminants.

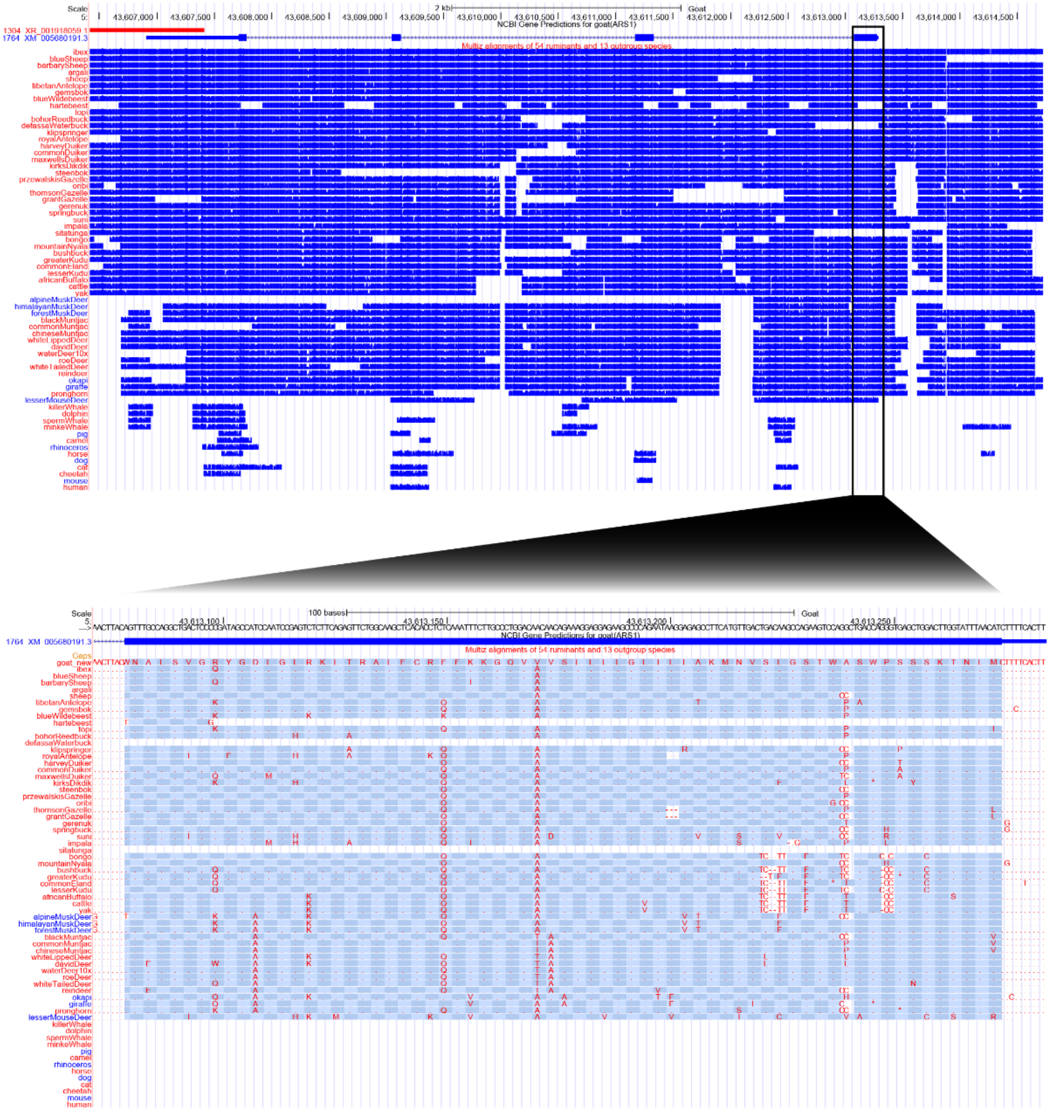

**Fig. S6** | The amino-terminal sequences of mammalian LYZ1 proteins (LOC102171764 in goat) and a probable transmembrane anchor present specifically in ruminants.

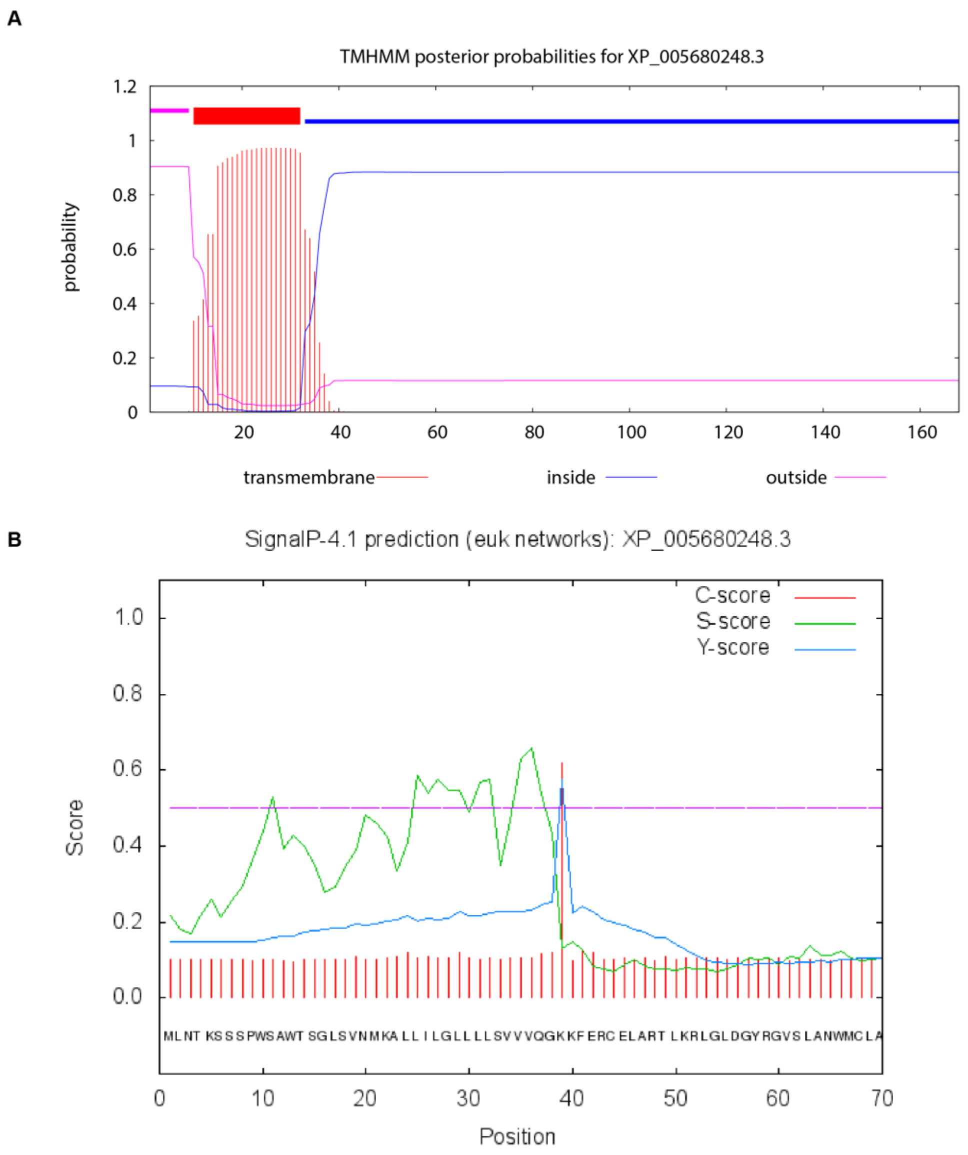

**Fig. S7 |** Transmembrane domain prediction and signal peptide prediction in the goat LYZ1 protein. (**A**) A 22 amino acid transmembrane domain was predicted in the amino terminus of the goat LOC102171764 (LYZ1) protein using TMHMM. (**B**) A 38 amino acid signal peptide was predicted using SignalP.

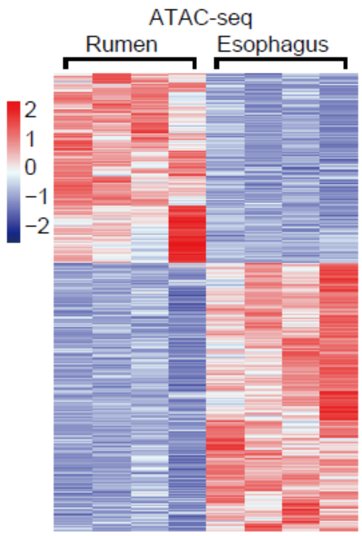

**Fig. S8** | Heatmap showing the differentially accessible peaks from ATAC-seq between the rumen and the esophagus.

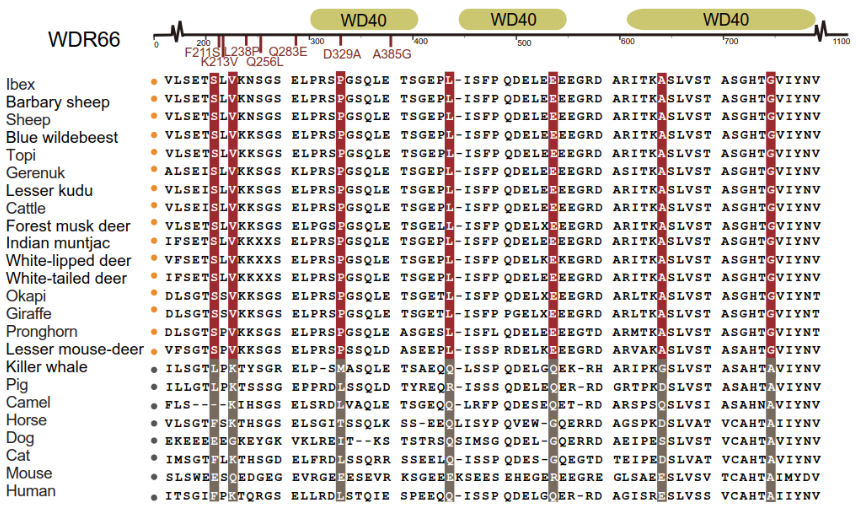

**Fig. S9 |** Top panels: Structural domains of the WDR66 protein and the location of the ruminant specific mutations. Lower panel: Peptide sequence alignment of WDR66. The species followed a yellow circle are belonging to the ruminants and grey circle belonging to the outgroup mammals. The red highlighting indicates ruminant-specific amino acid mutations.

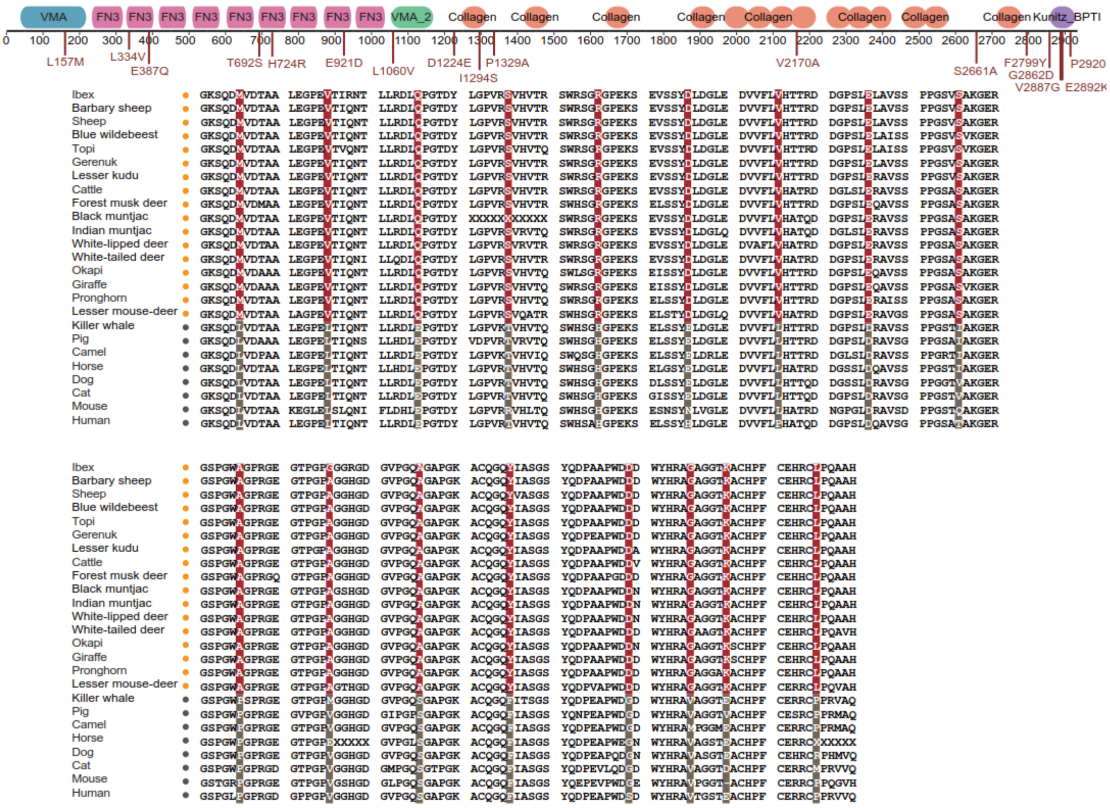

**Fig. S10 |** Top panels: Structural domains of the COL7A1 protein and the location of the ruminant specific mutations. Lower panel: Peptide sequence alignment of COL7A1. The species followed a yellow circle are belonging to the ruminants and grey circle belonging to the outgroup mammals. The red highlighting indicates ruminant-specific amino acid mutations.

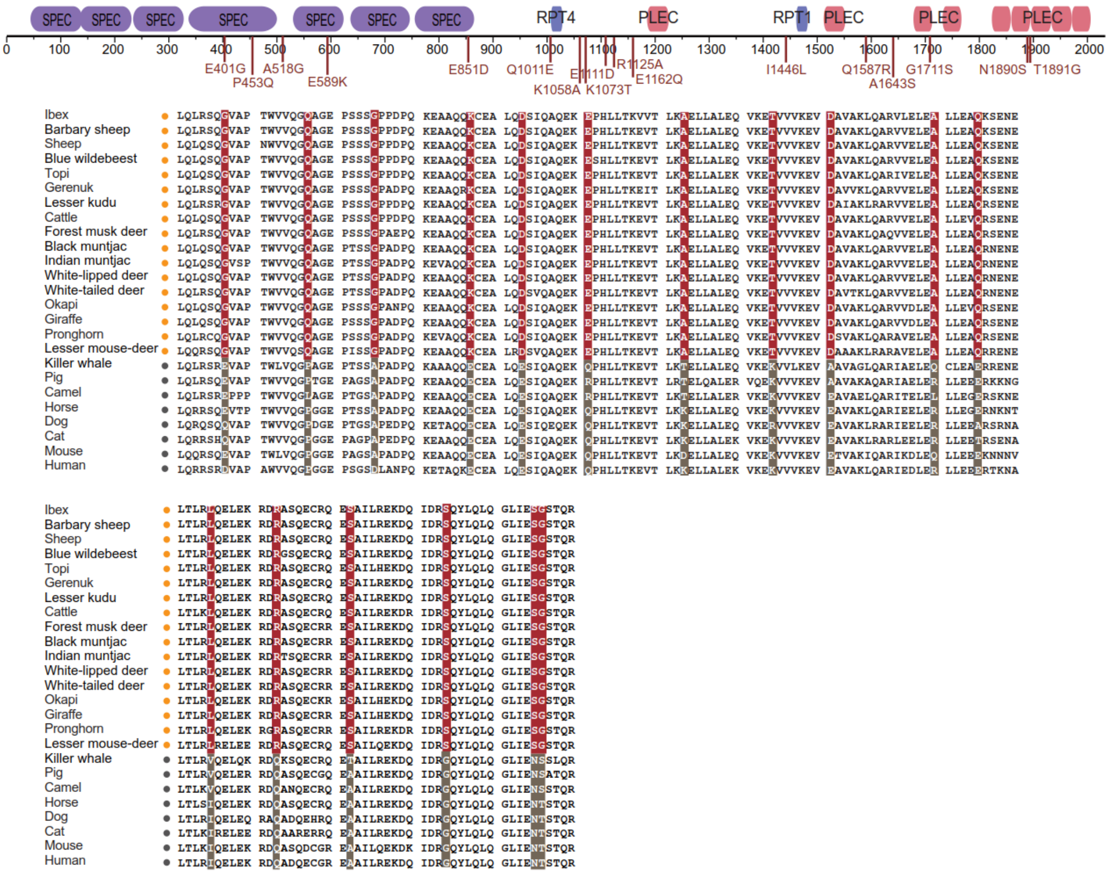

**Fig. S11 |** Top panels: Structural domains of the EVPL protein and the location of the ruminant specific mutations. Lower panel: Peptide sequence alignment of EVPL. The species followed a yellow circle are belonging to the ruminants and grey circle belonging to the outgroup mammals. The red highlighting indicates ruminant-specific amino acid mutations.

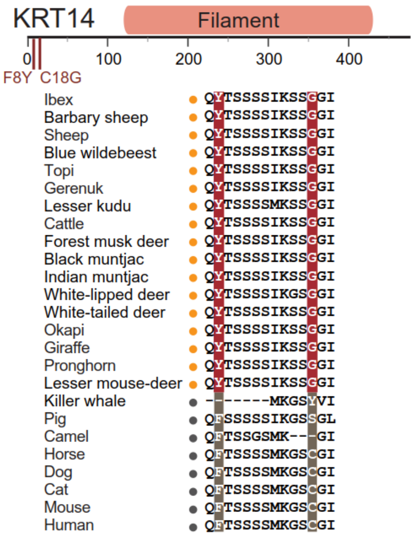

**Fig. S12 |** Top panels: Structural domains of the KRT14 protein and the location of the ruminant specific mutations. Lower panel: Peptide sequence alignment of KRT14. The species followed a yellow circle are belonging to the ruminants and grey circle belonging to the outgroup mammals. The red highlighting indicates ruminant-specific amino acid mutations.

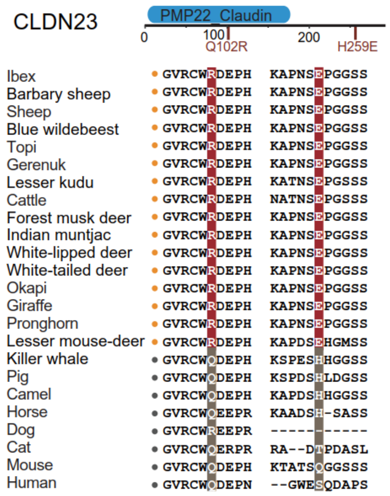

**Fig. S13 |** Top panels: Structural domains of the CLDN23 protein and the location of the ruminant specific mutations. Lower panel: Peptide sequence alignment of CLDN23. The species followed a yellow circle are belonging to the ruminants and grey circle belonging to the outgroup mammals. The red highlighting indicates ruminant-specific amino acid mutations.

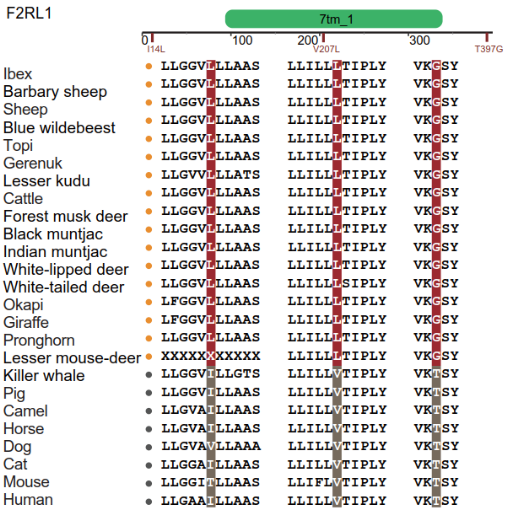

**Fig. S14 |** Top panels: Structural domains of the F2RL1 protein and the location of the ruminant specific mutations. Lower panel: Peptide sequence alignment of F2RL1. The species followed a yellow circle are belonging to the ruminants and grey circle belonging to the outgroup mammals. The red highlighting indicates ruminant-specific amino acid mutations.

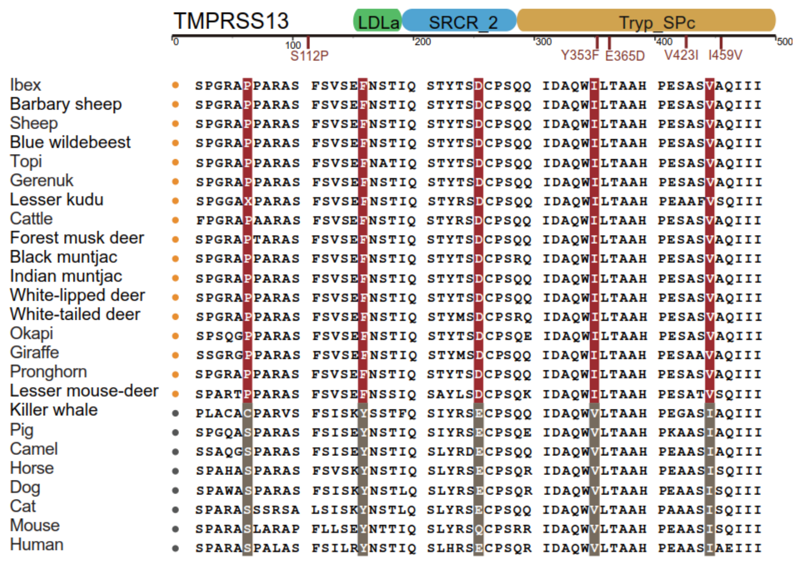

**Fig. S15 |** Top panels: Structural domains of the TMPRSS13 protein and the location of the ruminant specific mutations. Lower panel: Peptide sequence alignment of TMPRSS13. The species followed a yellow circle are belonging to the ruminants and grey circle belonging to the outgroup mammals. The red highlighting indicates ruminant-specific amino acid mutations.

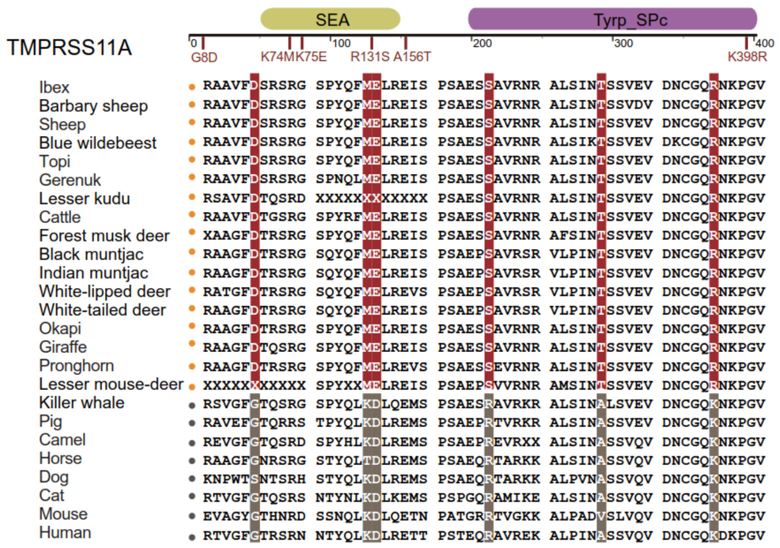

**Fig. S16 |** Top panels: Structural domains of the TMPRSS11A protein and the location of the ruminant specific mutations. Lower panel: Peptide sequence alignment of TMPRSS11A. The species followed a yellow circle are belonging to the ruminants and grey circle belonging to the outgroup mammals. The red highlighting indicates ruminant-specific amino acid mutations.

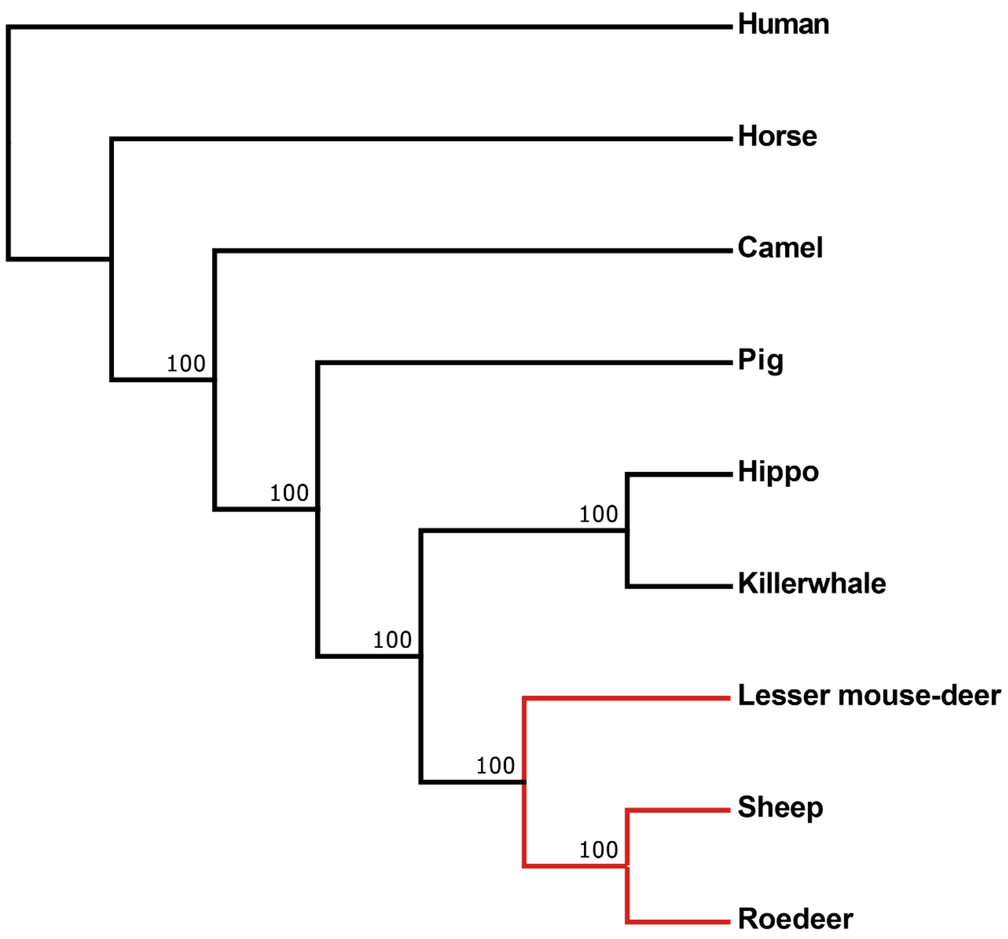

**Fig. S17 |** The ML phylogenetic tree for the major taxonomy of the Cetartiodactyla. A total of 3,316,385 four-fold degenerate sites with 11,567 single-copy orthologous genes in nine species (humans were used to as the outgroup) to construct the ML phylogenetic tree by IQ-Tree. A total of 100 bootstraps are analyzed and the values are marked at each node. All nodes have 100% percent support.

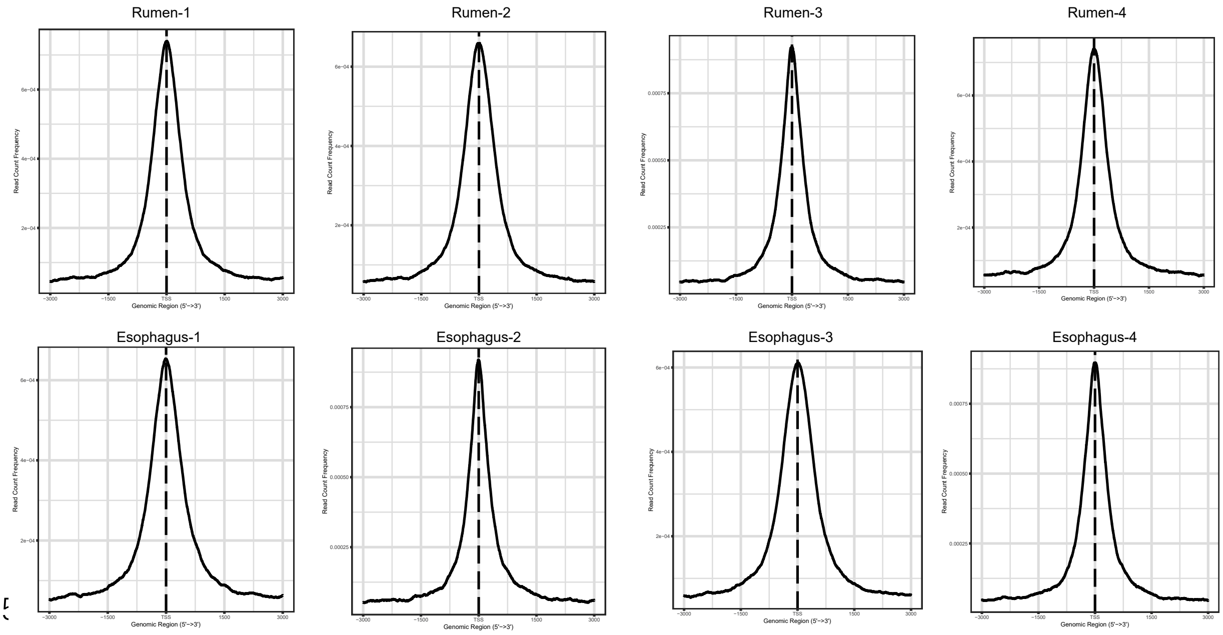

**Fig. S18 |** Distribution of accessible regions around the TSS identified from the rumen and esophagus ATAC-seq samples. The center of accessible regions was used to produce the distribution plots.

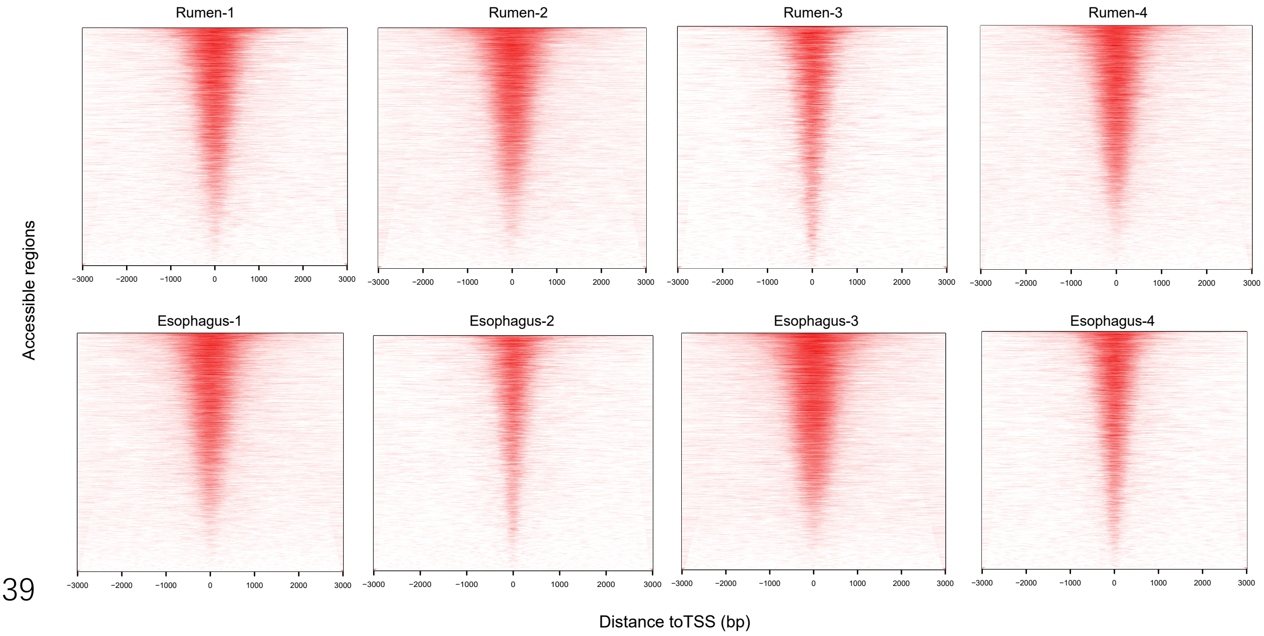

**Fig. S19 |** Heatmap showing the distribution of accessible regions around the TSS identified from the rumen and esophagus ATAC-seq samples.

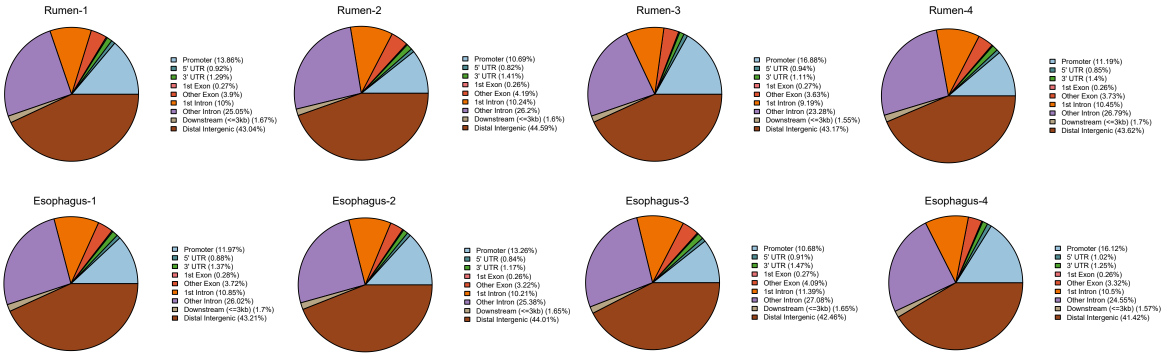

**Fig. S20 |** Pie charts showing the percentage of peaks from the rumen and esophagus ATAC-seq samples.

#### Supplementary Tables

**Supplementary Table 1 |** Details of transcriptomes samples used in this study.

[see the excel file]

**Supplementary Table 2 |** The gene expression profile of 655 rumen specifically expressed genes.

[see the excel file]

**Supplementary Table 3 |** The gene expression profile of 593 FC stomach specifically expressed gene of camels and the tissue source of gene co-expressed with.

[see the excel file]

**Supplementary Table 4 |** The gene expression profile of 375 FC stomach specifically expressed gene of cetaceans and the tissue source of gene co-expressed with.

[see the excel file]

**Supplementary Table 5 |** The gene expression profile of 18 FC stomachs co-expressed genes from their esophagus.

| Official name | Rumen of sheep | The FC stomach of cetacean | The FC stomach of camel |
|---|---|---|---|
| *SULT2B1* | 198.05 | 128.98 | 185.13 |
| *PKP1* | 288.50 | 39.47 | 433.03 |
| *SDCBP2* | 94.37 | 9.68 | 48.12 |
| *LOC101106941* | 277.21 | 0.00 | 0.00 |
| *DLK2* | 35.95 | 34.33 | 78.27 |
| *ANKRD35* | 25.87 | 7.80 | 95.59 |
| *PAX9* | 85.44 | 15.18 | 32.88 |
| *VSIG10L* | 56.78 | 21.15 | 44.28 |
| *TRIM29* | 223.12 | 55.94 | 1145.62 |
| *TMEM40* | 113.23 | 161.27 | 91.70 |
| *EVPL* | 186.51 | 42.39 | 317.39 |
| *C12H1orf106* | 29.02 | 0.00 | 0.00 |
| *GRHL3* | 37.03 | 8.56 | 62.47 |
| *ZNF750* | 63.17 | 17.66 | 73.31 |
| *TGM1* | 546.80 | 236.87 | 395.99 |
| *LYPD3* | 735.26 | 138.79 | 3408.74 |
| *LOC101116141* | 880.86 | 91.33 | 711.98 |
| *IVL* | 1519.64 | 357.51 | 1082.81 |

**Supplementary Table 6 |** GO enrichment analysis of 18 FC stomach of camels, cetaceans and ruminants co-expressed genes from their esophagus. BP denotes Biological Process, MF denotes Molecular Function, and CC denotes Cellular Component.

| ID | Type | Description | *p* value | adjusted *p* value | list1InGO | list1OutGO | list2InGO | list2OutGO |
|---|---|---|---|---|---|---|---|---|
| GO:0030216 | BP | keratinocyte differentiation | 2.91E-04 | 0.0098 | 3 | 10 | 31 | 10659 |
| GO:0001533 | CC | cornified envelope | 0.011793 | 0.1429 | 2 | 14 | 12 | 13162 |
| GO:0018149 | BP | peptide cross-linking | 0.013434 | 0.2054 | 2 | 10 | 16 | 10659 |
| GO:0008544 | BP | epidermis development | 0.013434 | 0.2054 | 2 | 10 | 16 | 10659 |
| GO:0045111 | CC | intermediate filament cytoskeleton | 0.034038 | 0.2016 | 2 | 14 | 35 | 13162 |
| GO:0001228 | MF | transcriptional activator activity, RNA polymerase II transcription regulatory region sequence-specific binding | 0.039732 | 0.4097 | 2 | 9 | 53 | 10488 |

**Supplementary Table 7 |** The gene expression and fold change of DEGs between the rumen and the FC stomach of camels. The UP represents the gene is upregulated in the rumen; the DOWN represents the gene is downregulated in the rumen when compared to the FC stomach of camels.

| Official name | Rumen of sheep | The FC stomach of camel | Fold Change of Sheep/Camel | Rumen |
|---|---|---|---|---|
| *PAQR7* | 1.911558 | 0.306675 | 6.2331719 | UP |
| *GJB2* | 138.9196 | 61.1398 | 2.2721632 | UP |
| *TEAD4* | 5.8731301 | 2.58409 | 2.272804 | UP |
| *BEST2* | 23.868862 | 0.136937 | 174.30543 | UP |
| *NTF4* | 2.923117 | 0.388378 | 7.5264742 | UP |
| *TWIST2* | 16.231501 | 1.12147 | 14.473415 | UP |
| *LOC101106941* | 277.21219 | 30.649 | 9.0447385 | UP |
| *IL36G* | 40.154309 | 14.681 | 2.7351208 | UP |
| *PRRT4* | 3.247948 | 0.106991 | 30.357208 | UP |
| *RIN1* | 19.890148 | 7.51326 | 2.6473392 | UP |
| *LOC101104705* | 148.01907 | 29.351 | 5.0430675 | UP |
| *IL20RB* | 62.029479 | 22.5241 | 2.753916 | UP |
| *USP35* | 6.685347 | 1.37795 | 4.8516615 | UP |
| *C3H22orf23* | 5.1275285 | 2.17833 | 2.3538805 | UP |
| *ASPRV1* | 68.646751 | 3.94189 | 17.41468 | UP |
| *PLEK2* | 68.003258 | 30.286 | 2.2453694 | UP |
| *WFIKKN1* | 10.426244 | 2.75436 | 3.7853599 | UP |
| *KLHL29* | 3.4505038 | 1.63671 | 2.108195 | UP |
| *C20H6orf141* | 11.415225 | 3.26339 | 3.4979653 | UP |
| *IL36RN* | 153.03899 | 16.4413 | 9.3082047 | UP |
| *SPTSSB* | 60.127381 | 18.0788 | 3.3258502 | UP |
| *KLK9* | 37.228439 | 3.14934 | 11.821029 | UP |
| *LOC101103862* | 707.79498 | 8.19744 | 86.343417 | UP |
| *VSTM2L* | 21.245537 | 2.08555 | 10.187019 | UP |
| *SRPK3* | 10.645239 | 1.45319 | 7.3254285 | UP |
| *FAM83B* | 18.08906 | 3.69837 | 4.8910899 | UP |
| *ZACN* | 7.692018 | 0.001 | 7692.018 | UP |
| *LOC101121414* | 6.5828919 | 2.4795 | 2.6549272 | UP |
| *NPAS1* | 6.030153 | 2.5147 | 2.3979612 | UP |
| *CD70* | 3.024992 | 0.156227 | 19.362799 | UP |
| *CD164L2* | 14.754008 | 0.001 | 14754.008 | UP |
| *AHNAK2* | 9.03539 | 2.90061 | 3.1149965 | UP |
| *LOC105608435* | 10.020885 | 4.00822 | 2.5000836 | UP |
| *AHRR* | 8.042508 | 2.53616 | 3.1711359 | UP |
| *VTCN1* | 7.740344 | 0.22725 | 34.06092 | UP |
| *COL7A1* | 146.03317 | 12.6261 | 11.565976 | UP |
| *B4GALNT3* | 60.219162 | 4.92909 | 12.217095 | UP |

| | | | | |
|---|---|---|---|---|
| *L3MBTL1* | 2.806673 | 1.03394 | 2.7145415 | UP |
| *DSC3* | 93.864174 | 37.789 | 2.483902 | UP |
| *LOC101112304* | 15279.522 | 0.001 | 15279522 | UP |
| *LOC101102714* | 10660.87 | 0.001 | 10660870 | UP |
| *LOC105613035* | 8160.1792 | 0.001 | 8160179.2 | UP |
| *LOC101113601* | 7766.5908 | 0.001 | 7766590.8 | UP |
| *HAS3* | 11.87929 | 4.30417 | 2.7599491 | UP |
| *A2ML1* | 6078.8599 | 0.001 | 6078859.9 | UP |
| *CCR10* | 2.714588 | 0.997137 | 2.7223822 | UP |
| *LOC101110777* | 5062.0308 | 0.001 | 5062030.8 | UP |
| *LOC101102105* | 4067.6902 | 0.001 | 4067690.2 | UP |
| *LOC101111178* | 3783.1057 | 0.001 | 3783105.7 | UP |
| *SNX31* | 15.884428 | 5.29311 | 3.0009631 | UP |
| *PRD-SPRRII* | 3708.9065 | 0.001 | 3708906.5 | UP |
| *LOC101104114* | 3006.6772 | 0.001 | 3006677.2 | UP |
| *LOC105615728* | 2962.5061 | 0.001 | 2962506.1 | UP |
| *CRISP3* | 1638.9224 | 0.001 | 1638922.4 | UP |
| *IVL* | 1519.6367 | 0.001 | 1519636.7 | UP |
| *LOC101107119* | 1239.1457 | 0.001 | 1239145.7 | UP |
| *VSIG2* | 15.634018 | 3.98627 | 3.9219667 | UP |
| *ACACB* | 12.159775 | 3.99821 | 3.0413047 | UP |
| *CCDC183* | 17.821924 | 6.03152 | 2.9547981 | UP |
| *LOC101121219* | 1217.0025 | 0.001 | 1217002.5 | UP |
| *LYPD2* | 1000.9 | 0.001 | 1000900 | UP |
| *TRIM62* | 12.532437 | 3.29566 | 3.8027093 | UP |
| *C7H14orf169* | 17.815302 | 8.19867 | 2.1729502 | UP |
| *HTR2A* | 3.628845 | 1.09939 | 3.3007804 | UP |
| *CPA6* | 4.304087 | 1.26286 | 3.408206 | UP |
| *EREG* | 12.742299 | 5.10363 | 2.4967129 | UP |
| *KRT79* | 955.04913 | 0.001 | 955049.13 | UP |
| *TMPRSS13* | 67.041283 | 24.2459 | 2.7650565 | UP |
| *LOC101116141* | 880.86263 | 0.001 | 880862.63 | UP |
| *LOC101106610* | 720.6295 | 0.001 | 720629.5 | UP |
| *LOC101122351* | 715.22168 | 0.001 | 715221.68 | UP |
| *LOC101104808* | 698.79773 | 0.001 | 698797.73 | UP |
| *LOC101111992* | 621.79816 | 0.001 | 621798.16 | UP |
| *LOC105604529* | 523.99908 | 0.001 | 523999.08 | UP |
| *SLC26A3* | 457.02723 | 0.001 | 457027.23 | UP |
| *KCNK9* | 4.23823 | 1.39798 | 3.0316814 | UP |
| *SBD2* | 452.69498 | 0.001 | 452694.98 | UP |
| *UGT1A9* | 436.05011 | 0.001 | 436050.11 | UP |
| *LOC101108147* | 414.42062 | 0.001 | 414420.62 | UP |

| | | | | |
|---|---|---|---|---|
| *LOC101103612* | 403.29273 | 0.001 | 403292.73 | UP |
| *IL36A* | 323.76747 | 0.001 | 323767.47 | UP |
| *LOC101119393* | 313.42225 | 0.001 | 313422.25 | UP |
| *LOC101115964* | 303.13001 | 0.001 | 303130.01 | UP |
| *ACTA1* | 279.39099 | 0.001 | 279390.99 | UP |
| *DENND2C* | 40.098075 | 14.8992 | 2.6912905 | UP |
| *LOC101107809* | 260.71158 | 0.001 | 260711.58 | UP |
| *PI3* | 258.79388 | 0.001 | 258793.88 | UP |
| *LOC101115172* | 242.40593 | 0.001 | 242405.93 | UP |
| *LOC101112555* | 239.55261 | 0.001 | 239552.61 | UP |
| *STC2* | 12.993917 | 4.39112 | 2.959135 | UP |
| *SERPINB12* | 205.45187 | 0.001 | 205451.87 | UP |
| *LOC105614373* | 203.7187 | 0.001 | 203718.7 | UP |
| *KRT23* | 196.98567 | 0.001 | 196985.67 | UP |
| *DSG3* | 623.44115 | 233.22 | 2.6731891 | UP |
| *JPH2* | 85.951279 | 33.394 | 2.573854 | UP |
| *GGT6* | 25.650059 | 7.91594 | 3.2403048 | UP |
| *LOC101121267* | 184.48232 | 0.001 | 184482.32 | UP |
| *LOC101117764* | 182.22461 | 0.001 | 182224.61 | UP |
| *LOC101120106* | 164.77068 | 0.001 | 164770.68 | UP |
| *APOBEC3Z1* | 146.42738 | 0.001 | 146427.38 | UP |
| *LOC101117163* | 138.39495 | 0.001 | 138394.95 | UP |
| *SCEL* | 134.95093 | 0.001 | 134950.93 | UP |
| *C1H2orf54* | 133.08426 | 0.001 | 133084.26 | UP |
| *VILL* | 39.628658 | 12.8796 | 3.0768547 | UP |
| *LOC443322* | 122.75221 | 0.001 | 122752.21 | UP |
| *LOC101102344* | 119.86746 | 0.001 | 119867.46 | UP |
| *LOC101108071* | 110.27437 | 0.001 | 110274.37 | UP |
| *PLA2G2F* | 107.20315 | 0.001 | 107203.15 | UP |
| *KLHL41* | 57.734577 | 16.7488 | 3.4470874 | UP |
| *IL19* | 99.40583 | 0.001 | 99405.83 | UP |
| *RBPMS2* | 95.175972 | 0.001 | 95175.972 | UP |
| *KRT80* | 91.305618 | 0.001 | 91305.618 | UP |
| *LOC101106641* | 90.530417 | 0.001 | 90530.417 | UP |
| *RPTN* | 80.383011 | 0.001 | 80383.011 | UP |
| *ZNF185* | 75.698352 | 0.001 | 75698.352 | UP |
| *LOC101118712* | 75.639404 | 0.001 | 75639.404 | UP |
| *LOC106991113* | 73.218643 | 0.001 | 73218.643 | UP |
| *ATP6V1C2* | 71.623978 | 0.001 | 71623.978 | UP |
| *KRT36* | 9886.4551 | 0.14044 | 70396.291 | UP |
| *KRT34* | 64.546219 | 0.001 | 64546.219 | UP |
| *RBM24* | 21.875119 | 8.39911 | 2.6044568 | UP |

| | | | | |
|---|---|---|---|---|
| *FUT5* | 57.321555 | 0.001 | 57321.555 | UP |
| *SERPINB13* | 57.01982 | 0.001 | 57019.82 | UP |
| *LOC101102969* | 56.238964 | 0.001 | 56238.964 | UP |
| *SLURP1* | 53.910942 | 0.001 | 53910.942 | UP |
| *LOC101103602* | 51.176943 | 0.001 | 51176.943 | UP |
| *LOC101107619* | 48.291035 | 0.001 | 48291.035 | UP |
| *ANKRD9* | 46.081627 | 0.001 | 46081.627 | UP |
| *FGB* | 16.158342 | 7.25417 | 2.2274557 | UP |
| *LOC101115983* | 45.111118 | 0.001 | 45111.118 | UP |
| *SOX21* | 44.414188 | 0.001 | 44414.188 | UP |
| *TNFRSF6B* | 41.547382 | 0.001 | 41547.382 | UP |
| *LOC101105114* | 40.204201 | 0.001 | 40204.201 | UP |
| *MUC21* | 1014.2468 | 268.466 | 3.777934 | UP |
| *FAM127A* | 38.44585 | 0.001 | 38445.85 | UP |
| *LOC101116036* | 37.639004 | 0.001 | 37639.004 | UP |
| *GPT* | 34.753342 | 0.001 | 34753.342 | UP |
| *LOC105602022* | 34.534691 | 0.001 | 34534.691 | UP |
| *CARD14* | 34.152442 | 0.001 | 34152.442 | UP |
| *B4GALNT2* | 33.783635 | 0.001 | 33783.635 | UP |
| *LOC101114663* | 33.395721 | 0.001 | 33395.721 | UP |
| *FAM150A* | 32.416817 | 0.001 | 32416.817 | UP |
| *LOC101114920* | 31.718086 | 0.001 | 31718.086 | UP |
| *ANKRD23* | 31.143512 | 0.001 | 31143.512 | UP |
| *LOC101105876* | 30.345701 | 0.001 | 30345.701 | UP |
| *SLC5A1* | 75.425423 | 27.0585 | 2.7874946 | UP |
| *CEACAM19* | 29.690464 | 0.001 | 29690.464 | UP |
| *CHST4* | 28.612474 | 0.001 | 28612.474 | UP |
| *KLC3* | 28.42285 | 0.001 | 28422.85 | UP |
| *LOC101107541* | 27.69426 | 0.001 | 27694.26 | UP |
| *LOC105616317* | 27.035503 | 0.001 | 27035.503 | UP |
| *LOC101103644* | 26.325151 | 0.001 | 26325.151 | UP |
| *ADGRF4* | 24.721505 | 0.001 | 24721.505 | UP |
| *CDR1* | 23.526815 | 0.001 | 23526.815 | UP |
| *LYPD5* | 91.734909 | 27.9778 | 3.2788464 | UP |
| *LOC101106229* | 23.180359 | 0.001 | 23180.359 | UP |
| *S100A5* | 22.641116 | 0.001 | 22641.116 | UP |
| *LOC101113860* | 33.355256 | 13.2671 | 2.5141332 | UP |
| *P2RY2* | 22.63269 | 0.001 | 22632.69 | UP |
| *UGT1A4* | 21.085617 | 0.001 | 21085.617 | UP |
| *GAL3ST2* | 20.688736 | 0.001 | 20688.736 | UP |
| *ACPP* | 20.055654 | 0.001 | 20055.654 | UP |
| *LOC106990121* | 19.693312 | 0.001 | 19693.312 | UP |

| | | | | |
|---|---|---|---|---|
| *LOC105616285* | 19.495518 | 0.001 | 19495.518 | UP |
| *SFTPC* | 19.144745 | 0.001 | 19144.745 | UP |
| *SPINK9* | 18.498169 | 0.001 | 18498.169 | UP |
| *PPP2R2C* | 17.989313 | 0.001 | 17989.313 | UP |
| *LOC101107098* | 17.471764 | 0.001 | 17471.764 | UP |
| *GSDMC* | 17.381283 | 0.001 | 17381.283 | UP |
| *CAND2* | 16.865402 | 0.001 | 16865.402 | UP |
| *LOC101110524* | 16.524918 | 0.001 | 16524.918 | UP |
| *PGLYRP4* | 16.229507 | 0.001 | 16229.507 | UP |
| *LOC101113335* | 15.405745 | 0.001 | 15405.745 | UP |
| *LOC105608915* | 14.530236 | 0.001 | 14530.236 | UP |
| *LOC105614885* | 13.989855 | 0.001 | 13989.855 | UP |
| *LOC105612196* | 13.947534 | 0.001 | 13947.534 | UP |
| *FCER2* | 13.908662 | 0.001 | 13908.662 | UP |
| *LOC105614940* | 13.655092 | 0.001 | 13655.092 | UP |
| *STK32C* | 13.119646 | 0.001 | 13119.646 | UP |
| *LOC101104661* | 12.933213 | 0.001 | 12933.213 | UP |
| *LOC105616478* | 12.211966 | 0.001 | 12211.966 | UP |
| *HOXB7* | 12.075035 | 0.001 | 12075.035 | UP |
| *TRPM6* | 12.066585 | 0.001 | 12066.585 | UP |
| *LOC101120775* | 12.005045 | 0.001 | 12005.045 | UP |
| *LTB4R2* | 11.963988 | 0.001 | 11963.988 | UP |
| *WNT7B* | 11.953721 | 0.001 | 11953.721 | UP |
| *HSPA1A* | 11.913466 | 0.001 | 11913.466 | UP |
| *SERPINB11* | 11.745147 | 0.001 | 11745.147 | UP |
| *LOC105604082* | 10.931283 | 0.001 | 10931.283 | UP |
| *KIF26A* | 10.660532 | 0.001 | 10660.532 | UP |
| *LOC105611786* | 10.545382 | 0.001 | 10545.382 | UP |
| *ADTRP* | 10.454161 | 0.001 | 10454.161 | UP |
| *LOC101121036* | 10.153899 | 0.001 | 10153.899 | UP |
| *LOC105605155* | 9.557295 | 0.001 | 9557.295 | UP |
| *C13H10orf113* | 9.206914 | 0.001 | 9206.914 | UP |
| *LOC105610609* | 8.921119 | 0.001 | 8921.119 | UP |
| *IL17REL* | 8.7226088 | 0.001 | 8722.6088 | UP |
| *ATP13A5* | 8.6974492 | 0.001 | 8697.4492 | UP |
| *LOC105612090* | 8.549753 | 0.001 | 8549.753 | UP |
| *LOC101117587* | 8.327113 | 0.001 | 8327.113 | UP |
| *LRFN4* | 8.313799 | 0.001 | 8313.799 | UP |
| *A1BG* | 8.307885 | 0.001 | 8307.885 | UP |
| *LOC101117120* | 8.285954 | 0.001 | 8285.954 | UP |
| *LOC101105566* | 8.140181 | 0.001 | 8140.181 | UP |
| *ADGRF2* | 7.9526373 | 0.001 | 7952.6373 | UP |

| | | | | |
|---|---|---|---|---|
| *WDR38* | 3.3837037 | 1.54204 | 2.1943035 | UP |
| *LOC101110696* | 7.895658 | 0.001 | 7895.658 | UP |
| *BHLHE23* | 7.758808 | 0.001 | 7758.808 | UP |
| *LOC105608606* | 7.647254 | 0.001 | 7647.254 | UP |
| *MCCD1* | 7.555327 | 0.001 | 7555.327 | UP |
| *LOC105606870* | 7.5335601 | 0.001 | 7533.5601 | UP |
| *LOC101120435* | 7.258325 | 0.001 | 7258.325 | UP |
| *LOC105604589* | 7.234852 | 0.001 | 7234.852 | UP |
| *LOC106991452* | 6.686908 | 0.001 | 6686.908 | UP |
| *LOC101105174* | 6.561944 | 0.001 | 6561.944 | UP |
| *LOC106991450* | 6.491086 | 0.001 | 6491.086 | UP |
| *LOC101115778* | 6.490631 | 0.001 | 6490.631 | UP |
| *VAX2* | 6.132115 | 0.001 | 6132.115 | UP |
| *LOC105615508* | 5.752361 | 0.001 | 5752.361 | UP |
| *C1QL1* | 5.433838 | 0.001 | 5433.838 | UP |
| *LOC101104501* | 5.417474 | 0.001 | 5417.474 | UP |
| *KLF13* | 5.207161 | 0.001 | 5207.161 | UP |
| *POC1A* | 5.1958163 | 0.001 | 5195.8163 | UP |
| *LOC101105258* | 5.052308 | 0.001 | 5052.308 | UP |
| *SH2D7* | 4.951905 | 0.001 | 4951.905 | UP |
| *LOC101122314* | 4.784063 | 0.001 | 4784.063 | UP |
| *LOC105602127* | 4.748694 | 0.001 | 4748.694 | UP |
| *LOC101120688* | 4.672022 | 0.001 | 4672.022 | UP |
| *LOC106990150* | 5.030138 | 1.56477 | 3.2146181 | UP |
| *LOC101111245* | 4.639668 | 0.001 | 4639.668 | UP |
| *LOC101104703* | 4.55153 | 0.001 | 4551.53 | UP |
| *LOC101116196* | 4.5095823 | 0.001 | 4509.5823 | UP |
| *NKX2-2* | 4.422982 | 0.001 | 4422.982 | UP |
| *APELA* | 4.212973 | 0.001 | 4212.973 | UP |
| *SPATA12* | 4.126355 | 0.001 | 4126.355 | UP |
| *LOC105615865* | 4.101434 | 0.001 | 4101.434 | UP |
| *LOC105605056* | 4.023712 | 0.001 | 4023.712 | UP |
| *LOC101106977* | 3.967016 | 0.001 | 3967.016 | UP |
| *LOC101114497* | 3.944958 | 0.001 | 3944.958 | UP |
| *LOC105606498* | 3.920981 | 0.001 | 3920.981 | UP |
| *ARSE* | 3.9071256 | 0.001 | 3907.1256 | UP |
| *LOC106990135* | 3.8417915 | 0.001 | 3841.7915 | UP |
| *LOC101113726* | 3.769403 | 0.001 | 3769.403 | UP |
| *LOC101113189* | 3.744442 | 0.001 | 3744.442 | UP |
| *LOC105616091* | 3.708944 | 0.001 | 3708.944 | UP |
| *LRRN1* | 6.686132 | 1.94715 | 3.4338043 | UP |
| *TLR5* | 3.6566019 | 0.001 | 3656.6019 | UP |

| | | | | |
|---|---|---|---|---|
| *POU2F3* | 7.0183467 | 1.87832 | 3.7365021 | UP |
| *LOC106991207* | 3.549698 | 0.001 | 3549.698 | UP |
| *LOC106991203* | 3.465721 | 0.001 | 3465.721 | UP |
| *LOC105616415* | 3.339548 | 0.001 | 3339.548 | UP |
| *LOC105614635* | 3.298318 | 0.001 | 3298.318 | UP |
| *LOC105610456* | 3.195928 | 0.001 | 3195.928 | UP |
| *CTRC* | 3.138362 | 0.001 | 3138.362 | UP |
| *LOC105604369* | 3.077245 | 0.001 | 3077.245 | UP |
| *LOC101107259* | 3.052632 | 0.001 | 3052.632 | UP |
| *CD207* | 7.652756 | 1.92049 | 3.9847935 | UP |
| *ATP6V1B1* | 423.49908 | 0.147714 | 2867.0206 | UP |
| *ACSM4* | 2.836116 | 0.001 | 2836.116 | UP |
| *LOC105607154* | 2.798793 | 0.001 | 2798.793 | UP |
| *LOC105604842* | 2.780288 | 0.001 | 2780.288 | UP |
| *LOC106991405* | 2.765373 | 0.001 | 2765.373 | UP |
| *LOC105615893* | 2.734533 | 0.001 | 2734.533 | UP |
| *LOC101101862* | 2.7211722 | 0.001 | 2721.1722 | UP |
| *LOC105615734* | 2.678692 | 0.001 | 2678.692 | UP |
| *LOC105608895* | 2.677712 | 0.001 | 2677.712 | UP |
| *C4H7orf31* | 2.648178 | 0.001 | 2648.178 | UP |
| *FCN3* | 2.637651 | 0.001 | 2637.651 | UP |
| *CD177* | 207.62043 | 0.0788412 | 2633.4001 | UP |
| *RBP2* | 412.65064 | 0.158464 | 2604.0655 | UP |
| *LOC105611549* | 2.5842173 | 0.001 | 2584.2173 | UP |
| *LOC101105379* | 2.581682 | 0.001 | 2581.682 | UP |
| *SLC9A2* | 87.317711 | 0.0349537 | 2498.0964 | UP |
| *CYP4F21* | 2.435596 | 0.001 | 2435.596 | UP |
| *LOC101107795* | 2.416286 | 0.001 | 2416.286 | UP |
| *LOC105604880* | 2.347296 | 0.001 | 2347.296 | UP |
| *LOC105616289* | 2.327093 | 0.001 | 2327.093 | UP |
| *LOC101112481* | 2.252598 | 0.001 | 2252.598 | UP |
| *LOC101103439* | 202.9718 | 0.0957064 | 2120.7756 | UP |
| *LOC105614078* | 2.086725 | 0.001 | 2086.725 | UP |
| *TMEM257* | 2.045439 | 0.001 | 2045.439 | UP |
| *LOC105615805* | 2.038133 | 0.001 | 2038.133 | UP |
| *LOC106991649* | 2.016159 | 0.001 | 2016.159 | UP |
| *LOC105612436* | 1.962022 | 0.001 | 1962.022 | UP |
| *LOC105609952* | 1.956774 | 0.001 | 1956.774 | UP |
| *LOC106990488* | 1.930994 | 0.001 | 1930.994 | UP |
| *LOC101109940* | 1.887533 | 0.001 | 1887.533 | UP |
| *LOC105604781* | 1.854786 | 0.001 | 1854.786 | UP |
| *LOC106990202* | 1.849398 | 0.001 | 1849.398 | UP |

| | | | | |
|---|---|---|---|---|
| *MIP* | 1.83593 | 0.001 | 1835.93 | UP |
| *RNF224* | 12.764742 | 4.91109 | 2.5991668 | UP |
| *LOC101116576* | 1.83387 | 0.001 | 1833.87 | UP |
| *CES2* | 453.75674 | 0.255181 | 1778.1761 | UP |
| *LOC101120476* | 1.775138 | 0.001 | 1775.138 | UP |
| *LOC105615807* | 1.583146 | 0.001 | 1583.146 | UP |
| *LOC101109475* | 1.5360383 | 0.001 | 1536.0383 | UP |
| *LOC101102291* | 1.527983 | 0.001 | 1527.983 | UP |
| *LOC106991725* | 1.345085 | 0.001 | 1345.085 | UP |
| *ENPP3* | 142.69919 | 0.107065 | 1332.8276 | UP |
| *SLC9A3* | 474.46475 | 0.381049 | 1245.1542 | UP |
| *LOC101121777* | 69.381836 | 0.0617623 | 1123.3687 | UP |
| *AKR1D1* | 405.96985 | 0.422032 | 961.94092 | UP |
| *SLC5A8* | 23.881535 | 0.0312024 | 765.37493 | UP |
| *KYNU* | 16.942459 | 0.023661 | 716.05 | UP |
| *SLC14A1* | 297.22721 | 0.503768 | 590.00812 | UP |
| *AQP5* | 327.50834 | 0.558485 | 586.4228 | UP |
| *APOBEC1* | 87.081261 | 0.154748 | 562.72948 | UP |
| *ST8SIA5* | 24.017414 | 0.0491597 | 488.559 | UP |
| *NOS2* | 40.516427 | 0.113414 | 357.24361 | UP |
| *PGLYRP2* | 37.726051 | 0.112852 | 334.2967 | UP |
| *ADGRF1* | 9.972401 | 0.0312011 | 319.61697 | UP |
| *CYP11A1* | 43.035049 | 0.146662 | 293.43013 | UP |
| *ALOX15B* | 91.782303 | 0.318433 | 288.23113 | UP |
| *PADI1* | 524.37415 | 1.83873 | 285.18279 | UP |
| *LOC101103238* | 72.687386 | 0.284193 | 255.76769 | UP |
| *GAP43* | 58.035931 | 0.259051 | 224.03284 | UP |
| *HAL* | 18.38686 | 0.0833092 | 220.70624 | UP |
| *MUC20* | 25.970194 | 0.120507 | 215.50776 | UP |
| *ADH1C* | 251.61824 | 1.18117 | 213.02458 | UP |
| *NXPE2* | 69.019848 | 0.345316 | 199.87446 | UP |
| *HCN2* | 20.325872 | 0.132458 | 153.45145 | UP |
| *TLL2* | 9.4796533 | 0.0618937 | 153.16023 | UP |
| *LOC101105864* | 305.74728 | 2.03272 | 150.41289 | UP |
| *CAV3* | 31.425636 | 0.220915 | 142.25216 | UP |
| *LOC101117044* | 122.21789 | 0.932664 | 131.04171 | UP |
| *DUOXA2* | 198.8734 | 1.90818 | 104.22151 | UP |
| *SLC4A9* | 35.489193 | 0.355289 | 99.88824 | UP |
| *UPK1B* | 116.92923 | 1.24974 | 93.562845 | UP |
| *PATL2* | 9.671702 | 0.10435 | 92.685213 | UP |
| *IL36B* | 20.29016 | 0.239877 | 84.585682 | UP |
| *DHRS9* | 44.312925 | 0.552439 | 80.213246 | UP |

| | | | | |
|---|---|---|---|---|
| *COLQ* | 8.3823298 | 0.105784 | 79.240054 | UP |
| *PRSS48* | 13.578557 | 0.175345 | 77.439089 | UP |
| *DAPL1* | 221.58798 | 3.04582 | 72.751503 | UP |
| *TNNT1* | 39.863798 | 0.561781 | 70.959676 | UP |
| *WSCD2* | 14.633092 | 0.214665 | 68.167108 | UP |
| *KCNG3* | 5.450149 | 0.0806443 | 67.582569 | UP |
| *HSD17B13* | 573.1944 | 8.64125 | 66.332347 | UP |
| *LOC101113369* | 376.17856 | 5.97321 | 62.977621 | UP |
| *HSBP1L1* | 24.859006 | 0.45318 | 54.854597 | UP |
| *CLDN3* | 11.393651 | 0.215345 | 52.908825 | UP |
| *NOXO1* | 50.31762 | 0.984273 | 51.121609 | UP |
| *KRT35* | 104.67263 | 2.08632 | 50.170935 | UP |
| *SYT8* | 8.752247 | 0.186769 | 46.861347 | UP |
| *AHSG* | 195.67271 | 4.20157 | 46.571333 | UP |
| *PRSS27* | 508.34326 | 11.9364 | 42.587653 | UP |
| *HSD17B6* | 25.585267 | 0.608613 | 42.038647 | UP |
| *LMO3* | 15.94971 | 0.389392 | 40.960549 | UP |
| *SERPINB10* | 186.46059 | 4.59679 | 40.563216 | UP |
| *TFF1* | 60.500359 | 1.49514 | 40.464678 | UP |
| *CDKL4* | 7.4574694 | 0.185013 | 40.307813 | UP |
| *GCNT3* | 222.82629 | 5.56397 | 40.048076 | UP |
| *LOC101112671* | 27.417829 | 0.711735 | 38.522525 | UP |
| *LIX1* | 16.506363 | 0.435207 | 37.927614 | UP |
| *MUC4* | 14.166592 | 0.378134 | 37.464476 | UP |
| *MUC1* | 44.921982 | 1.21103 | 37.094029 | UP |
| *FAM69C* | 53.599319 | 1.52461 | 35.156085 | UP |
| *PRSS35* | 59.16396 | 1.73305 | 34.138634 | UP |
| *DUSP27* | 30.648546 | 0.980912 | 31.24495 | UP |
| *TTLL10* | 10.742294 | 0.352866 | 30.442984 | UP |
| *NPR3* | 20.357536 | 0.69093 | 29.463963 | UP |
| *CCL20* | 67.358562 | 2.34706 | 28.699122 | UP |
| *SLC6A14* | 145.07994 | 5.53319 | 26.219945 | UP |
| *TMPRSS11A* | 90.160489 | 3.56615 | 25.282304 | UP |
| *ANKRD24* | 118.34734 | 4.78033 | 24.757148 | UP |
| *HAP1* | 9.502826 | 0.392372 | 24.21892 | UP |
| *TRIM71* | 5.684439 | 0.236394 | 24.046461 | UP |
| *LOC101114973* | 40.096695 | 1.70907 | 23.461119 | UP |
| *KCNG2* | 5.692732 | 0.246249 | 23.117787 | UP |
| *TACR1* | 3.084111 | 0.139691 | 22.078094 | UP |
| *STYK1* | 16.762373 | 0.759497 | 22.070361 | UP |
| *SDK2* | 5.660719 | 0.259939 | 21.777105 | UP |
| *ATP12A* | 198.99991 | 9.21291 | 21.600114 | UP |

| | | | | |
|---|---|---|---|---|
| *WDR66* | 9.769868 | 0.462412 | 21.128059 | UP |
| *ADAMTSL2* | 12.561535 | 0.613372 | 20.479472 | UP |
| *ARG1* | 87.806265 | 4.52556 | 19.402298 | UP |
| *TGM3* | 1612.35 | 84.0667 | 19.179413 | UP |
| *SCGB1C1* | 78.67699 | 4.1597 | 18.914102 | UP |
| *ACER1* | 51.264503 | 2.81016 | 18.242557 | UP |
| *TCN1* | 147.86495 | 8.11064 | 18.230984 | UP |
| *PAX9* | 85.438232 | 32.8803 | 2.5984627 | UP |
| *ISLR2* | 13.861473 | 0.786461 | 17.625124 | UP |
| *PRSS22* | 139.84729 | 7.96451 | 17.558807 | UP |
| *RYR3* | 6.313796 | 0.383915 | 16.445817 | UP |
| *FOSL1* | 14.408758 | 0.941384 | 15.30593 | UP |
| *DUOX2* | 134.68902 | 8.87378 | 15.178315 | UP |
| *IL26* | 2.429616 | 0.167568 | 14.499284 | UP |
| *PABPC1L* | 6.4797514 | 0.487187 | 13.300337 | UP |
| *TGM5* | 31.36367 | 2.39913 | 13.072935 | UP |
| *C12H1orf167* | 5.137315 | 0.396574 | 12.954241 | UP |
| *NPTXR* | 13.982965 | 1.08789 | 12.853289 | UP |
| *GLYCTK* | 46.036873 | 3.59286 | 12.813433 | UP |
| *COL17A1* | 699.55652 | 56.0719 | 12.476062 | UP |
| *ISG20* | 14.940276 | 1.21248 | 12.32208 | UP |
| *TRPC4* | 5.992506 | 0.492901 | 12.157626 | UP |
| *HMGCS2* | 1739.5532 | 145.036 | 11.993941 | UP |
| *NOD2* | 16.351261 | 1.37817 | 11.864473 | UP |
| *XKR6* | 3.034569 | 0.257843 | 11.769057 | UP |
| *CEMIP* | 59.927315 | 5.11767 | 11.709883 | UP |
| *WFIKKN2* | 27.736105 | 2.489 | 11.143473 | UP |
| *HDC* | 7.606122 | 0.710336 | 10.707781 | UP |
| *LOC101116795* | 72.79969 | 6.96937 | 10.445663 | UP |
| *DAB1* | 9.0204842 | 0.864142 | 10.43866 | UP |
| *IL17F* | 13.478943 | 1.29425 | 10.414482 | UP |
| *NPTX1* | 32.253757 | 3.43777 | 9.3821742 | UP |
| *C24H16orf89* | 21.518782 | 2.31928 | 9.2782165 | UP |
| *OTUB2* | 51.870735 | 5.59108 | 9.2774088 | UP |
| *MACC1* | 27.318498 | 3.14652 | 8.6821308 | UP |
| *RAB19* | 19.374962 | 2.33332 | 8.3036026 | UP |
| *FGF5* | 8.2021629 | 1.00049 | 8.1981458 | UP |
| *COL24A1* | 5.5042624 | 0.700998 | 7.8520372 | UP |
| *CLCA2* | 529.25873 | 68.0284 | 7.7799673 | UP |
| *TMEM95* | 5.0830349 | 0.65705 | 7.7361463 | UP |
| *KRT17* | 10910.678 | 1418.74 | 7.6903997 | UP |
| *C3H9orf172* | 3.506388 | 0.457804 | 7.6591467 | UP |

| | | | | |
|---|---|---|---|---|
| *PRR19* | 28.879236 | 3.84085 | 7.51897 | UP |
| *KLHL30* | 7.828433 | 1.08373 | 7.2236009 | UP |
| *GSDMA* | 32.486237 | 4.57024 | 7.1082125 | UP |
| *SCN9A* | 5.0866922 | 0.719333 | 7.0714012 | UP |
| *HSD17B2* | 77.46463 | 11.3814 | 6.8062479 | UP |
| *CLDN23* | 18.111824 | 2.66955 | 6.7845982 | UP |
| *LOC101106609* | 6.427092 | 0.949223 | 6.7708979 | UP |
| *IL22* | 2.1529694 | 0.324966 | 6.6252143 | UP |
| *ZNF662* | 3.911326 | 0.604856 | 6.4665408 | UP |
| *TMEM171* | 42.942453 | 6.6906 | 6.4183262 | UP |
| *TMPRSS4* | 166.54516 | 26.3746 | 6.3146042 | UP |
| *ENDOU* | 602.04717 | 96.8987 | 6.2131604 | UP |
| *ATG9B* | 102.61218 | 16.8227 | 6.0996263 | UP |
| *LOC101102014* | 46.602486 | 7.75254 | 6.0112539 | UP |
| *LOC101116267* | 118.30653 | 20.1225 | 5.8793159 | UP |
| *CARTPT* | 19.205036 | 3.26976 | 5.8735308 | UP |
| *LOC101106871* | 283.29346 | 48.5574 | 5.8341974 | UP |
| *PLA2G4E* | 60.229904 | 10.3818 | 5.8014895 | UP |
| *DCST2* | 6.514465 | 1.13553 | 5.7369378 | UP |
| *LOC101123254* | 135.57785 | 23.9799 | 5.6538122 | UP |
| *KRT78* | 3323.9919 | 595.64 | 5.5805385 | UP |
| *SLC28A3* | 24.741346 | 4.61379 | 5.3624777 | UP |
| *BBOX1* | 17.741981 | 3.32118 | 5.3420714 | UP |
| *PDE6A* | 7.7405146 | 1.52746 | 5.0675727 | UP |
| *ATP7B* | 7.092519 | 1.4466 | 4.9028888 | UP |
| *SLC26A9* | 21.821831 | 4.53372 | 4.8132287 | UP |
| *SMYD1* | 226.03348 | 49.4949 | 4.5668034 | UP |
| *MAP3K7CL* | 5.9383955 | 1.33431 | 4.4505366 | UP |
| *MUC15* | 60.834198 | 13.9031 | 4.3755852 | UP |
| *FAM83C* | 37.643192 | 8.64313 | 4.3552731 | UP |
| *RAX2* | 7.029022 | 1.644 | 4.2755608 | UP |
| *TRPV3* | 24.714767 | 5.7938 | 4.2657267 | UP |
| *MFAP3L* | 10.280406 | 2.48604 | 4.1352537 | UP |
| *C14H19orf33* | 374.06604 | 789.12 | 0.474029349 | DOWN |
| *CTNNBIP1* | 6.857236984 | 78.1315 | 0.087765331 | DOWN |
| *LOC101110243* | 4.53059 | 156.508 | 0.028947977 | DOWN |
| *CAPNS2* | 7.457073 | 213.295 | 0.034961312 | DOWN |
| *LOC105615767* | 21.144239 | 273.063 | 0.077433556 | DOWN |
| *LOC101122488* | 6.633393 | 123.124 | 0.053875711 | DOWN |
| *LOC105610290* | 2.869406 | 64.8738 | 0.044230583 | DOWN |
| *FXYD3* | 249.3502923 | 1400.82 | 0.178003093 | DOWN |
| *CEBPA* | 8.4515 | 204.101 | 0.04140842 | DOWN |

| | | | | |
|---|---|---|---|---|
| *CDCA4* | 3.161831 | 31.1532 | 0.101492977 | DOWN |
| *S1PR5* | 6.356143 | 30.3071 | 0.209724553 | DOWN |
| *CNFN* | 251.5022392 | 1112.56 | 0.226057237 | DOWN |
| *RHOV* | 8.617823 | 302.177 | 0.028519123 | DOWN |
| *RBBP8NL* | 8.291495 | 81.4697 | 0.101773972 | DOWN |
| *KLK10* | 159.4083101 | 650.483 | 0.245061455 | DOWN |
| *SLC35A4* | 4.440803 | 36.1474 | 0.122852626 | DOWN |
| *KLHDC9* | 2.579979 | 11.2416 | 0.229502829 | DOWN |
| *HOXC5* | 9.316009 | 22.398 | 0.415930396 | DOWN |
| *LOC101102814* | 1.603957 | 7.80787 | 0.205428241 | DOWN |
| *RPP25* | 3.286834 | 9.79964 | 0.335403545 | DOWN |
| *EFNA3* | 10.1716334 | 264.542 | 0.038449975 | DOWN |
| *NKPD1* | 13.798231 | 43.5404 | 0.31690639 | DOWN |
| *TMEM154* | 7.437961617 | 32.2661 | 0.230519388 | DOWN |
| *STX19* | 5.235295029 | 10.7123 | 0.488718112 | DOWN |
| *ARHGEF4* | 21.65247324 | 88.8711 | 0.243639082 | DOWN |
| *PPARG* | 17.36245546 | 46.4012 | 0.374181173 | DOWN |
| *LOC101118459* | 3471.791748 | 14797.2 | 0.234624912 | DOWN |
| *CCDC64B* | 37.079433 | 134.577 | 0.275525781 | DOWN |
| *DLK2* | 35.953114 | 78.2719 | 0.45933616 | DOWN |
| *FKBPL* | 2.495618 | 8.74435 | 0.285397771 | DOWN |
| *STAC* | 6.438572314 | 16.2241 | 0.396852356 | DOWN |
| *HOXC6* | 17.842428 | 117.575 | 0.151753587 | DOWN |
| *EDARADD* | 3.391306 | 8.85984 | 0.382772827 | DOWN |
| *IRX5* | 5.887524 | 24.222 | 0.243065147 | DOWN |
| *LOC101117431* | 17621.99023 | 39187.1 | 0.449688551 | DOWN |
| *PITX1* | 275.995056 | 1807.42 | 0.152701119 | DOWN |
| *ZNF296* | 10.39789 | 34.0096 | 0.305733969 | DOWN |
| *CDKN2B* | 11.517081 | 34.797 | 0.330979136 | DOWN |
| *C1QTNF7* | 3.477997762 | 27.4566 | 0.126672558 | DOWN |
| *FAM26E* | 5.293519 | 16.2912 | 0.32493119 | DOWN |
| *LOC105610993* | 3.524554 | 11.8503 | 0.297423188 | DOWN |
| *TMEM177* | 2.520193 | 13.6476 | 0.184661992 | DOWN |
| *HOXC8* | 19.439707 | 52.4905 | 0.370347149 | DOWN |
| *EPS8L1* | 57.413067 | 122.089 | 0.470255854 | DOWN |
| *SH3RF2* | 56.10287183 | 126.086 | 0.444957187 | DOWN |
| *BARX1* | 23.421793 | 92.6771 | 0.252724708 | DOWN |
| *LOC101104157* | 7.393049 | 229.143 | 0.032263909 | DOWN |
| *SLC52A3* | 15.354591 | 48.796 | 0.314669051 | DOWN |
| *VIPR2* | 3.701156 | 9.37113 | 0.39495301 | DOWN |
| *LYPD3* | 735.259216 | 3408.74 | 0.215698239 | DOWN |
| *ANKRD35* | 25.870457 | 95.5896 | 0.270640917 | DOWN |

| | | | | |
|---|---|---|---|---|
| *TP53INP2* | 20.50809788 | 43.9218 | 0.466922983 | DOWN |
| *SOSTDC1* | 21.898233 | 62.6186 | 0.349708122 | DOWN |
| *TMEM61* | 8.461056 | 20.4739 | 0.41326059 | DOWN |
| *SLC25A21* | 9.535874702 | 20.8133 | 0.458162555 | DOWN |
| *STBD1* | 8.091275 | 20.9951 | 0.385388734 | DOWN |
| *PCSK4* | 4.790973 | 22.403 | 0.213854082 | DOWN |
| *KLK12* | 155.4306637 | 447.086 | 0.347652719 | DOWN |
| *HS3ST2* | 5.253508 | 29.8009 | 0.176286891 | DOWN |
| *KRT4* | 16500.06641 | 41870.6 | 0.394072844 | DOWN |
| *CHRM3* | 3.677505005 | 9.34015 | 0.393730829 | DOWN |
| *PLCH2* | 9.073736 | 49.1688 | 0.184542555 | DOWN |
| *ADRA2C* | 5.837276 | 26.2891 | 0.222041683 | DOWN |
| *CHRNA3* | 7.698875 | 16.4026 | 0.469369185 | DOWN |
| *ARL10* | 2.587248 | 5.62239 | 0.460168718 | DOWN |
| *RNF223* | 5.868876 | 46.7074 | 0.125651952 | DOWN |
| *LTB4R* | 5.932357492 | 13.4759 | 0.440219762 | DOWN |
| *UGT1A1* | 111.218872 | 490.916 | 0.226553773 | DOWN |
| *PIGY* | 8.548679 | 66.5011 | 0.128549438 | DOWN |
| *PCP4L1* | 14.814951 | 36.337 | 0.4077098 | DOWN |
| *MGAT3* | 15.141333 | 44.3214 | 0.341625783 | DOWN |
| *DUOXA1* | 19.55050017 | 152.438 | 0.128252143 | DOWN |
| *LOC101121908* | 21.94349203 | 69.1422 | 0.31736757 | DOWN |
| *LOC101103771* | 9437.467368 | 55972.9 | 0.168607797 | DOWN |
| *BARX2* | 62.958668 | 336.837 | 0.186911379 | DOWN |
| *TRIM29* | 223.115372 | 1145.62 | 0.19475513 | DOWN |
| *KRTDAP* | 1611.144046 | 3769.67 | 0.427396575 | DOWN |
| *S100A9* | 4921.76416 | 26740.6 | 0.184055861 | DOWN |
| *S100A8* | 8764.375977 | 38088.5 | 0.230105569 | DOWN |

**Supplementary Table 8 |** The gene expression and fold change of DEGs between the rumen and the FC stomach of cetaceans. The UP represents the gene is upregulated in the rumen; the DOWN represents the gene is downregulated in the rumen when compared to the FC stomach of cetaceans.

| **Official name** | **Rumen of Sheep** | **The FC stomach of cetaceans** | **Fold change of Sheep/Cetaceans** | **Rumen** |
|---|---|---|---|---|
| *C1QTNF7* | 3.477998 | 1.692016 | 2.05553521 | UP |
| *FAM26E* | 5.293519 | 2.531592 | 2.09098425 | UP |
| *KLK9* | 37.22844 | 17.72658 | 2.10014809 | UP |
| *TFAP2A* | 17.00198 | 7.986775 | 2.12876698 | UP |
| *GJB6* | 129.7337 | 60.37995 | 2.14862225 | UP |
| *LOC105610993* | 3.524554 | 1.616964 | 2.1797356 | UP |
| *CERS3* | 16.58827 | 7.530555 | 2.20279519 | UP |
| *LOC101103862* | 707.795 | 313.5804 | 2.25714072 | UP |
| *VSTM2L* | 21.24554 | 9.238019 | 2.2997936 | UP |
| *TMEM177* | 2.520193 | 1.07964 | 2.33428914 | UP |
| *SRPK3* | 10.64524 | 4.449088 | 2.39267899 | UP |
| *FAM83B* | 18.08906 | 7.553164 | 2.39489835 | UP |
| *GRHL1* | 60.32241 | 24.6841 | 2.4437765 | UP |
| *ZACN* | 7.692018 | 3.122662 | 2.46328869 | UP |
| *LOC101121414* | 6.582892 | 2.640858 | 2.49270953 | UP |
| *NPAS1* | 6.030153 | 2.385069 | 2.52829289 | UP |
| *CD70* | 3.024992 | 1.186656 | 2.54917348 | UP |
| *GCC1* | 3.954875 | 1.542238 | 2.56437398 | UP |
| *HOXC8* | 19.43971 | 7.551766 | 2.57419351 | UP |
| *CD164L2* | 14.75401 | 5.716718 | 2.58085291 | UP |
| *SH3RF3* | 4.854718 | 1.860044 | 2.6100017 | UP |
| *AHNAK2* | 9.03539 | 3.399268 | 2.65803991 | UP |
| *EPS8L1* | 57.41307 | 21.53972 | 2.66545107 | UP |
| *SH3RF2* | 56.10287 | 20.91143 | 2.68288018 | UP |
| *BARX1* | 23.42179 | 8.642412 | 2.7100991 | UP |
| *GRIK5* | 7.673132 | 2.711081 | 2.83028465 | UP |
| *CCDC136* | 9.486832 | 3.323746 | 2.85425927 | UP |
| *LOC105608435* | 10.02089 | 3.499694 | 2.86336034 | UP |
| *NEURL2* | 7.459035 | 2.575146 | 2.8965484 | UP |
| *CBLC* | 32.60547 | 10.88366 | 2.99581951 | UP |
| *LOC101104157* | 7.393049 | 2.443104 | 3.02608853 | UP |
| *CDA* | 65.93362 | 21.52013 | 3.06381099 | UP |
| *LOC106991601* | 4.604927 | 1.466479 | 3.14012501 | UP |
| *AHRR* | 8.042508 | 2.530831 | 3.17781314 | UP |

| | | | | |
|---|---|---|---|---|
| *VTCN1* | 7.740344 | 2.411588 | 3.20964611 | UP |
| *COL7A1* | 146.0332 | 45.1793 | 3.23230283 | UP |
| *KCNQ4* | 7.220798 | 2.218674 | 3.25455565 | UP |
| *B4GALNT3* | 60.21916 | 17.62143 | 3.41738261 | UP |
| *ARL4A* | 21.59941 | 6.2647 | 3.44779657 | UP |
| *ZNF750* | 63.1736 | 17.65586 | 3.57805258 | UP |
| *L3MBTL1* | 2.806673 | 0.77612 | 3.61628743 | UP |
| *FAT2* | 13.34394 | 3.636741 | 3.66920328 | UP |
| *ZBTB7C* | 10.92888 | 2.975055 | 3.67350692 | UP |
| *HAPLN3* | 11.57171 | 2.978164 | 3.88551724 | UP |
| *C12H1orf116* | 55.69708 | 14.2587 | 3.90618119 | UP |
| *DSC3* | 93.86417 | 23.66929 | 3.96565151 | UP |
| *LOC101112304* | 15279.52 | 0.001 | 15279522.5 | UP |
| *ANKRD22* | 22.34352 | 5.219203 | 4.28102181 | UP |
| *TP63* | 57.99331 | 13.51653 | 4.29054777 | UP |
| *GRHL3* | 37.02918 | 8.564453 | 4.32358903 | UP |
| *LOC101102714* | 10660.87 | 0.001 | 10660870.1 | UP |
| *EVPL* | 186.5052 | 42.39204 | 4.3995328 | UP |
| *MSX1* | 19.52972 | 4.374823 | 4.46411546 | UP |
| *AIM1L* | 40.39148 | 9.039898 | 4.46813437 | UP |
| *LOC105613035* | 8160.179 | 0.001 | 8160179.2 | UP |
| *LOC101113601* | 7766.591 | 0.151575 | 51239.3959 | UP |
| *HAS3* | 11.87929 | 2.57071 | 4.62101532 | UP |
| *A2ML1* | 6078.86 | 0.001 | 6078859.89 | UP |
| *SLC52A3* | 15.35459 | 3.178597 | 4.8306195 | UP |
| *CCR10* | 2.714588 | 0.557358 | 4.87045669 | UP |
| *C12H1orf106* | 29.01655 | 5.813163 | 4.99152596 | UP |
| *LOC101110777* | 5062.031 | 0.001 | 5062030.76 | UP |
| *VIPR2* | 3.701156 | 0.722563 | 5.12226062 | UP |
| *LOC101102105* | 4067.69 | 0.001 | 4067690.19 | UP |
| *LYPD3* | 735.2592 | 138.7923 | 5.29755174 | UP |
| *STEAP3* | 7.608882 | 1.416484 | 5.37166865 | UP |
| *HS3ST6* | 69.9433 | 12.95506 | 5.39891775 | UP |
| *GPR1* | 6.231207 | 1.14351 | 5.44919266 | UP |
| *ANKRD35* | 25.87046 | 4.708093 | 5.49489127 | UP |
| *LOC101111178* | 3783.106 | 0.001 | 3783105.71 | UP |
| *IRS2* | 3.095991 | 0.548846 | 5.6409102 | UP |
| *SNX31* | 15.88443 | 2.652551 | 5.98835895 | UP |
| *LOC101119041* | 31.11536 | 5.161098 | 6.02882487 | UP |
| *IL1RN* | 56.84334 | 9.399039 | 6.04778252 | UP |
| *PRD-SPRRII* | 3708.906 | 0.001 | 3708906.49 | UP |
| *ANKRD37* | 12.20403 | 1.993658 | 6.12142528 | UP |

| | | | | |
|---|---|---|---|---|
| *TP53INP2* | 20.5081 | 3.329977 | 6.15862987 | UP |
| *LOC101104114* | 3006.677 | 0.001 | 3006677.25 | UP |
| *MN1* | 6.998863 | 1.089726 | 6.42258972 | UP |
| *LOC105615728* | 2962.506 | 0.001 | 2962506.1 | UP |
| *CRISP3* | 1638.922 | 0.001 | 1638922.36 | UP |
| *RNF39* | 29.49555 | 4.197665 | 7.02665744 | UP |
| *SOSTDC1* | 21.89823 | 3.034231 | 7.21706192 | UP |
| *PKP1* | 288.4993 | 39.4666 | 7.30996119 | UP |
| *EPHA4* | 7.162824 | 0.956451 | 7.48896472 | UP |
| *IVL* | 1519.637 | 0.001 | 1519636.65 | UP |
| *LOC101107119* | 1239.146 | 0.001 | 1239145.67 | UP |
| *SOX2* | 132.4701 | 15.89663 | 8.33321861 | UP |
| *VSIG2* | 15.63402 | 1.870931 | 8.35627717 | UP |
| *TMEM61* | 8.461056 | 0.978452 | 8.64738996 | UP |
| *ACACB* | 12.15978 | 1.378151 | 8.82325304 | UP |
| *CCDC183* | 17.82192 | 2.009396 | 8.86929406 | UP |
| *LOC101121219* | 1217.002 | 0.001 | 1217002.46 | UP |
| *SLC25A21* | 9.535875 | 1.036492 | 9.20014308 | UP |
| *PLA2G4F* | 43.10029 | 4.602343 | 9.36485842 | UP |
| *LYPD2* | 1000.9 | 0.001 | 1000899.96 | UP |
| *SDCBP2* | 94.37186 | 9.679737 | 9.74942367 | UP |
| *TRIM62* | 12.53244 | 1.280977 | 9.78349703 | UP |
| *C7H14orf169* | 17.8153 | 1.763522 | 10.1021149 | UP |
| *BFSP1* | 5.809955 | 0.5681 | 10.2269935 | UP |
| *STBD1* | 8.091275 | 0.783161 | 10.3315602 | UP |
| *PROM2* | 124.2034 | 11.86801 | 10.4653883 | UP |
| *LINGO1* | 4.663729 | 0.443876 | 10.5068285 | UP |
| *HTR2A* | 3.628845 | 0.34297 | 10.580636 | UP |
| *CPA6* | 4.304087 | 0.392305 | 10.9712774 | UP |
| *EDAR* | 12.36471 | 1.123908 | 11.0015319 | UP |
| *EREG* | 12.7423 | 1.138663 | 11.1905797 | UP |
| *PROSER2* | 8.397456 | 0.70605 | 11.8935791 | UP |
| *PCSK4* | 4.790973 | 0.396191 | 12.0925841 | UP |
| *KRT79* | 955.0491 | 0.001 | 955049.133 | UP |
| *TMPRSS13* | 67.04128 | 5.409249 | 12.3938245 | UP |
| *KLK12* | 155.4307 | 12.47755 | 12.4568305 | UP |
| *HS3ST2* | 5.253508 | 0.411022 | 12.7815737 | UP |
| *TMEM45B* | 53.09155 | 4.138809 | 12.8277381 | UP |
| *LOC101116141* | 880.8626 | 0.001 | 880862.63 | UP |
| *LOC101106610* | 720.6295 | 0.001 | 720629.498 | UP |
| *LOC101122351* | 715.2217 | 0.001 | 715221.68 | UP |
| *LOC101104808* | 698.7977 | 0.001 | 698797.729 | UP |

| | | | | |
|---|---|---|---|---|
| *LRAT* | 5.558094 | 0.385444 | 14.4199786 | UP |
| *LOC101111992* | 621.7982 | 0.001 | 621798.157 | UP |
| *LOC105604529* | 523.9991 | 0.001 | 523999.084 | UP |
| *LOC101122545* | 89.71052 | 5.822931 | 15.4064196 | UP |
| *SLC26A3* | 457.0272 | 0.001 | 457027.234 | UP |
| *KCNK9* | 4.23823 | 0.273933 | 15.4717759 | UP |
| *ZNF385D* | 8.599665 | 0.552262 | 15.5717123 | UP |
| *SBD2* | 452.695 | 0.001 | 452694.977 | UP |
| *NXNL2* | 4.431918 | 0.274567 | 16.141481 | UP |
| *UGT1A9* | 436.0501 | 0.001 | 436050.11 | UP |
| *LOC101108147* | 414.4206 | 0.001 | 414420.624 | UP |
| *LOC101103612* | 403.2927 | 0.001 | 403292.725 | UP |
| *KRT4* | 16500.07 | 885.6228 | 18.6310316 | UP |
| *LOC101120889* | 14.05127 | 0.751768 | 18.6909765 | UP |
| *LOC106991749* | 8.373987 | 0.440137 | 19.0258555 | UP |
| *IL36A* | 323.7675 | 0.001 | 323767.474 | UP |
| *LOC101119393* | 313.4223 | 0.001 | 313422.254 | UP |
| *MST1R* | 21.40555 | 1.018824 | 21.0100547 | UP |
| *FZD10* | 13.88646 | 0.632165 | 21.9665087 | UP |
| *LOC101115964* | 303.13 | 0.001 | 303130.005 | UP |
| *OTX1* | 8.534991 | 0.357412 | 23.8799548 | UP |
| *PRDM6* | 7.007299 | 0.29292 | 23.9222279 | UP |
| *ACTA1* | 279.391 | 0.001 | 279390.991 | UP |
| *DENND2C* | 40.09808 | 1.638447 | 24.4732209 | UP |
| *LOC101107809* | 260.7116 | 0.001 | 260711.578 | UP |
| *PNCK* | 38.86594 | 1.465353 | 26.5232612 | UP |
| *PI3* | 258.7939 | 0.400744 | 645.783553 | UP |
| *LOC101115172* | 242.4059 | 0.001 | 242405.93 | UP |
| *CHRM3* | 3.677505 | 0.121252 | 30.3294379 | UP |
| *LOC101112555* | 239.5526 | 0.001 | 239552.612 | UP |
| *STC2* | 12.99392 | 0.40116 | 32.390859 | UP |
| *SERPINB12* | 205.4519 | 0.001 | 205451.87 | UP |
| *LOC105614373* | 203.7187 | 0.001 | 203718.704 | UP |
| *KRT23* | 196.9857 | 0.001 | 196985.669 | UP |
| *DMRT2* | 5.497377 | 0.16117 | 34.1091829 | UP |
| *CPAMD8* | 5.532654 | 0.161167 | 34.3287025 | UP |
| *DSG3* | 623.4412 | 18.12918 | 34.3888206 | UP |
| *JPH2* | 85.95128 | 2.489614 | 34.5239378 | UP |
| *GGT6* | 25.65006 | 0.729627 | 35.1550295 | UP |
| *LIPH* | 14.19718 | 0.39786 | 35.683859 | UP |
| *LOC101121267* | 184.4823 | 0.001 | 184482.315 | UP |
| *FBXL22* | 9.989536 | 0.26949 | 37.0682994 | UP |

| | | | | |
|---|---|---|---|---|
| *OVOL1* | 42.47857 | 1.088158 | 39.0371334 | UP |
| *LOC101117764* | 182.2246 | 0.001 | 182224.609 | UP |
| *LOC101120106* | 164.7707 | 0.001 | 164770.676 | UP |
| *APOBEC3Z1* | 146.4274 | 0.001 | 146427.383 | UP |
| *LOC101117163* | 138.395 | 0.001 | 138394.951 | UP |
| *SCEL* | 134.9509 | 0.001 | 134950.933 | UP |
| *C1H2orf54* | 133.0843 | 0.001 | 133084.259 | UP |
| *VILL* | 39.62866 | 0.738179 | 53.6843476 | UP |
| *LOC443322* | 122.7522 | 0.001 | 122752.208 | UP |
| *LOC101102344* | 119.8675 | 0.001 | 119867.455 | UP |
| *SSC4D* | 7.227684 | 0.119127 | 60.6720895 | UP |
| *LOC101108071* | 110.2744 | 0.001 | 110274.368 | UP |
| *PLA2G2F* | 107.2031 | 0.001 | 107203.15 | UP |
| *KLHL41* | 57.73458 | 0.930301 | 62.0601042 | UP |
| *IL19* | 99.40583 | 7.251646 | 13.7080368 | UP |
| *TOB2* | 24.82166 | 0.372757 | 66.5893732 | UP |
| *MTUS2* | 8.910978 | 0.133104 | 66.9472842 | UP |
| *MYOCD* | 9.687936 | 0.143049 | 67.7245979 | UP |
| *RBPMS2* | 95.17597 | 0.001 | 95175.972 | UP |
| *KRT80* | 91.30562 | 0.001 | 91305.618 | UP |
| *LOC101106641* | 90.53042 | 0.001 | 90530.4173 | UP |
| *RPTN* | 80.38301 | 0.001 | 80383.011 | UP |
| *ZNF185* | 75.69835 | 0.001 | 75698.3523 | UP |
| *LOC101118712* | 75.6394 | 0.001 | 75639.404 | UP |
| *LOC106991113* | 73.21864 | 0.001 | 73218.643 | UP |
| *ATP6V1C2* | 71.62398 | 0.001 | 71623.978 | UP |
| *SLITRK5* | 7.129467 | 0.084485 | 84.3873705 | UP |
| *KRT36* | 9886.455 | 1.395109 | 7086.51086 | UP |
| *KRT34* | 64.54622 | 0.001 | 64546.219 | UP |
| *RBM24* | 21.87512 | 0.245138 | 89.2359357 | UP |
| *FUT5* | 57.32155 | 0.001 | 57321.5545 | UP |
| *SERPINB13* | 57.01982 | 0.001 | 57019.82 | UP |
| *IL1R2* | 41.27016 | 0.418037 | 98.7236241 | UP |
| *LOC101102969* | 56.23896 | 0.001 | 56238.964 | UP |
| *PLCH2* | 9.073736 | 0.091189 | 99.504721 | UP |
| *SLURP1* | 53.91094 | 0.001 | 53910.942 | UP |
| *LOC101103602* | 51.17694 | 0.001 | 51176.943 | UP |
| *LOC101107619* | 48.29104 | 0.001 | 48291.035 | UP |
| *ANKRD9* | 46.08163 | 0.001 | 46081.627 | UP |
| *FGB* | 16.15834 | 0.125076 | 129.18819 | UP |
| *ADRA2C* | 5.837276 | 0.04324 | 134.997132 | UP |
| *CWH43* | 31.42029 | 0.202951 | 154.81711 | UP |

| | | | | |
|---|---|---|---|---|
| *LOC101115983* | 45.11112 | 0.001 | 45111.118 | UP |
| *SOX21* | 44.41419 | 0.001 | 44414.188 | UP |
| *TNFRSF6B* | 41.54738 | 8.651921 | 4.80209909 | UP |
| *LOC101105114* | 40.2042 | 0.001 | 40204.201 | UP |
| *MUC21* | 1014.247 | 5.82616 | 174.084959 | UP |
| *FAM127A* | 38.44585 | 0.001 | 38445.85 | UP |
| *LOC101116036* | 37.639 | 0.001 | 37639.004 | UP |
| *GPT* | 34.75334 | 0.001 | 34753.342 | UP |
| *LOC105602022* | 34.53469 | 0.001 | 34534.691 | UP |
| *CARD14* | 34.15244 | 0.001 | 34152.4424 | UP |
| *B4GALNT2* | 33.78363 | 0.001 | 33783.635 | UP |
| *ACTN2* | 46.42421 | 0.197489 | 235.072871 | UP |
| *LOC101114663* | 33.39572 | 0.001 | 33395.721 | UP |
| *FAM150A* | 32.41682 | 0.001 | 32416.817 | UP |
| *CHRNA3* | 7.698875 | 0.028647 | 268.754186 | UP |
| *LOC101114920* | 31.71809 | 0.001 | 31718.086 | UP |
| *ANKRD23* | 31.14351 | 0.001 | 31143.5123 | UP |
| *LOC101105876* | 30.3457 | 0.001 | 30345.701 | UP |
| *SLC5A1* | 75.42542 | 0.156051 | 483.33784 | UP |
| *CEACAM19* | 29.69046 | 0.001 | 29690.464 | UP |
| *CHST4* | 28.61247 | 0.001 | 28612.474 | UP |
| *KLC3* | 28.42285 | 0.001 | 28422.85 | UP |
| *LOC101107541* | 27.69426 | 0.001 | 27694.26 | UP |
| *LOC105616317* | 27.0355 | 0.001 | 27035.503 | UP |
| *LOC101103644* | 26.32515 | 0.001 | 26325.151 | UP |
| *ADGRF4* | 24.72151 | 0.001 | 24721.5053 | UP |
| *CDR1* | 23.52682 | 0.001 | 23526.815 | UP |
| *LYPD5* | 91.73491 | 0.09001 | 1019.16353 | UP |
| *LOC101106229* | 23.18036 | 0.001 | 23180.359 | UP |
| *HPSE2* | 15.14695 | 0.012467 | 1214.98912 | UP |
| *S100A5* | 22.64112 | 0.001 | 22641.1162 | UP |
| *LOC101113860* | 33.35526 | 0.026537 | 1256.93396 | UP |
| *POF1B* | 72.0788 | 0.057324 | 1257.39299 | UP |
| *P2RY2* | 22.63269 | 0.001 | 22632.69 | UP |
| *UGT1A4* | 21.08562 | 0.001 | 21085.617 | UP |
| *GAL3ST2* | 20.68874 | 0.001 | 20688.736 | UP |
| *ACPP* | 20.05565 | 0.001 | 20055.6542 | UP |
| *LOC106990121* | 19.69331 | 0.001 | 19693.3124 | UP |
| *THEM5* | 81.16286 | 0.050122 | 1619.30607 | UP |
| *LOC105616285* | 19.49552 | 0.001 | 19495.518 | UP |
| *SFTPC* | 19.14474 | 0.114403 | 167.344316 | UP |
| *SPINK9* | 18.49817 | 0.001 | 18498.169 | UP |

| | | | | |
|---|---|---|---|---|
| *PPP2R2C* | 17.98931 | 0.001 | 17989.3131 | UP |
| *LOC101107098* | 17.47176 | 0.001 | 17471.764 | UP |
| *GSDMC* | 17.38128 | 0.001 | 17381.283 | UP |
| *CAND2* | 16.8654 | 0.001 | 16865.402 | UP |
| *LOC101110524* | 16.52492 | 0.001 | 16524.918 | UP |
| *PGLYRP4* | 16.22951 | 0.001 | 16229.507 | UP |
| *LOC101113335* | 15.40575 | 0.001 | 15405.7451 | UP |
| *LOC105608915* | 14.53024 | 0.001 | 14530.236 | UP |
| *LOC105614885* | 13.98985 | 0.001 | 13989.855 | UP |
| *LOC105612196* | 13.94753 | 0.001 | 13947.5343 | UP |
| *POPDC2* | 50.65335 | 0.024239 | 2089.74578 | UP |
| *FCER2* | 13.90866 | 0.001 | 13908.6621 | UP |
| *LOC105614940* | 13.65509 | 0.001 | 13655.092 | UP |
| *STK32C* | 13.11965 | 0.001 | 13119.646 | UP |
| *LOC101104661* | 12.93321 | 0.001 | 12933.2127 | UP |
| *LOC105616478* | 12.21197 | 0.001 | 12211.966 | UP |
| *HOXB7* | 12.07503 | 0.001 | 12075.0348 | UP |
| *TRPM6* | 12.06659 | 0.001 | 12066.5851 | UP |
| *LOC101120775* | 12.00505 | 0.001 | 12005.045 | UP |
| *LTB4R2* | 11.96399 | 0.001 | 11963.988 | UP |
| *WNT7B* | 11.95372 | 0.001 | 11953.721 | UP |
| *ARL10* | 2.587248 | 0.001 | 2587.248 | UP |
| *HSPA1A* | 11.91347 | 0.001 | 11913.466 | UP |
| *SERPINB11* | 11.74515 | 0.001 | 11745.147 | UP |
| *LOC105604082* | 10.93128 | 0.001 | 10931.283 | UP |
| *KIF26A* | 10.66053 | 0.001 | 10660.532 | UP |
| *LOC105611786* | 10.54538 | 0.001 | 10545.382 | UP |
| *ADTRP* | 10.45416 | 0.001 | 10454.1605 | UP |
| *LOC101121036* | 10.1539 | 0.001 | 10153.899 | UP |
| *LOC105605155* | 9.557295 | 0.001 | 9557.295 | UP |
| *C13H10orf113* | 9.206914 | 0.001 | 9206.914 | UP |
| *LOC105610609* | 8.921119 | 0.001 | 8921.119 | UP |
| *IL17REL* | 8.722609 | 0.001 | 8722.60876 | UP |
| *ATP13A5* | 8.697449 | 0.001 | 8697.44925 | UP |
| *LOC105612090* | 8.549753 | 0.001 | 8549.753 | UP |
| *LOC101117587* | 8.327113 | 0.001 | 8327.113 | UP |
| *LRFN4* | 8.313799 | 0.001 | 8313.799 | UP |
| *A1BG* | 8.307885 | 0.001 | 8307.885 | UP |
| *LOC101117120* | 8.285954 | 0.001 | 8285.954 | UP |
| *LOC101105566* | 8.140181 | 0.001 | 8140.181 | UP |
| *ADGRF2* | 7.952637 | 0.001 | 7952.63725 | UP |
| *WDR38* | 3.383704 | 0.001 | 3383.70374 | UP |

| | | | | |
|---|---|---|---|---|
| *LOC101110696* | 7.895658 | 0.001 | 7895.658 | UP |
| *BHLHE23* | 7.758808 | 0.001 | 7758.808 | UP |
| *LOC105608606* | 7.647254 | 0.001 | 7647.254 | UP |
| *MCCD1* | 7.555327 | 0.001 | 7555.327 | UP |
| *LOC105606870* | 7.53356 | 0.001 | 7533.56009 | UP |
| *LOC101120435* | 7.258325 | 0.001 | 7258.325 | UP |
| *LOC105604589* | 7.234852 | 0.001 | 7234.852 | UP |
| *LOC106991452* | 6.686908 | 0.001 | 6686.908 | UP |
| *LOC101105174* | 6.561944 | 0.001 | 6561.944 | UP |
| *RASSF9* | 3.932756 | 0.001 | 3932.7562 | UP |
| *LOC106991450* | 6.491086 | 0.001 | 6491.086 | UP |
| *LOC101115778* | 6.490631 | 0.001 | 6490.631 | UP |
| *EMILIN3* | 3.974493 | 0.001 | 3974.493 | UP |
| *VAX2* | 6.132115 | 0.001 | 6132.115 | UP |
| *LOC105615508* | 5.752361 | 0.001 | 5752.361 | UP |
| *C1QL1* | 5.433838 | 0.001 | 5433.838 | UP |
| *LOC101104501* | 5.417474 | 0.001 | 5417.474 | UP |
| *KLF13* | 5.207161 | 0.001 | 5207.161 | UP |
| *POC1A* | 5.195816 | 0.001 | 5195.8163 | UP |
| *LOC101105258* | 5.052308 | 0.001 | 5052.308 | UP |
| *SH2D7* | 4.951905 | 0.09436 | 52.4788576 | UP |
| *LOC101122314* | 4.784063 | 0.001 | 4784.063 | UP |
| *LOC105602127* | 4.748694 | 0.001 | 4748.694 | UP |
| *LOC101120688* | 4.672022 | 0.001 | 4672.022 | UP |
| *LOC106990150* | 5.030138 | 0.001 | 5030.138 | UP |
| *LOC101111245* | 4.639668 | 0.001 | 4639.668 | UP |
| *LOC101104703* | 4.55153 | 0.001 | 4551.53 | UP |
| *LOC101116196* | 4.509582 | 0.001 | 4509.58228 | UP |
| *NKX2-2* | 4.422982 | 0.001 | 4422.982 | UP |
| *APELA* | 4.212973 | 0.049898 | 84.4319504 | UP |
| *SPATA12* | 4.126355 | 0.001 | 4126.355 | UP |
| *LOC105615865* | 4.101434 | 0.001 | 4101.434 | UP |
| *LOC105605056* | 4.023712 | 0.001 | 4023.712 | UP |
| *LOC101106977* | 3.967016 | 0.001 | 3967.016 | UP |
| *RNF223* | 5.868876 | 0.001 | 5868.876 | UP |
| *LTB4R* | 5.932357 | 0.001 | 5932.35749 | UP |
| *LOC101114497* | 3.944958 | 0.001 | 3944.958 | UP |
| *PLA2G3* | 6.110258 | 0.001 | 6110.258 | UP |
| *LOC105606498* | 3.920981 | 0.001 | 3920.981 | UP |
| *ARSE* | 3.907126 | 0.001 | 3907.12556 | UP |
| *LOC106990135* | 3.841791 | 0.001 | 3841.79145 | UP |
| *LOC101113726* | 3.769403 | 0.001 | 3769.403 | UP |

| | | | | |
|---|---|---|---|---|
| *LOC101113189* | 3.744442 | 0.001 | 3744.442 | UP |
| *LOC105616091* | 3.708944 | 0.001 | 3708.944 | UP |
| *LRRN1* | 6.686132 | 0.001 | 6686.132 | UP |
| *TLR5* | 3.656602 | 0.001 | 3656.60193 | UP |
| *UGT1A1* | 111.2189 | 0.016249 | 6844.64243 | UP |
| *POU2F3* | 7.018347 | 0.001 | 7018.3467 | UP |
| *LOC106991207* | 3.549698 | 0.001 | 3549.698 | UP |
| *LOC106991203* | 3.465721 | 0.001 | 3465.721 | UP |
| *LOC105616415* | 3.339548 | 0.001 | 3339.548 | UP |
| *LOC105614635* | 3.298318 | 0.001 | 3298.318 | UP |
| *LOC105610456* | 3.195928 | 0.001 | 3195.928 | UP |
| *CTRC* | 3.138362 | 0.568199 | 5.52335009 | UP |
| *LOC105604369* | 3.077245 | 0.001 | 3077.245 | UP |
| *LOC101107259* | 3.052632 | 0.001 | 3052.632 | UP |
| *CD207* | 7.652756 | 0.001 | 7652.756 | UP |
| *ATP6V1B1* | 423.4991 | 0.001 | 423499.084 | UP |
| *ACSM4* | 2.836116 | 0.001 | 2836.116 | UP |
| *LOC105607154* | 2.798793 | 0.001 | 2798.793 | UP |
| *LOC105604842* | 2.780288 | 0.001 | 2780.288 | UP |
| *LOC106991405* | 2.765373 | 0.001 | 2765.373 | UP |
| *LOC105615893* | 2.734533 | 0.001 | 2734.533 | UP |
| *LOC101101862* | 2.721172 | 0.001 | 2721.17215 | UP |
| *LOC105615734* | 2.678692 | 0.001 | 2678.692 | UP |
| *LOC105608895* | 2.677712 | 0.001 | 2677.712 | UP |
| *PIGY* | 8.548679 | 0.001 | 8548.679 | UP |
| *C4H7orf31* | 2.648178 | 0.001 | 2648.178 | UP |
| *FCN3* | 2.637651 | 0.001 | 2637.651 | UP |
| *CD177* | 207.6204 | 0.001 | 207620.426 | UP |
| *RBP2* | 412.6506 | 0.001 | 412650.635 | UP |
| *LOC105611549* | 2.584217 | 0.001 | 2584.21726 | UP |
| *LOC101105379* | 2.581682 | 0.001 | 2581.682 | UP |
| *SLC9A2* | 87.31771 | 0.334641 | 260.929507 | UP |
| *CYP4F21* | 2.435596 | 0.001 | 2435.596 | UP |
| *LOC101107795* | 2.416286 | 0.001 | 2416.286 | UP |
| *PDZRN4* | 9.578681 | 0.001 | 9578.681 | UP |
| *LOC105604880* | 2.347296 | 0.001 | 2347.296 | UP |
| *LOC105616289* | 2.327093 | 0.001 | 2327.093 | UP |
| *LOC101112481* | 2.252598 | 0.001 | 2252.598 | UP |
| *LOC101103439* | 202.9718 | 0.883172 | 229.821369 | UP |
| *LOC105614078* | 2.086725 | 0.495517 | 4.21120769 | UP |
| *TACR2* | 10.96095 | 0.001 | 10960.951 | UP |
| *TMEM257* | 2.045439 | 0.001 | 2045.439 | UP |

| | | | | |
|---|---|---|---|---|
| *LOC105615805* | 2.038133 | 0.001 | 2038.133 | UP |
| *LOC106991649* | 2.016159 | 0.001 | 2016.159 | UP |
| *LOC105612436* | 1.962022 | 0.001 | 1962.022 | UP |
| *LOC105609952* | 1.956774 | 0.001 | 1956.774 | UP |
| *LOC106990488* | 1.930994 | 0.001 | 1930.994 | UP |
| *LOC101109940* | 1.887533 | 0.001 | 1887.533 | UP |
| *LOC105604781* | 1.854786 | 0.001 | 1854.786 | UP |
| *LOC106990202* | 1.849398 | 0.001 | 1849.398 | UP |
| *MIP* | 1.83593 | 0.001 | 1835.93 | UP |
| *RNF224* | 12.76474 | 0.001 | 12764.742 | UP |
| *LOC101116576* | 1.83387 | 0.001 | 1833.87003 | UP |
| *CES2* | 453.7567 | 1.883363 | 240.929022 | UP |
| *LOC101120476* | 1.775138 | 0.001 | 1775.138 | UP |
| *LOC105615807* | 1.583146 | 0.001 | 1583.146 | UP |
| *LOC101109475* | 1.536038 | 0.001 | 1536.03826 | UP |
| *LOC101102291* | 1.527983 | 0.001 | 1527.983 | UP |
| *LOC106991725* | 1.345085 | 0.001 | 1345.085 | UP |
| *ENPP3* | 142.6992 | 0.064602 | 2208.89737 | UP |
| *SLC9A3* | 474.4648 | 0.018711 | 25357.5304 | UP |
| *PCP4L1* | 14.81495 | 0.001 | 14814.951 | UP |
| *LOC101121777* | 69.38184 | 0.001 | 69381.836 | UP |
| *MGAT3* | 15.14133 | 0.001 | 15141.333 | UP |
| *AKR1D1* | 405.9698 | 0.001 | 405969.849 | UP |
| *SLC5A8* | 23.88153 | 0.001 | 23881.5347 | UP |
| *KYNU* | 16.94246 | 0.175737 | 96.4080359 | UP |
| *SLC14A1* | 297.2272 | 0.001 | 297227.212 | UP |
| *AQP5* | 327.5083 | 0.001 | 327508.34 | UP |
| *APOBEC1* | 87.08126 | 0.520434 | 167.324386 | UP |
| *ST8SIA5* | 24.01741 | 0.001 | 24017.414 | UP |
| *NOS2* | 40.51643 | 0.001 | 40516.4266 | UP |
| *PGLYRP2* | 37.72605 | 2.594064 | 14.5432229 | UP |
| *ADGRF1* | 9.972401 | 0.270723 | 36.8361779 | UP |
| *CYP11A1* | 43.03505 | 0.001 | 43035.049 | UP |
| *ALOX15B* | 91.7823 | 0.001 | 91782.303 | UP |
| *PADI1* | 524.3741 | 0.275159 | 1905.71323 | UP |
| *SOWAHC* | 19.25818 | 0.001 | 19258.181 | UP |
| *LOC101103238* | 72.68739 | 0.001 | 72687.386 | UP |
| *DUOXA1* | 19.5505 | 0.001 | 19550.5002 | UP |
| *GAP43* | 58.03593 | 0.001 | 58035.931 | UP |
| *HAL* | 18.38686 | 0.001 | 18386.86 | UP |
| *MUC20* | 25.97019 | 0.763773 | 34.0025034 | UP |
| *ADH1C* | 251.6182 | 0.05486 | 4586.55195 | UP |

| | | | | |
|---|---|---|---|---|
| *NXPE2* | 69.01985 | 0.001 | 69019.8477 | UP |
| *LYPD6* | 21.29732 | 0.001 | 21297.3222 | UP |
| *HCN2* | 20.32587 | 0.839213 | 24.2201586 | UP |
| *BVES* | 21.82825 | 0.001 | 21828.2534 | UP |
| *LOC101121908* | 21.94349 | 0.001 | 21943.492 | UP |
| *RNF152* | 22.54105 | 0.001 | 22541.052 | UP |
| *TLL2* | 9.479653 | 0.001 | 9479.65332 | UP |
| *LOC101105864* | 305.7473 | 0.001 | 305747.284 | UP |
| *CAV3* | 31.42564 | 0.001 | 31425.636 | UP |
| *LOC101117044* | 122.2179 | 0.001 | 122217.888 | UP |
| *DUOXA2* | 198.8734 | 1.561859 | 127.331211 | UP |
| *SLC4A9* | 35.48919 | 0.227223 | 156.186623 | UP |
| *UPK1B* | 116.9292 | 0.019238 | 6077.93188 | UP |
| *PATL2* | 9.671702 | 0.119145 | 81.1757645 | UP |
| *IL36B* | 20.29016 | 0.001 | 20290.1597 | UP |
| *DHRS9* | 44.31293 | 4.878036 | 9.08417346 | UP |
| *COLQ* | 8.38233 | 0.273785 | 30.6164235 | UP |
| *PRSS48* | 13.57856 | 0.192909 | 70.3884059 | UP |
| *DAPL1* | 221.588 | 0.967138 | 229.117232 | UP |
| *TNNT1* | 39.8638 | 0.585588 | 68.0748203 | UP |
| *WSCD2* | 14.63309 | 0.432037 | 33.8699977 | UP |
| *KCNG3* | 5.450149 | 0.216132 | 25.2167611 | UP |
| *HSD17B13* | 573.1944 | 0.001 | 573194.397 | UP |
| *LOC101103771* | 9437.467 | 0.317666 | 29708.7739 | UP |
| *LOC101113369* | 376.1786 | 0.001 | 376178.558 | UP |
| *HSBP1L1* | 24.85901 | 4.757486 | 5.22523999 | UP |
| *CLDN3* | 11.39365 | 0.001 | 11393.651 | UP |
| *NOXO1* | 50.31762 | 0.001 | 50317.6198 | UP |
| *KRT35* | 104.6726 | 0.001 | 104672.625 | UP |
| *SYT8* | 8.752247 | 0.001 | 8752.247 | UP |
| *AHSG* | 195.6727 | 0.068086 | 2873.90527 | UP |
| *PRSS27* | 508.3433 | 1.209885 | 420.158331 | UP |
| *HSD17B6* | 25.58527 | 1.669805 | 15.3223119 | UP |
| *LMO3* | 15.94971 | 1.173776 | 13.5883819 | UP |
| *SERPINB10* | 186.4606 | 10.54421 | 17.6836962 | UP |
| *TFF1* | 60.50036 | 0.329656 | 183.525733 | UP |
| *CDKL4* | 7.457469 | 0.001 | 7457.46935 | UP |
| *TPRG1* | 36.48298 | 0.001 | 36482.9804 | UP |
| *GCNT3* | 222.8263 | 0.17479 | 1274.8229 | UP |
| *LOC101112671* | 27.41783 | 0.231755 | 118.305234 | UP |
| *HOXC4* | 38.76928 | 0.001 | 38769.276 | UP |
| *LIX1* | 16.50636 | 0.001 | 16506.363 | UP |

| | | | | |
|---|---|---|---|---|
| *MUC4* | 14.16659 | 0.001 | 14166.592 | UP |
| *OTOP3* | 42.96501 | 0.001 | 42965.008 | UP |
| *MUC1* | 44.92198 | 0.660409 | 68.0214564 | UP |
| *FAM69C* | 53.59932 | 0.001 | 53599.319 | UP |
| *PRSS35* | 59.16396 | 0.106162 | 557.298849 | UP |
| *DUSP27* | 30.64855 | 0.001 | 30648.546 | UP |
| *TTLL10* | 10.74229 | 1.729766 | 6.2102585 | UP |
| *SLC22A18* | 49.78351 | 0.001 | 49783.508 | UP |
| *NPR3* | 20.35754 | 1.140952 | 17.8425886 | UP |
| *SLC34A3* | 50.4657 | 0.001 | 50465.702 | UP |
| *CCL20* | 67.35856 | 0.127704 | 527.459004 | UP |
| *SLC6A14* | 145.0799 | 0.165644 | 875.853878 | UP |
| *TMPRSS11A* | 90.16049 | 3.025581 | 29.7993969 | UP |
| *ANKRD24* | 118.3473 | 7.670643 | 15.428607 | UP |
| *HAP1* | 9.502826 | 0.001 | 9502.826 | UP |
| *VSIG10L* | 56.78225 | 0.001 | 56782.246 | UP |
| *TRIM71* | 5.684439 | 0.001 | 5684.439 | UP |
| *LOC101114973* | 40.0967 | 0.066189 | 605.790918 | UP |
| *KCNG2* | 5.692732 | 0.104435 | 54.5098099 | UP |
| *TNS4* | 58.85282 | 0.001 | 58852.818 | UP |
| *BARX2* | 62.95867 | 0.001 | 62958.668 | UP |
| *TACR1* | 3.084111 | 0.001 | 3084.111 | UP |
| *STYK1* | 16.76237 | 0.390133 | 42.9657916 | UP |
| *SDK2* | 5.660719 | 0.359258 | 15.7566957 | UP |
| *ATP12A* | 198.9999 | 0.001 | 198999.908 | UP |
| *WDR66* | 9.769868 | 0.722948 | 13.5139225 | UP |
| *ADAMTSL2* | 12.56154 | 0.001 | 12561.535 | UP |
| *ARG1* | 87.80626 | 19.56838 | 4.48715111 | UP |
| *TGM3* | 1612.35 | 0.536504 | 3005.28976 | UP |
| *SCGB1C1* | 78.67699 | 0.001 | 78676.9901 | UP |
| *ACER1* | 51.2645 | 0.62209 | 82.4068915 | UP |
| *TCN1* | 147.865 | 0.216195 | 683.942513 | UP |
| *PSD* | 82.57651 | 0.001 | 82576.508 | UP |
| *PAX9* | 85.43823 | 0.001 | 85438.232 | UP |
| *HAND2* | 89.80869 | 0.001 | 89808.693 | UP |
| *ISLR2* | 13.86147 | 1.435449 | 9.65654161 | UP |
| *PRSS22* | 139.8473 | 17.00621 | 8.22330967 | UP |
| *RYR3* | 6.313796 | 0.001 | 6313.796 | UP |
| *FOSL1* | 14.40876 | 0.232422 | 61.9939509 | UP |
| *DUOX2* | 134.689 | 4.101098 | 32.8421862 | UP |
| *IL26* | 2.429616 | 0.001 | 2429.616 | UP |
| *PABPC1L* | 6.479751 | 0.077867 | 83.2156287 | UP |

| | | | | |
|---|---|---|---|---|
| *TGM5* | 31.36367 | 0.001 | 31363.67 | UP |
| *C12H1orf167* | 5.137315 | 0.001 | 5137.315 | UP |
| *NPTXR* | 13.98297 | 0.709887 | 19.6974518 | UP |
| *GLYCTK* | 46.03687 | 0.750751 | 61.3210873 | UP |
| *COL17A1* | 699.5565 | 159.6337 | 4.38226053 | UP |
| *ISG20* | 14.94028 | 0.001 | 14940.276 | UP |
| *TRPC4* | 5.992506 | 0.050705 | 118.183246 | UP |
| *HMGCS2* | 1739.553 | 0.0165 | 105427.468 | UP |
| *NOD2* | 16.35126 | 0.001 | 16351.261 | UP |
| *XKR6* | 3.034569 | 0.001 | 3034.569 | UP |
| *CEMIP* | 59.92732 | 0.788749 | 75.9776748 | UP |
| *WFIKKN2* | 27.73611 | 0.089326 | 310.504276 | UP |
| *SULT2B1* | 198.0508 | 0.001 | 198050.781 | UP |
| *HDC* | 7.606122 | 0.044145 | 172.298607 | UP |
| *LOC101116795* | 72.79969 | 0.001 | 72799.69 | UP |
| *DAB1* | 9.020484 | 0.194841 | 46.2967249 | UP |
| *IL17F* | 13.47894 | 0.147497 | 91.3845231 | UP |
| *TRIM29* | 223.1154 | 0.001 | 223115.372 | UP |
| *NPTX1* | 32.25376 | 0.089037 | 362.251165 | UP |
| *C24H16orf89* | 21.51878 | 0.001 | 21518.782 | UP |
| *OTUB2* | 51.87073 | 0.816859 | 63.5002308 | UP |
| *MACC1* | 27.3185 | 0.001 | 27318.4982 | UP |
| *SMPX* | 272.752 | 0.001 | 272751.953 | UP |
| *RAB19* | 19.37496 | 0.368569 | 52.5680571 | UP |
| *FGF5* | 8.202163 | 0.001 | 8202.1629 | UP |
| *COL24A1* | 5.504262 | 0.069361 | 79.3567331 | UP |
| *CLCA2* | 529.2587 | 66.9393 | 7.90654698 | UP |
| *TMEM95* | 5.083035 | 0.001 | 5083.03494 | UP |
| *KRT17* | 10910.68 | 107.2951 | 101.688545 | UP |
| *C3H9orf172* | 3.506388 | 0.016255 | 215.71135 | UP |
| *PRR19* | 28.87924 | 0.001 | 28879.236 | UP |
| *KLHL30* | 7.828433 | 0.147495 | 53.0759212 | UP |
| *GSDMA* | 32.48624 | 0.001 | 32486.237 | UP |
| *SCN9A* | 5.086692 | 0.836521 | 6.08077296 | UP |
| *HSD17B2* | 77.46463 | 0.118961 | 651.176688 | UP |
| *CLDN23* | 18.11182 | 0.017643 | 1026.57281 | UP |
| *LOC101106609* | 6.427092 | 0.065025 | 98.840323 | UP |
| *IL22* | 2.152969 | 0.001 | 2152.9694 | UP |
| *ZNF662* | 3.911326 | 0.859624 | 4.55004281 | UP |
| *TMEM171* | 42.94245 | 0.497756 | 86.2720571 | UP |
| *TGM1* | 546.7969 | 0.001 | 546796.936 | UP |
| *TMPRSS4* | 166.5452 | 24.55932 | 6.78134279 | UP |

| | | | | |
|---|---|---|---|---|
| *ENDOU* | 602.0472 | 2.963783 | 203.134699 | UP |
| *ATG9B* | 102.6122 | 1.880452 | 54.5678289 | UP |
| *LOC101102014* | 46.60249 | 9.233101 | 5.04732765 | UP |
| *LOC101116267* | 118.3065 | 0.008339 | 14187.1368 | UP |
| *CARTPT* | 19.20504 | 0.001 | 19205.036 | UP |
| *LOC101106871* | 283.2935 | 0.423282 | 669.278299 | UP |
| *PLA2G4E* | 60.2299 | 2.068551 | 29.1169538 | UP |
| *DCST2* | 6.514465 | 0.37594 | 17.32847 | UP |
| *LOC101123254* | 135.5779 | 6.840536 | 19.81977 | UP |
| *KRT78* | 3323.992 | 0.001 | 3323991.94 | UP |
| *KRTDAP* | 1611.144 | 0.001 | 1611144.05 | UP |
| *SLC28A3* | 24.74135 | 0.001 | 24741.3461 | UP |
| *BBOX1* | 17.74198 | 0.001 | 17741.9807 | UP |
| *PDE6A* | 7.740515 | 0.006178 | 1252.91592 | UP |
| *ATP7B* | 7.092519 | 0.173467 | 40.8868488 | UP |
| *SLC26A9* | 21.82183 | 1.761326 | 12.3894333 | UP |
| *SMYD1* | 226.0335 | 0.001 | 226033.477 | UP |
| *MAP3K7CL* | 5.938395 | 0.416558 | 14.2558671 | UP |
| *S100A9* | 4921.764 | 0.001 | 4921764.16 | UP |
| *MUC15* | 60.8342 | 2.601443 | 23.3847899 | UP |
| *FAM83C* | 37.64319 | 5.430504 | 6.93180449 | UP |
| *RAX2* | 7.029022 | 0.001 | 7029.022 | UP |
| *S100A8* | 8764.376 | 0.001 | 8764375.98 | UP |
| *TRPV3* | 24.71477 | 0.003832 | 6449.57389 | UP |
| *MFAP3L* | 10.28041 | 0.058251 | 176.485614 | UP |
| *C14H19orf33* | 374.06604 | 5567.646484 | 0.067185667 | DOWN |
| *CTNNBIP1* | 6.857236984 | 77.613846 | 0.088350692 | DOWN |
| *LOC101110243* | 4.53059 | 47.161739 | 0.096064948 | DOWN |
| *CSTA* | 456.510956 | 4743.056152 | 0.096248271 | DOWN |
| *LOC101112249* | 985.175964 | 8561.130859 | 0.115075447 | DOWN |
| *CAPNS2* | 7.457073 | 62.673996 | 0.11898193 | DOWN |
| *LOC105615767* | 21.144239 | 170.918518 | 0.123709468 | DOWN |
| *LOC101122488* | 6.633393 | 52.140503 | 0.1272215 | DOWN |
| *LOC105610290* | 2.869406 | 18.616365 | 0.154133527 | DOWN |
| *FXYD3* | 249.3502923 | 1513.405273 | 0.164761083 | DOWN |
| *CEBPA* | 8.4515 | 49.922371 | 0.169292841 | DOWN |
| *CDCA4* | 3.161831 | 16.768902 | 0.188553252 | DOWN |
| *S1PR5* | 6.356143 | 31.14323 | 0.204093891 | DOWN |
| *CNFN* | 251.5022392 | 1204.307129 | 0.208835631 | DOWN |
| *RHOV* | 8.617823 | 38.184357 | 0.225689881 | DOWN |
| *TTC22* | 12.333975 | 43.38567102 | 0.284286833 | DOWN |
| *RBBP8NL* | 8.291495 | 28.102636 | 0.295043319 | DOWN |

| | | | | |
|---|---|---|---|---|
| *CLEC14A* | 15.896101 | 52.331429 | 0.303758206 | DOWN |
| *KRT5* | 3509.210693 | 11292.66992 | 0.310751197 | DOWN |
| *KLK10* | 159.4083101 | 476.427094 | 0.334591194 | DOWN |
| *UFSP1* | 5.381833 | 14.690583 | 0.366345774 | DOWN |
| *PAQR7* | 1.911558 | 5.168675 | 0.369835209 | DOWN |
| *WNT10A* | 8.639623 | 20.42939 | 0.422901663 | DOWN |
| *SLC35A4* | 4.440803 | 10.19944208 | 0.435396658 | DOWN |
| *KLHDC9* | 2.579979 | 5.827551355 | 0.442720938 | DOWN |
| *HOXC5* | 9.316009 | 19.644098 | 0.474239591 | DOWN |
| *GJB2* | 138.919601 | 287.189911 | 0.483720339 | DOWN |
| *LIPM* | 9.214626264 | 18.852257 | 0.488781066 | DOWN |
| *LOC101102814* | 1.603957 | 3.22216467 | 0.49778865 | DOWN |

**Supplementary Table 9 |** The gene expression and fold change of 427 rumen up-regulated genes compared to both the FC stomach of camels and cetaceans.

| Official name | Rumen of Sheep | The FC stomach of Camel | The FC stomach of Cetaceans | Fold change of Sheep/Camel | Fold change of Sheep/Cetaceans |
|---|---|---|---|---|---|
| *KLK9* | 37.22844 | 3.14934 | 17.72658 | 11.8210289 | 2.10014809 |
| *LOC101103862* | 707.795 | 8.19744 | 313.5804 | 86.3434173 | 2.25714072 |
| *VSTM2L* | 21.24554 | 2.08555 | 9.238019 | 10.1870188 | 2.2997936 |
| *SRPK3* | 10.64524 | 1.45319 | 4.449088 | 7.32542845 | 2.39267899 |
| *FAM83B* | 18.08906 | 3.69837 | 7.553164 | 4.89108986 | 2.39489835 |
| *ZACN* | 7.692018 | 0.001 | 3.122662 | 7692.018 | 2.46328869 |
| *LOC101121414* | 6.582892 | 2.4795 | 2.640858 | 2.65492717 | 2.49270953 |
| *NPAS1* | 6.030153 | 2.5147 | 2.385069 | 2.39796119 | 2.52829289 |
| *CD70* | 3.024992 | 0.156227 | 1.186656 | 19.362799 | 2.54917348 |
| *CD164L2* | 14.75401 | 0.001 | 5.716718 | 14754.0083 | 2.58085291 |
| *AHNAK2* | 9.03539 | 2.90061 | 3.399268 | 3.1149965 | 2.65803991 |
| *LOC105608435* | 10.02089 | 4.00822 | 3.499694 | 2.50008358 | 2.86336034 |
| *AHRR* | 8.042508 | 2.53616 | 2.530831 | 3.17113589 | 3.17781314 |
| *VTCN1* | 7.740344 | 0.22725 | 2.411588 | 34.0609198 | 3.20964611 |
| *COL7A1* | 146.0332 | 12.6261 | 45.1793 | 11.5659763 | 3.23230283 |
| *B4GALNT3* | 60.21916 | 4.92909 | 17.62143 | 12.2170952 | 3.41738261 |
| *L3MBTL1* | 2.806673 | 1.03394 | 0.77612 | 2.71454146 | 3.61628743 |
| *DSC3* | 93.86417 | 37.789 | 23.66929 | 2.48390203 | 3.96565151 |
| *LOC101112304* | 15279.52 | 0.001 | 0.001 | 15279522.5 | 15279522.5 |
| *LOC101102714* | 10660.87 | 0.001 | 0.001 | 10660870.1 | 10660870.1 |
| *LOC105613035* | 8160.179 | 0.001 | 0.001 | 8160179.2 | 8160179.2 |
| *LOC101113601* | 7766.591 | 0.001 | 0.151575 | 7766590.82 | 51239.3959 |
| *HAS3* | 11.87929 | 4.30417 | 2.57071 | 2.75994914 | 4.62101532 |
| *A2ML1* | 6078.86 | 0.001 | 0.001 | 6078859.89 | 6078859.89 |
| *CCR10* | 2.714588 | 0.997137 | 0.557358 | 2.72238218 | 4.87045669 |
| *LOC101110777* | 5062.031 | 0.001 | 0.001 | 5062030.76 | 5062030.76 |
| *LOC101102105* | 4067.69 | 0.001 | 0.001 | 4067690.19 | 4067690.19 |
| *LOC101111178* | 3783.106 | 0.001 | 0.001 | 3783105.71 | 3783105.71 |
| *SNX31* | 15.88443 | 5.29311 | 2.652551 | 3.00096314 | 5.98835895 |
| *PRD-SPRRII* | 3708.906 | 0.001 | 0.001 | 3708906.49 | 3708906.49 |
| *LOC101104114* | 3006.677 | 0.001 | 0.001 | 3006677.25 | 3006677.25 |
| *LOC105615728* | 2962.506 | 0.001 | 0.001 | 2962506.1 | 2962506.1 |
| *CRISP3* | 1638.922 | 0.001 | 0.001 | 1638922.36 | 1638922.36 |
| *IVL* | 1519.637 | 0.001 | 0.001 | 1519636.65 | 1519636.65 |
| *LOC101107119* | 1239.146 | 0.001 | 0.001 | 1239145.67 | 1239145.67 |
| *VSIG2* | 15.63402 | 3.98627 | 1.870931 | 3.92196665 | 8.35627717 |

| | | | | | |
|---|---|---|---|---|---|
| *ACACB* | 12.15978 | 3.99821 | 1.378151 | 3.04130473 | 8.82325304 |
| *CCDC183* | 17.82192 | 6.03152 | 2.009396 | 2.95479813 | 8.86929406 |
| *LOC101121219* | 1217.002 | 0.001 | 0.001 | 1217002.46 | 1217002.46 |
| *LYPD2* | 1000.9 | 0.001 | 0.001 | 1000899.96 | 1000899.96 |
| *TRIM62* | 12.53244 | 3.29566 | 1.280977 | 3.80270932 | 9.78349703 |
| *C7H14orf169* | 17.8153 | 8.19867 | 1.763522 | 2.17295025 | 10.1021149 |
| *HTR2A* | 3.628845 | 1.09939 | 0.34297 | 3.30078043 | 10.580636 |
| *CPA6* | 4.304087 | 1.26286 | 0.392305 | 3.40820598 | 10.9712774 |
| *EREG* | 12.7423 | 5.10363 | 1.138663 | 2.49671293 | 11.1905797 |
| *KRT79* | 955.0491 | 0.001 | 0.001 | 955049.133 | 955049.133 |
| *TMPRSS13* | 67.04128 | 24.2459 | 5.409249 | 2.76505648 | 12.3938245 |
| *LOC101116141* | 880.8626 | 0.001 | 0.001 | 880862.63 | 880862.63 |
| *LOC101106610* | 720.6295 | 0.001 | 0.001 | 720629.498 | 720629.498 |
| *LOC101122351* | 715.2217 | 0.001 | 0.001 | 715221.68 | 715221.68 |
| *LOC101104808* | 698.7977 | 0.001 | 0.001 | 698797.729 | 698797.729 |
| *LOC101111992* | 621.7982 | 0.001 | 0.001 | 621798.157 | 621798.157 |
| *LOC105604529* | 523.9991 | 0.001 | 0.001 | 523999.084 | 523999.084 |
| *SLC26A3* | 457.0272 | 0.001 | 0.001 | 457027.234 | 457027.234 |
| *KCNK9* | 4.23823 | 1.39798 | 0.273933 | 3.03168143 | 15.4717759 |
| *SBD2* | 452.695 | 0.001 | 0.001 | 452694.977 | 452694.977 |
| *UGT1A9* | 436.0501 | 0.001 | 0.001 | 436050.11 | 436050.11 |
| *LOC101108147* | 414.4206 | 0.001 | 0.001 | 414420.624 | 414420.624 |
| *LOC101103612* | 403.2927 | 0.001 | 0.001 | 403292.725 | 403292.725 |
| *IL36A* | 323.7675 | 0.001 | 0.001 | 323767.474 | 323767.474 |
| *LOC101119393* | 313.4223 | 0.001 | 0.001 | 313422.254 | 313422.254 |
| *LOC101115964* | 303.13 | 0.001 | 0.001 | 303130.005 | 303130.005 |
| *ACTA1* | 279.391 | 0.001 | 0.001 | 279390.991 | 279390.991 |
| *DENND2C* | 40.09808 | 14.8992 | 1.638447 | 2.69129049 | 24.4732209 |
| *LOC101107809* | 260.7116 | 0.001 | 0.001 | 260711.578 | 260711.578 |
| *PI3* | 258.7939 | 0.001 | 0.400744 | 258793.884 | 645.783553 |
| *LOC101115172* | 242.4059 | 0.001 | 0.001 | 242405.93 | 242405.93 |
| *LOC101112555* | 239.5526 | 0.001 | 0.001 | 239552.612 | 239552.612 |
| *STC2* | 12.99392 | 4.39112 | 0.40116 | 2.95913503 | 32.390859 |
| *SERPINB12* | 205.4519 | 0.001 | 0.001 | 205451.87 | 205451.87 |
| *LOC105614373* | 203.7187 | 0.001 | 0.001 | 203718.704 | 203718.704 |
| *KRT23* | 196.9857 | 0.001 | 0.001 | 196985.669 | 196985.669 |
| *DSG3* | 623.4412 | 233.22 | 18.12918 | 2.67318906 | 34.3888206 |
| *JPH2* | 85.95128 | 33.394 | 2.489614 | 2.57385396 | 34.5239378 |
| *GGT6* | 25.65006 | 7.91594 | 0.729627 | 3.24030484 | 35.1550295 |
| *LOC101121267* | 184.4823 | 0.001 | 0.001 | 184482.315 | 184482.315 |
| *LOC101117764* | 182.2246 | 0.001 | 0.001 | 182224.609 | 182224.609 |
| *LOC101120106* | 164.7707 | 0.001 | 0.001 | 164770.676 | 164770.676 |

| | | | | | |
|---|---|---|---|---|---|
| *APOBEC3Z1* | 146.4274 | 0.001 | 0.001 | 146427.383 | 146427.383 |
| *LOC101117163* | 138.395 | 0.001 | 0.001 | 138394.951 | 138394.951 |
| *SCEL* | 134.9509 | 0.001 | 0.001 | 134950.933 | 134950.933 |
| *C1H2orf54* | 133.0843 | 0.001 | 0.001 | 133084.259 | 133084.259 |
| *VILL* | 39.62866 | 12.8796 | 0.738179 | 3.07685472 | 53.6843476 |
| *LOC443322* | 122.7522 | 0.001 | 0.001 | 122752.208 | 122752.208 |
| *LOC101102344* | 119.8675 | 0.001 | 0.001 | 119867.455 | 119867.455 |
| *LOC101108071* | 110.2744 | 0.001 | 0.001 | 110274.368 | 110274.368 |
| *PLA2G2F* | 107.2031 | 0.001 | 0.001 | 107203.15 | 107203.15 |
| *KLHL41* | 57.73458 | 16.7488 | 0.930301 | 3.44708737 | 62.0601042 |
| *IL19* | 99.40583 | 0.001 | 7.251646 | 99405.83 | 13.7080368 |
| *RBPMS2* | 95.17597 | 0.001 | 0.001 | 95175.972 | 95175.972 |
| *KRT80* | 91.30562 | 0.001 | 0.001 | 91305.618 | 91305.618 |
| *LOC101106641* | 90.53042 | 0.001 | 0.001 | 90530.4173 | 90530.4173 |
| *RPTN* | 80.38301 | 0.001 | 0.001 | 80383.011 | 80383.011 |
| *ZNF185* | 75.69835 | 0.001 | 0.001 | 75698.3523 | 75698.3523 |
| *LOC101118712* | 75.6394 | 0.001 | 0.001 | 75639.404 | 75639.404 |
| *LOC106991113* | 73.21864 | 0.001 | 0.001 | 73218.643 | 73218.643 |
| *ATP6V1C2* | 71.62398 | 0.001 | 0.001 | 71623.978 | 71623.978 |
| *KRT36* | 9886.455 | 0.14044 | 1.395109 | 70396.2908 | 7086.51086 |
| *KRT34* | 64.54622 | 0.001 | 0.001 | 64546.219 | 64546.219 |
| *RBM24* | 21.87512 | 8.39911 | 0.245138 | 2.60445676 | 89.2359357 |
| *FUT5* | 57.32155 | 0.001 | 0.001 | 57321.5545 | 57321.5545 |
| *SERPINB13* | 57.01982 | 0.001 | 0.001 | 57019.82 | 57019.82 |
| *LOC101102969* | 56.23896 | 0.001 | 0.001 | 56238.964 | 56238.964 |
| *SLURP1* | 53.91094 | 0.001 | 0.001 | 53910.942 | 53910.942 |
| *LOC101103602* | 51.17694 | 0.001 | 0.001 | 51176.943 | 51176.943 |
| *LOC101107619* | 48.29104 | 0.001 | 0.001 | 48291.035 | 48291.035 |
| *ANKRD9* | 46.08163 | 0.001 | 0.001 | 46081.627 | 46081.627 |
| *FGB* | 16.15834 | 7.25417 | 0.125076 | 2.22745566 | 129.18819 |
| *LOC101115983* | 45.11112 | 0.001 | 0.001 | 45111.118 | 45111.118 |
| *SOX21* | 44.41419 | 0.001 | 0.001 | 44414.188 | 44414.188 |
| *TNFRSF6B* | 41.54738 | 0.001 | 8.651921 | 41547.382 | 4.80209909 |
| *LOC101105114* | 40.2042 | 0.001 | 0.001 | 40204.201 | 40204.201 |
| *MUC21* | 1014.247 | 268.466 | 5.82616 | 3.77793399 | 174.084959 |
| *FAM127A* | 38.44585 | 0.001 | 0.001 | 38445.85 | 38445.85 |
| *LOC101116036* | 37.639 | 0.001 | 0.001 | 37639.004 | 37639.004 |
| *GPT* | 34.75334 | 0.001 | 0.001 | 34753.342 | 34753.342 |
| *LOC105602022* | 34.53469 | 0.001 | 0.001 | 34534.691 | 34534.691 |
| *CARD14* | 34.15244 | 0.001 | 0.001 | 34152.4424 | 34152.4424 |
| *B4GALNT2* | 33.78363 | 0.001 | 0.001 | 33783.635 | 33783.635 |
| *LOC101114663* | 33.39572 | 0.001 | 0.001 | 33395.721 | 33395.721 |

| | | | | | |
|---|---|---|---|---|---|
| *FAM150A* | 32.41682 | 0.001 | 0.001 | 32416.817 | 32416.817 |
| *LOC101114920* | 31.71809 | 0.001 | 0.001 | 31718.086 | 31718.086 |
| *ANKRD23* | 31.14351 | 0.001 | 0.001 | 31143.5123 | 31143.5123 |
| *LOC101105876* | 30.3457 | 0.001 | 0.001 | 30345.701 | 30345.701 |
| *SLC5A1* | 75.42542 | 27.0585 | 0.156051 | 2.78749461 | 483.33784 |
| *CEACAM19* | 29.69046 | 0.001 | 0.001 | 29690.464 | 29690.464 |
| *CHST4* | 28.61247 | 0.001 | 0.001 | 28612.474 | 28612.474 |
| *KLC3* | 28.42285 | 0.001 | 0.001 | 28422.85 | 28422.85 |
| *LOC101107541* | 27.69426 | 0.001 | 0.001 | 27694.26 | 27694.26 |
| *LOC105616317* | 27.0355 | 0.001 | 0.001 | 27035.503 | 27035.503 |
| *LOC101103644* | 26.32515 | 0.001 | 0.001 | 26325.151 | 26325.151 |
| *ADGRF4* | 24.72151 | 0.001 | 0.001 | 24721.5053 | 24721.5053 |
| *CDR1* | 23.52682 | 0.001 | 0.001 | 23526.815 | 23526.815 |
| *LYPD5* | 91.73491 | 27.9778 | 0.09001 | 3.27884641 | 1019.16353 |
| *LOC101106229* | 23.18036 | 0.001 | 0.001 | 23180.359 | 23180.359 |
| *S100A5* | 22.64112 | 0.001 | 0.001 | 22641.1162 | 22641.1162 |
| *LOC101113860* | 33.35526 | 13.2671 | 0.026537 | 2.51413319 | 1256.93396 |
| *P2RY2* | 22.63269 | 0.001 | 0.001 | 22632.69 | 22632.69 |
| *UGT1A4* | 21.08562 | 0.001 | 0.001 | 21085.617 | 21085.617 |
| *GAL3ST2* | 20.68874 | 0.001 | 0.001 | 20688.736 | 20688.736 |
| *ACPP* | 20.05565 | 0.001 | 0.001 | 20055.6542 | 20055.6542 |
| *LOC106990121* | 19.69331 | 0.001 | 0.001 | 19693.3124 | 19693.3124 |
| *LOC105616285* | 19.49552 | 0.001 | 0.001 | 19495.518 | 19495.518 |
| *SFTPC* | 19.14474 | 0.001 | 0.114403 | 19144.745 | 167.344316 |
| *SPINK9* | 18.49817 | 0.001 | 0.001 | 18498.169 | 18498.169 |
| *PPP2R2C* | 17.98931 | 0.001 | 0.001 | 17989.3131 | 17989.3131 |
| *LOC101107098* | 17.47176 | 0.001 | 0.001 | 17471.764 | 17471.764 |
| *GSDMC* | 17.38128 | 0.001 | 0.001 | 17381.283 | 17381.283 |
| *CAND2* | 16.8654 | 0.001 | 0.001 | 16865.402 | 16865.402 |
| *LOC101110524* | 16.52492 | 0.001 | 0.001 | 16524.918 | 16524.918 |
| *PGLYRP4* | 16.22951 | 0.001 | 0.001 | 16229.507 | 16229.507 |
| *LOC101113335* | 15.40575 | 0.001 | 0.001 | 15405.7451 | 15405.7451 |
| *LOC105608915* | 14.53024 | 0.001 | 0.001 | 14530.236 | 14530.236 |
| *LOC105614885* | 13.98985 | 0.001 | 0.001 | 13989.855 | 13989.855 |
| *LOC105612196* | 13.94753 | 0.001 | 0.001 | 13947.5343 | 13947.5343 |
| *FCER2* | 13.90866 | 0.001 | 0.001 | 13908.6621 | 13908.6621 |
| *LOC105614940* | 13.65509 | 0.001 | 0.001 | 13655.092 | 13655.092 |
| *STK32C* | 13.11965 | 0.001 | 0.001 | 13119.646 | 13119.646 |
| *LOC101104661* | 12.93321 | 0.001 | 0.001 | 12933.2127 | 12933.2127 |
| *LOC105616478* | 12.21197 | 0.001 | 0.001 | 12211.966 | 12211.966 |
| *HOXB7* | 12.07503 | 0.001 | 0.001 | 12075.0348 | 12075.0348 |
| *TRPM6* | 12.06659 | 0.001 | 0.001 | 12066.5851 | 12066.5851 |

| | | | | | |
|---|---|---|---|---|---|
| *LOC101120775* | 12.00505 | 0.001 | 0.001 | 12005.045 | 12005.045 |
| *LTB4R2* | 11.96399 | 0.001 | 0.001 | 11963.988 | 11963.988 |
| *WNT7B* | 11.95372 | 0.001 | 0.001 | 11953.721 | 11953.721 |
| *HSPA1A* | 11.91347 | 0.001 | 0.001 | 11913.466 | 11913.466 |
| *SERPINB11* | 11.74515 | 0.001 | 0.001 | 11745.147 | 11745.147 |
| *LOC105604082* | 10.93128 | 0.001 | 0.001 | 10931.283 | 10931.283 |
| *KIF26A* | 10.66053 | 0.001 | 0.001 | 10660.532 | 10660.532 |
| *LOC105611786* | 10.54538 | 0.001 | 0.001 | 10545.382 | 10545.382 |
| *ADTRP* | 10.45416 | 0.001 | 0.001 | 10454.1605 | 10454.1605 |
| *LOC101121036* | 10.1539 | 0.001 | 0.001 | 10153.899 | 10153.899 |
| *LOC105605155* | 9.557295 | 0.001 | 0.001 | 9557.295 | 9557.295 |
| *C13H10orf113* | 9.206914 | 0.001 | 0.001 | 9206.914 | 9206.914 |
| *LOC105610609* | 8.921119 | 0.001 | 0.001 | 8921.119 | 8921.119 |
| *IL17REL* | 8.722609 | 0.001 | 0.001 | 8722.60876 | 8722.60876 |
| *ATP13A5* | 8.697449 | 0.001 | 0.001 | 8697.44925 | 8697.44925 |
| *LOC105612090* | 8.549753 | 0.001 | 0.001 | 8549.753 | 8549.753 |
| *LOC101117587* | 8.327113 | 0.001 | 0.001 | 8327.113 | 8327.113 |
| *LRFN4* | 8.313799 | 0.001 | 0.001 | 8313.799 | 8313.799 |
| *A1BG* | 8.307885 | 0.001 | 0.001 | 8307.885 | 8307.885 |
| *LOC101117120* | 8.285954 | 0.001 | 0.001 | 8285.954 | 8285.954 |
| *LOC101105566* | 8.140181 | 0.001 | 0.001 | 8140.181 | 8140.181 |
| *ADGRF2* | 7.952637 | 0.001 | 0.001 | 7952.63725 | 7952.63725 |
| *WDR38* | 3.383704 | 1.54204 | 0.001 | 2.19430348 | 3383.70374 |
| *LOC101110696* | 7.895658 | 0.001 | 0.001 | 7895.658 | 7895.658 |
| *BHLHE23* | 7.758808 | 0.001 | 0.001 | 7758.808 | 7758.808 |
| *LOC105608606* | 7.647254 | 0.001 | 0.001 | 7647.254 | 7647.254 |
| *MCCD1* | 7.555327 | 0.001 | 0.001 | 7555.327 | 7555.327 |
| *LOC105606870* | 7.53356 | 0.001 | 0.001 | 7533.56009 | 7533.56009 |
| *LOC101120435* | 7.258325 | 0.001 | 0.001 | 7258.325 | 7258.325 |
| *LOC105604589* | 7.234852 | 0.001 | 0.001 | 7234.852 | 7234.852 |
| *LOC106991452* | 6.686908 | 0.001 | 0.001 | 6686.908 | 6686.908 |
| *LOC101105174* | 6.561944 | 0.001 | 0.001 | 6561.944 | 6561.944 |
| *LOC106991450* | 6.491086 | 0.001 | 0.001 | 6491.086 | 6491.086 |
| *LOC101115778* | 6.490631 | 0.001 | 0.001 | 6490.631 | 6490.631 |
| *VAX2* | 6.132115 | 0.001 | 0.001 | 6132.115 | 6132.115 |
| *LOC105615508* | 5.752361 | 0.001 | 0.001 | 5752.361 | 5752.361 |
| *C1QL1* | 5.433838 | 0.001 | 0.001 | 5433.838 | 5433.838 |
| *LOC101104501* | 5.417474 | 0.001 | 0.001 | 5417.474 | 5417.474 |
| *KLF13* | 5.207161 | 0.001 | 0.001 | 5207.161 | 5207.161 |
| *POC1A* | 5.195816 | 0.001 | 0.001 | 5195.8163 | 5195.8163 |
| *LOC101105258* | 5.052308 | 0.001 | 0.001 | 5052.308 | 5052.308 |
| *SH2D7* | 4.951905 | 0.001 | 0.09436 | 4951.905 | 52.4788576 |

| | | | | | |
|---|---|---|---|---|---|
| *LOC101122314* | 4.784063 | 0.001 | 0.001 | 4784.063 | 4784.063 |
| *LOC105602127* | 4.748694 | 0.001 | 0.001 | 4748.694 | 4748.694 |
| *LOC101120688* | 4.672022 | 0.001 | 0.001 | 4672.022 | 4672.022 |
| *LOC106990150* | 5.030138 | 1.56477 | 0.001 | 3.21461812 | 5030.138 |
| *LOC101111245* | 4.639668 | 0.001 | 0.001 | 4639.668 | 4639.668 |
| *LOC101104703* | 4.55153 | 0.001 | 0.001 | 4551.53 | 4551.53 |
| *LOC101116196* | 4.509582 | 0.001 | 0.001 | 4509.58228 | 4509.58228 |
| *NKX2-2* | 4.422982 | 0.001 | 0.001 | 4422.982 | 4422.982 |
| *APELA* | 4.212973 | 0.001 | 0.049898 | 4212.973 | 84.4319504 |
| *SPATA12* | 4.126355 | 0.001 | 0.001 | 4126.355 | 4126.355 |
| *LOC105615865* | 4.101434 | 0.001 | 0.001 | 4101.434 | 4101.434 |
| *LOC105605056* | 4.023712 | 0.001 | 0.001 | 4023.712 | 4023.712 |
| *LOC101106977* | 3.967016 | 0.001 | 0.001 | 3967.016 | 3967.016 |
| *LOC101114497* | 3.944958 | 0.001 | 0.001 | 3944.958 | 3944.958 |
| *LOC105606498* | 3.920981 | 0.001 | 0.001 | 3920.981 | 3920.981 |
| *ARSE* | 3.907126 | 0.001 | 0.001 | 3907.12556 | 3907.12556 |
| *LOC106990135* | 3.841791 | 0.001 | 0.001 | 3841.79145 | 3841.79145 |
| *LOC101113726* | 3.769403 | 0.001 | 0.001 | 3769.403 | 3769.403 |
| *LOC101113189* | 3.744442 | 0.001 | 0.001 | 3744.442 | 3744.442 |
| *LOC105616091* | 3.708944 | 0.001 | 0.001 | 3708.944 | 3708.944 |
| *LRRN1* | 6.686132 | 1.94715 | 0.001 | 3.43380428 | 6686.132 |
| *TLR5* | 3.656602 | 0.001 | 0.001 | 3656.60193 | 3656.60193 |
| *POU2F3* | 7.018347 | 1.87832 | 0.001 | 3.73650214 | 7018.3467 |
| *LOC106991207* | 3.549698 | 0.001 | 0.001 | 3549.698 | 3549.698 |
| *LOC106991203* | 3.465721 | 0.001 | 0.001 | 3465.721 | 3465.721 |
| *LOC105616415* | 3.339548 | 0.001 | 0.001 | 3339.548 | 3339.548 |
| *LOC105614635* | 3.298318 | 0.001 | 0.001 | 3298.318 | 3298.318 |
| *LOC105610456* | 3.195928 | 0.001 | 0.001 | 3195.928 | 3195.928 |
| *CTRC* | 3.138362 | 0.001 | 0.568199 | 3138.362 | 5.52335009 |
| *LOC105604369* | 3.077245 | 0.001 | 0.001 | 3077.245 | 3077.245 |
| *LOC101107259* | 3.052632 | 0.001 | 0.001 | 3052.632 | 3052.632 |
| *CD207* | 7.652756 | 1.92049 | 0.001 | 3.98479346 | 7652.756 |
| *ATP6V1B1* | 423.4991 | 0.147714 | 0.001 | 2867.02062 | 423499.084 |
| *ACSM4* | 2.836116 | 0.001 | 0.001 | 2836.116 | 2836.116 |
| *LOC105607154* | 2.798793 | 0.001 | 0.001 | 2798.793 | 2798.793 |
| *LOC105604842* | 2.780288 | 0.001 | 0.001 | 2780.288 | 2780.288 |
| *LOC106991405* | 2.765373 | 0.001 | 0.001 | 2765.373 | 2765.373 |
| *LOC105615893* | 2.734533 | 0.001 | 0.001 | 2734.533 | 2734.533 |
| *LOC101101862* | 2.721172 | 0.001 | 0.001 | 2721.17215 | 2721.17215 |
| *LOC105615734* | 2.678692 | 0.001 | 0.001 | 2678.692 | 2678.692 |
| *LOC105608895* | 2.677712 | 0.001 | 0.001 | 2677.712 | 2677.712 |
| *C4H7orf31* | 2.648178 | 0.001 | 0.001 | 2648.178 | 2648.178 |

| | | | | | |
|---|---|---|---|---|---|
| *FCN3* | 2.637651 | 0.001 | 0.001 | 2637.651 | 2637.651 |
| *CD177* | 207.6204 | 0.078841 | 0.001 | 2633.40012 | 207620.426 |
| *RBP2* | 412.6506 | 0.158464 | 0.001 | 2604.0655 | 412650.635 |
| *LOC105611549* | 2.584217 | 0.001 | 0.001 | 2584.21726 | 2584.21726 |
| *LOC101105379* | 2.581682 | 0.001 | 0.001 | 2581.682 | 2581.682 |
| *SLC9A2* | 87.31771 | 0.034954 | 0.334641 | 2498.09637 | 260.929507 |
| *CYP4F21* | 2.435596 | 0.001 | 0.001 | 2435.596 | 2435.596 |
| *LOC101107795* | 2.416286 | 0.001 | 0.001 | 2416.286 | 2416.286 |
| *LOC105604880* | 2.347296 | 0.001 | 0.001 | 2347.296 | 2347.296 |
| *LOC105616289* | 2.327093 | 0.001 | 0.001 | 2327.093 | 2327.093 |
| *LOC101112481* | 2.252598 | 0.001 | 0.001 | 2252.598 | 2252.598 |
| *LOC101103439* | 202.9718 | 0.095706 | 0.883172 | 2120.7756 | 229.821369 |
| *LOC105614078* | 2.086725 | 0.001 | 0.495517 | 2086.725 | 4.21120769 |
| *TMEM257* | 2.045439 | 0.001 | 0.001 | 2045.439 | 2045.439 |
| *LOC105615805* | 2.038133 | 0.001 | 0.001 | 2038.133 | 2038.133 |
| *LOC106991649* | 2.016159 | 0.001 | 0.001 | 2016.159 | 2016.159 |
| *LOC105612436* | 1.962022 | 0.001 | 0.001 | 1962.022 | 1962.022 |
| *LOC105609952* | 1.956774 | 0.001 | 0.001 | 1956.774 | 1956.774 |
| *LOC106990488* | 1.930994 | 0.001 | 0.001 | 1930.994 | 1930.994 |
| *LOC101109940* | 1.887533 | 0.001 | 0.001 | 1887.533 | 1887.533 |
| *LOC105604781* | 1.854786 | 0.001 | 0.001 | 1854.786 | 1854.786 |
| *LOC106990202* | 1.849398 | 0.001 | 0.001 | 1849.398 | 1849.398 |
| *MIP* | 1.83593 | 0.001 | 0.001 | 1835.93 | 1835.93 |
| *RNF224* | 12.76474 | 4.91109 | 0.001 | 2.59916678 | 12764.742 |
| *LOC101116576* | 1.83387 | 0.001 | 0.001 | 1833.87003 | 1833.87003 |
| *CES2* | 453.7567 | 0.255181 | 1.883363 | 1778.17606 | 240.929022 |
| *LOC101120476* | 1.775138 | 0.001 | 0.001 | 1775.138 | 1775.138 |
| *LOC105615807* | 1.583146 | 0.001 | 0.001 | 1583.146 | 1583.146 |
| *LOC101109475* | 1.536038 | 0.001 | 0.001 | 1536.03826 | 1536.03826 |
| *LOC101102291* | 1.527983 | 0.001 | 0.001 | 1527.983 | 1527.983 |
| *LOC106991725* | 1.345085 | 0.001 | 0.001 | 1345.085 | 1345.085 |
| *ENPP3* | 142.6992 | 0.107065 | 0.064602 | 1332.82761 | 2208.89737 |
| *SLC9A3* | 474.4648 | 0.381049 | 0.018711 | 1245.15417 | 25357.5304 |
| *LOC101121777* | 69.38184 | 0.061762 | 0.001 | 1123.36872 | 69381.836 |
| *AKR1D1* | 405.9698 | 0.422032 | 0.001 | 961.940917 | 405969.849 |
| *SLC5A8* | 23.88153 | 0.031202 | 0.001 | 765.374929 | 23881.5347 |
| *KYNU* | 16.94246 | 0.023661 | 0.175737 | 716.049998 | 96.4080359 |
| *SLC14A1* | 297.2272 | 0.503768 | 0.001 | 590.008122 | 297227.212 |
| *AQP5* | 327.5083 | 0.558485 | 0.001 | 586.422805 | 327508.34 |
| *APOBEC1* | 87.08126 | 0.154748 | 0.520434 | 562.729476 | 167.324386 |
| *ST8SIA5* | 24.01741 | 0.04916 | 0.001 | 488.559003 | 24017.414 |
| *NOS2* | 40.51643 | 0.113414 | 0.001 | 357.243609 | 40516.4266 |

| | | | | | |
|---|---|---|---|---|---|
| *PGLYRP2* | 37.72605 | 0.112852 | 2.594064 | 334.296698 | 14.5432229 |
| *ADGRF1* | 9.972401 | 0.031201 | 0.270723 | 319.616968 | 36.8361779 |
| *CYP11A1* | 43.03505 | 0.146662 | 0.001 | 293.430125 | 43035.049 |
| *ALOX15B* | 91.7823 | 0.318433 | 0.001 | 288.231129 | 91782.303 |
| *PADI1* | 524.3741 | 1.83873 | 0.275159 | 285.182787 | 1905.71323 |
| *LOC101103238* | 72.68739 | 0.284193 | 0.001 | 255.767686 | 72687.386 |
| *GAP43* | 58.03593 | 0.259051 | 0.001 | 224.032839 | 58035.931 |
| *HAL* | 18.38686 | 0.083309 | 0.001 | 220.706237 | 18386.86 |
| *MUC20* | 25.97019 | 0.120507 | 0.763773 | 215.507763 | 34.0025034 |
| *ADH1C* | 251.6182 | 1.18117 | 0.05486 | 213.024577 | 4586.55195 |
| *NXPE2* | 69.01985 | 0.345316 | 0.001 | 199.874456 | 69019.8477 |
| *HCN2* | 20.32587 | 0.132458 | 0.839213 | 153.451449 | 24.2201586 |
| *TLL2* | 9.479653 | 0.061894 | 0.001 | 153.16023 | 9479.65332 |
| *LOC101105864* | 305.7473 | 2.03272 | 0.001 | 150.412887 | 305747.284 |
| *CAV3* | 31.42564 | 0.220915 | 0.001 | 142.25216 | 31425.636 |
| *LOC101117044* | 122.2179 | 0.932664 | 0.001 | 131.041713 | 122217.888 |
| *DUOXA2* | 198.8734 | 1.90818 | 1.561859 | 104.221508 | 127.331211 |
| *SLC4A9* | 35.48919 | 0.355289 | 0.227223 | 99.8882403 | 156.186623 |
| *UPK1B* | 116.9292 | 1.24974 | 0.019238 | 93.5628451 | 6077.93188 |
| *PATL2* | 9.671702 | 0.10435 | 0.119145 | 92.6852132 | 81.1757645 |
| *IL36B* | 20.29016 | 0.239877 | 0.001 | 84.5856821 | 20290.1597 |
| *DHRS9* | 44.31293 | 0.552439 | 4.878036 | 80.2132456 | 9.08417346 |
| *COLQ* | 8.38233 | 0.105784 | 0.273785 | 79.2400537 | 30.6164235 |
| *PRSS48* | 13.57856 | 0.175345 | 0.192909 | 77.4390887 | 70.3884059 |
| *DAPL1* | 221.588 | 3.04582 | 0.967138 | 72.7515027 | 229.117232 |
| *TNNT1* | 39.8638 | 0.561781 | 0.585588 | 70.9596762 | 68.0748203 |
| *WSCD2* | 14.63309 | 0.214665 | 0.432037 | 68.1671077 | 33.8699977 |
| *KCNG3* | 5.450149 | 0.080644 | 0.216132 | 67.5825694 | 25.2167611 |
| *HSD17B13* | 573.1944 | 8.64125 | 0.001 | 66.3323474 | 573194.397 |
| *LOC101113369* | 376.1786 | 5.97321 | 0.001 | 62.9776214 | 376178.558 |
| *HSBP1L1* | 24.85901 | 0.45318 | 4.757486 | 54.8545966 | 5.22523999 |
| *CLDN3* | 11.39365 | 0.215345 | 0.001 | 52.9088254 | 11393.651 |
| *NOXO1* | 50.31762 | 0.984273 | 0.001 | 51.1216094 | 50317.6198 |
| *KRT35* | 104.6726 | 2.08632 | 0.001 | 50.170935 | 104672.625 |
| *SYT8* | 8.752247 | 0.186769 | 0.001 | 46.8613474 | 8752.247 |
| *AHSG* | 195.6727 | 4.20157 | 0.068086 | 46.5713326 | 2873.90527 |
| *PRSS27* | 508.3433 | 11.9364 | 1.209885 | 42.5876531 | 420.158331 |
| *HSD17B6* | 25.58527 | 0.608613 | 1.669805 | 42.0386471 | 15.3223119 |
| *LMO3* | 15.94971 | 0.389392 | 1.173776 | 40.960549 | 13.5883819 |
| *SERPINB10* | 186.4606 | 4.59679 | 10.54421 | 40.5632163 | 17.6836962 |
| *TFF1* | 60.50036 | 1.49514 | 0.329656 | 40.4646782 | 183.525733 |
| *CDKL4* | 7.457469 | 0.185013 | 0.001 | 40.3078127 | 7457.46935 |

| | | | | | |
|---|---|---|---|---|---|
| *GCNT3* | 222.8263 | 5.56397 | 0.17479 | 40.0480761 | 1274.8229 |
| *LOC101112671* | 27.41783 | 0.711735 | 0.231755 | 38.5225252 | 118.305234 |
| *LIX1* | 16.50636 | 0.435207 | 0.001 | 37.9276138 | 16506.363 |
| *MUC4* | 14.16659 | 0.378134 | 0.001 | 37.4644756 | 14166.592 |
| *MUC1* | 44.92198 | 1.21103 | 0.660409 | 37.094029 | 68.0214564 |
| *FAM69C* | 53.59932 | 1.52461 | 0.001 | 35.1560852 | 53599.319 |
| *PRSS35* | 59.16396 | 1.73305 | 0.106162 | 34.1386344 | 557.298849 |
| *DUSP27* | 30.64855 | 0.980912 | 0.001 | 31.2449496 | 30648.546 |
| *TTLL10* | 10.74229 | 0.352866 | 1.729766 | 30.442984 | 6.2102585 |
| *NPR3* | 20.35754 | 0.69093 | 1.140952 | 29.4639629 | 17.8425886 |
| *CCL20* | 67.35856 | 2.34706 | 0.127704 | 28.6991224 | 527.459004 |
| *SLC6A14* | 145.0799 | 5.53319 | 0.165644 | 26.2199454 | 875.853878 |
| *TMPRSS11A* | 90.16049 | 3.56615 | 3.025581 | 25.2823042 | 29.7993969 |
| *ANKRD24* | 118.3473 | 4.78033 | 7.670643 | 24.7571477 | 15.428607 |
| *HAP1* | 9.502826 | 0.392372 | 0.001 | 24.2189198 | 9502.826 |
| *TRIM71* | 5.684439 | 0.236394 | 0.001 | 24.0464606 | 5684.439 |
| *LOC101114973* | 40.0967 | 1.70907 | 0.066189 | 23.4611192 | 605.790918 |
| *KCNG2* | 5.692732 | 0.246249 | 0.104435 | 23.1177873 | 54.5098099 |
| *TACR1* | 3.084111 | 0.139691 | 0.001 | 22.0780938 | 3084.111 |
| *STYK1* | 16.76237 | 0.759497 | 0.390133 | 22.0703613 | 42.9657916 |
| *SDK2* | 5.660719 | 0.259939 | 0.359258 | 21.7771054 | 15.7566957 |
| *ATP12A* | 198.9999 | 9.21291 | 0.001 | 21.6001142 | 198999.908 |
| *WDR66* | 9.769868 | 0.462412 | 0.722948 | 21.128059 | 13.5139225 |
| *ADAMTSL2* | 12.56154 | 0.613372 | 0.001 | 20.4794725 | 12561.535 |
| *ARG1* | 87.80626 | 4.52556 | 19.56838 | 19.4022982 | 4.48715111 |
| *TGM3* | 1612.35 | 84.0667 | 0.536504 | 19.1794132 | 3005.28976 |
| *SCGB1C1* | 78.67699 | 4.1597 | 0.001 | 18.914102 | 78676.9901 |
| *ACER1* | 51.2645 | 2.81016 | 0.62209 | 18.2425567 | 82.4068915 |
| *TCN1* | 147.865 | 8.11064 | 0.216195 | 18.2309844 | 683.942513 |
| *PAX9* | 85.43823 | 32.8803 | 0.001 | 2.59846267 | 85438.232 |
| *ISLR2* | 13.86147 | 0.786461 | 1.435449 | 17.6251244 | 9.65654161 |
| *PRSS22* | 139.8473 | 7.96451 | 17.00621 | 17.5588065 | 8.22330967 |
| *RYR3* | 6.313796 | 0.383915 | 0.001 | 16.4458174 | 6313.796 |
| *FOSL1* | 14.40876 | 0.941384 | 0.232422 | 15.3059305 | 61.9939509 |
| *DUOX2* | 134.689 | 8.87378 | 4.101098 | 15.1783146 | 32.8421862 |
| *IL26* | 2.429616 | 0.167568 | 0.001 | 14.4992839 | 2429.616 |
| *PABPC1L* | 6.479751 | 0.487187 | 0.077867 | 13.3003372 | 83.2156287 |
| *TGM5* | 31.36367 | 2.39913 | 0.001 | 13.0729348 | 31363.67 |
| *C12H1orf167* | 5.137315 | 0.396574 | 0.001 | 12.9542406 | 5137.315 |
| *NPTXR* | 13.98297 | 1.08789 | 0.709887 | 12.8532894 | 19.6974518 |
| *GLYCTK* | 46.03687 | 3.59286 | 0.750751 | 12.8134335 | 61.3210873 |
| *COL17A1* | 699.5565 | 56.0719 | 159.6337 | 12.4760623 | 4.38226053 |

| | | | | | |
|---|---|---|---|---|---|
| *ISG20* | 14.94028 | 1.21248 | 0.001 | 12.3220804 | 14940.276 |
| *TRPC4* | 5.992506 | 0.492901 | 0.050705 | 12.157626 | 118.183246 |
| *HMGCS2* | 1739.553 | 145.036 | 0.0165 | 11.993941 | 105427.468 |
| *NOD2* | 16.35126 | 1.37817 | 0.001 | 11.8644732 | 16351.261 |
| *XKR6* | 3.034569 | 0.257843 | 0.001 | 11.7690571 | 3034.569 |
| *CEMIP* | 59.92732 | 5.11767 | 0.788749 | 11.7098826 | 75.9776748 |
| *WFIKKN2* | 27.73611 | 2.489 | 0.089326 | 11.1434733 | 310.504276 |
| *HDC* | 7.606122 | 0.710336 | 0.044145 | 10.7077805 | 172.298607 |
| *LOC101116795* | 72.79969 | 6.96937 | 0.001 | 10.445663 | 72799.69 |
| *DAB1* | 9.020484 | 0.864142 | 0.194841 | 10.4386597 | 46.2967249 |
| *IL17F* | 13.47894 | 1.29425 | 0.147497 | 10.4144817 | 91.3845231 |
| *NPTX1* | 32.25376 | 3.43777 | 0.089037 | 9.3821742 | 362.251165 |
| *C24H16orf89* | 21.51878 | 2.31928 | 0.001 | 9.27821652 | 21518.782 |
| *OTUB2* | 51.87073 | 5.59108 | 0.816859 | 9.27740884 | 63.5002308 |
| *MACC1* | 27.3185 | 3.14652 | 0.001 | 8.68213081 | 27318.4982 |
| *RAB19* | 19.37496 | 2.33332 | 0.368569 | 8.30360259 | 52.5680571 |
| *FGF5* | 8.202163 | 1.00049 | 0.001 | 8.19814581 | 8202.1629 |
| *COL24A1* | 5.504262 | 0.700998 | 0.069361 | 7.85203719 | 79.3567331 |
| *CLCA2* | 529.2587 | 68.0284 | 66.9393 | 7.77996731 | 7.90654698 |
| *TMEM95* | 5.083035 | 0.65705 | 0.001 | 7.73614632 | 5083.03494 |
| *KRT17* | 10910.68 | 1418.74 | 107.2951 | 7.69039974 | 101.688545 |
| *C3H9orf172* | 3.506388 | 0.457804 | 0.016255 | 7.65914671 | 215.71135 |
| *PRR19* | 28.87924 | 3.84085 | 0.001 | 7.51897002 | 28879.236 |
| *KLHL30* | 7.828433 | 1.08373 | 0.147495 | 7.2236009 | 53.0759212 |
| *GSDMA* | 32.48624 | 4.57024 | 0.001 | 7.10821248 | 32486.237 |
| *SCN9A* | 5.086692 | 0.719333 | 0.836521 | 7.07140116 | 6.08077296 |
| *HSD17B2* | 77.46463 | 11.3814 | 0.118961 | 6.80624791 | 651.176688 |
| *CLDN23* | 18.11182 | 2.66955 | 0.017643 | 6.78459815 | 1026.57281 |
| *LOC101106609* | 6.427092 | 0.949223 | 0.065025 | 6.77089788 | 98.840323 |
| *IL22* | 2.152969 | 0.324966 | 0.001 | 6.62521434 | 2152.9694 |
| *ZNF662* | 3.911326 | 0.604856 | 0.859624 | 6.4665408 | 4.55004281 |
| *TMEM171* | 42.94245 | 6.6906 | 0.497756 | 6.41832617 | 86.2720571 |
| *TMPRSS4* | 166.5452 | 26.3746 | 24.55932 | 6.31460424 | 6.78134279 |
| *ENDOU* | 602.0472 | 96.8987 | 2.963783 | 6.21316041 | 203.134699 |
| *ATG9B* | 102.6122 | 16.8227 | 1.880452 | 6.09962628 | 54.5678289 |
| *LOC101102014* | 46.60249 | 7.75254 | 9.233101 | 6.01125386 | 5.04732765 |
| *LOC101116267* | 118.3065 | 20.1225 | 0.008339 | 5.87931589 | 14187.1368 |
| *CARTPT* | 19.20504 | 3.26976 | 0.001 | 5.87353078 | 19205.036 |
| *LOC101106871* | 283.2935 | 48.5574 | 0.423282 | 5.8341974 | 669.278299 |
| *PLA2G4E* | 60.2299 | 10.3818 | 2.068551 | 5.80148953 | 29.1169538 |
| *DCST2* | 6.514465 | 1.13553 | 0.37594 | 5.73693782 | 17.32847 |
| *LOC101123254* | 135.5779 | 23.9799 | 6.840536 | 5.65381215 | 19.81977 |

| | | | | | |
|---|---|---|---|---|---|
| *KRT78* | 3323.992 | 595.64 | 0.001 | 5.58053848 | 3323991.94 |
| *SLC28A3* | 24.74135 | 4.61379 | 0.001 | 5.36247773 | 24741.3461 |
| *BBOX1* | 17.74198 | 3.32118 | 0.001 | 5.3420714 | 17741.9807 |
| *PDE6A* | 7.740515 | 1.52746 | 0.006178 | 5.06757268 | 1252.91592 |
| *ATP7B* | 7.092519 | 1.4466 | 0.173467 | 4.90288884 | 40.8868488 |
| *SLC26A9* | 21.82183 | 4.53372 | 1.761326 | 4.81322865 | 12.3894333 |
| *SMYD1* | 226.0335 | 49.4949 | 0.001 | 4.56680339 | 226033.477 |
| *MAP3K7CL* | 5.938395 | 1.33431 | 0.416558 | 4.4505366 | 14.2558671 |
| *MUC15* | 60.8342 | 13.9031 | 2.601443 | 4.37558516 | 23.3847899 |
| *FAM83C* | 37.64319 | 8.64313 | 5.430504 | 4.35527315 | 6.93180449 |
| *RAX2* | 7.029022 | 1.644 | 0.001 | 4.27556083 | 7029.022 |
| *TRPV3* | 24.71477 | 5.7938 | 0.003832 | 4.26572666 | 6449.57389 |
| *MFAP3L* | 10.28041 | 2.48604 | 0.058251 | 4.13525373 | 176.485614 |

**Supplementary Table 10 |** KEGG pathway enrichment results for 427 rumen up-regulated genes compared to both the FC stomachs of camels and cetaceans.

| **Ko ID** | **Description** | **GeneRatio** | **BgRatio** | ***p* value** | **Adjusted *p* value** | **Gene Entrez ID** | **Count** |
|---|---|---|---|---|---|---|---|
| oas00830 | Retinol metabolism | 13/139 | 74/8981 | 7.76E-11 | 1.54E-08 | 101104808/442993/101119393/105614373/101117764/101117163/101106641/100534646/101104703/101103439/101111965/101120807/101105400 | 13 |
| oas00140 | Steroid hormone biosynthesis | 11/139 | 69/8981 | 6.95E-09 | 6.88E-07 | 442993/101119393/105614373/101117764/101117163/101106641/100534646/101103641/100048994/101105400/100913161 | 11 |
| oas05204 | Chemical carcinogenesis | 10/139 | 93/8981 | 1.54E-06 | 0.0001019 | 442993/101119393/101115172/105614373/101117764/101117163/101106641/101114663/100534646/101111965 | 10 |
| oas00072 | Synthesis and degradation of ketone bodies | 4/139 | 12/8981 | 2.47E-05 | 0.0012087 | 101111590/101104112/101119699/101104601 | 4 |
| oas00980 | Metabolism of xenobiotics by cytochrome P450 | 8/139 | 80/8981 | 3.05E-05 | 0.0012087 | 442993/101119393/101115172/101117764/101117163/101114663/100534646/101111965 | 8 |
| oas00650 | Butanoate metabolism | 5/139 | 29/8981 | 7.28E-05 | 0.0024013 | 101120794/101111590/101104112/101119699/101104601 | 5 |
| oas00053 | Ascorbate and aldarate metabolism | 5/139 | 30/8981 | 8.62E-05 | 0.0024394 | 442993/101119393/101117764/101117163/100534646 | 5 |
| oas00590 | Arachidonic acid metabolism | 7/139 | 75/8981 | 0.000151 | 0.0033215 | 101115172/105614373/101106664/101106641/101114663/101121185/101115878 | 7 |
| oas00982 | Drug metabolism - cytochrome P450 | 7/139 | 75/8981 | 0.000151 | 0.0033215 | 101104808/442993/101119393/101117764/101117163/100534646/101111965 | 7 |

| | | | | | | | |
|---|---|---|---|---|---|---|---|
| oas00040 | Pentose and glucuronate interconversions | 5/139 | 38/8981 | 0.000275 | 0.005008 | 442993/101119393/101117764/101117163/100534646 | 5 |
| oas04915 | Estrogen signaling pathway | 9/139 | 138/8981 | 0.0002782 | 0.005008 | 101108147/101111379/101118712/101117175/100526781/100913152/100141296/101104450/100303604 | 9 |
| oas00860 | Porphyrin and chlorophyll metabolism | 5/139 | 48/8981 | 0.000829 | 0.013461 | 442993/101119393/101117764/101117163/100534646 | 5 |
| oas05150 | Staphylococcus aureus infection | 7/139 | 100/8981 | 0.0008838 | 0.013461 | 101108147/101111379/101118712/101117175/100526781/100141296/100303604 | 7 |
| oas04978 | Mineral absorption | 5/139 | 52/8981 | 0.0011975 | 0.0169363 | 100462650/492300/101114337/494437/101111646 | 5 |
| oas00983 | Drug metabolism - other enzymes | 6/139 | 90/8981 | 0.0026485 | 0.0349599 | 442993/101119393/101117764/101117163/100534646/101109730 | 6 |
| oas04060 | Cytokine-cytokine receptor interaction | 13/139 | 359/8981 | 0.003675 | 0.0454785 | 101105074/678671/101106251/101112663/101117352/101104661/101103238/101105996/101104302/101107268/101104186/100885770/101106871 | 13 |

**Supplementary Table 11 |** GO enrichment analysis of 96 rumen specifically expressed genes co-expressed with esophagus. BP denotes Biological Process, MF denotes Molecular Function, and CC denotes Cellular Component.

| ID | Type | Description | *p* value | adjusted *p* value | list1InGO | list1OutGO | list2InGO | list2OutGO |
|---|---|---|---|---|---|---|---|---|
| GO:0050786 | MF | RAGE receptor binding | 6.70E-34 | 1.14E-29 | 3 | 62 | 5 | 12528 |
| GO:0009913 | BP | epidermal cell differentiation | 2.21E-29 | 3.77E-25 | 6 | 59 | 37 | 12496 |
| GO:0050543 | MF | icosatetraenoic acid binding | 5.38E-25 | 9.16E-21 | 2 | 63 | 2 | 12531 |
| GO:0050544 | MF | arachidonic acid binding | 5.38E-25 | 9.16E-21 | 2 | 63 | 2 | 12531 |
| GO:0070488 | BP | neutrophil aggregation | 5.38E-25 | 9.16E-21 | 2 | 63 | 2 | 12531 |
| GO:0030216 | BP | keratinocyte differentiation | 1.88E-24 | 3.21E-20 | 5 | 60 | 30 | 12503 |
| GO:0035662 | MF | Toll-like receptor 4 binding | 3.45E-20 | 5.88E-16 | 2 | 63 | 3 | 12530 |
| GO:0001533 | CC | cornified envelope | 2.41E-18 | 4.11E-14 | 3 | 62 | 12 | 12521 |
| GO:0017014 | BP | protein nitrosylation | 5.62E-17 | 9.57E-13 | 2 | 63 | 4 | 12529 |
| GO:0018119 | BP | peptidyl-cysteine S-nitrosylation | 5.62E-17 | 9.57E-13 | 2 | 63 | 4 | 12529 |
| GO:0008236 | MF | serine-type peptidase activity | 5.72E-13 | 9.74E-09 | 7 | 58 | 122 | 12411 |
| GO:0035325 | MF | Toll-like receptor binding | 5.98E-13 | 1.02E-08 | 2 | 63 | 6 | 12527 |
| GO:0017171 | MF | serine hydrolase activity | 7.33E-13 | 1.25E-08 | 7 | 58 | 123 | 12410 |
| GO:0036041 | MF | long-chain fatty acid binding | 1.33E-11 | 2.27E-07 | 2 | 63 | 7 | 12526 |
| GO:0070486 | BP | leukocyte aggregation | 1.33E-11 | 2.27E-07 | 2 | 63 | 7 | 12526 |
| GO:0002523 | BP | leukocyte migration involved in inflammatory response | 1.60E-10 | 2.72E-06 | 2 | 63 | 8 | 12525 |
| GO:0004252 | MF | serine-type endopeptidase activity | 2.71E-10 | 4.62E-06 | 6 | 59 | 112 | 12421 |
| GO:0030676 | MF | Rac guanyl-nucleotide exchange factor activity | 1.23E-09 | 2.09E-05 | 2 | 63 | 9 | 12524 |
| GO:0030855 | BP | epithelial cell differentiation | 5.57E-09 | 9.49E-05 | 6 | 59 | 128 | 12405 |
| GO:0030414 | MF | peptidase inhibitor activity | 5.08E-07 | 0.008645 | 5 | 60 | 113 | 12420 |

| | | | | | | | | |
|---|---|---|---|---|---|---|---|---|
| GO:0016755 | MF | transferase activity, transferring amino-acyl groups | 7.45E-07 | 0.012689 | 2 | 63 | 14 | 12519 |
| GO:0018198 | BP | peptidyl-cysteine modification | 7.45E-07 | 0.012689 | 2 | 63 | 14 | 12519 |
| GO:0052547 | BP | regulation of peptidase activity | 1.17E-06 | 0.020009 | 7 | 58 | 228 | 12305 |
| GO:0000103 | BP | sulfate assimilation | 1.34E-06 | 0.022865 | 1 | 64 | 1 | 12532 |
| GO:0002785 | BP | negative regulation of antimicrobial peptide production | 1.34E-06 | 0.022865 | 1 | 64 | 1 | 12532 |
| GO:0002787 | BP | negative regulation of antibacterial peptide production | 1.34E-06 | 0.022865 | 1 | 64 | 1 | 12532 |
| GO:0035606 | BP | peptidyl-cysteine S-trans-nitrosylation | 1.34E-06 | 0.022865 | 1 | 64 | 1 | 12532 |
| GO:0042608 | MF | T cell receptor binding | 1.34E-06 | 0.022865 | 1 | 64 | 1 | 12532 |
| GO:1902572 | BP | negative regulation of serine-type peptidase activity | 1.34E-06 | 0.022865 | 1 | 64 | 1 | 12532 |
| GO:0006914 | BP | autophagy | 2.79E-06 | 0.047491 | 4 | 61 | 80 | 12453 |
| GO:0044421 | CC | extracellular region part | 2.92E-06 | 0.049783 | 27 | 38 | 2287 | 10246 |

**Supplementary Table 12 |** GO enrichment analysis of 88 rumen specifically expressed genes co-expressed with keratinization-associated tissues. BP denotes Biological Process, MF denotes Molecular Function, and CC denotes Cellular Component.

| ID | Type | Description | *p* value | adjusted *p* value | list1InGO | list1OutGO | list2InGO | list2OutGO |
|---|---|---|---|---|---|---|---|---|
| GO:0004867 | MF | serine-type endopeptidase inhibitor activity | 1.11E-18 | 1.89E-14 | 4 | 27 | 55 | 12478 |
| GO:0005149 | MF | interleukin-1 receptor binding | 2.20E-16 | 3.75E-12 | 2 | 29 | 11 | 12522 |
| GO:0003409 | BP | optic cup structural organization | 1.71E-12 | 2.90E-08 | 1 | 30 | 1 | 12532 |
| GO:0048144 | BP | fibroblast proliferation | 1.71E-12 | 2.90E-08 | 1 | 30 | 1 | 12532 |
| GO:0060648 | BP | mammary gland bud morphogenesis | 1.71E-12 | 2.90E-08 | 1 | 30 | 1 | 12532 |
| GO:0031069 | BP | hair follicle morphogenesis | 1.76E-11 | 3.00E-07 | 2 | 29 | 17 | 12516 |
| GO:0005615 | CC | extracellular space | 1.05E-10 | 1.79E-06 | 11 | 20 | 757 | 11776 |
| GO:0042481 | BP | regulation of odontogenesis | 1.83E-10 | 3.11E-06 | 2 | 29 | 19 | 12514 |
| GO:0004866 | MF | endopeptidase inhibitor activity | 3.00E-10 | 5.11E-06 | 4 | 27 | 104 | 12429 |
| GO:0061135 | MF | endopeptidase regulator activity | 5.66E-10 | 9.64E-06 | 4 | 27 | 107 | 12426 |
| GO:0030414 | MF | peptidase inhibitor activity | 1.83E-09 | 3.12E-05 | 4 | 27 | 113 | 12420 |
| GO:0001632 | MF | leukotriene B4 receptor activity | 9.85E-09 | 0.000168 | 1 | 30 | 2 | 12531 |
| GO:0002528 | BP | regulation of vascular permeability involved in acute inflammatory response | 9.85E-09 | 0.000168 | 1 | 30 | 2 | 12531 |
| GO:0003404 | BP | optic vesicle morphogenesis | 9.85E-09 | 0.000168 | 1 | 30 | 2 | 12531 |
| GO:0005152 | MF | interleukin-1 receptor antagonist activity | 9.85E-09 | 0.000168 | 1 | 30 | 2 | 12531 |
| GO:0019732 | BP | antifungal humoral response | 9.85E-09 | 0.000168 | 1 | 30 | 2 | 12531 |
| GO:0021623 | BP | oculomotor nerve formation | 9.85E-09 | 0.000168 | 1 | 30 | 2 | 12531 |
| GO:0061134 | MF | peptidase regulator activity | 3.32E-08 | 0.000565 | 4 | 27 | 131 | 12402 |
| GO:0010951 | BP | negative regulation of endopeptidase activity | 5.00E-08 | 0.000852 | 4 | 27 | 134 | 12399 |
| GO:0005109 | MF | frizzled binding | 1.49E-07 | 0.002535 | 2 | 29 | 28 | 12505 |
| GO:0010466 | BP | negative regulation of peptidase activity | 1.96E-07 | 0.003333 | 4 | 27 | 145 | 12388 |

| | | | | | | | | |
|---|---|---|---|---|---|---|---|---|
| GO:0043392 | BP | negative regulation of DNA binding | 2.47E-07 | 0.00421 | 2 | 29 | 29 | 12504 |
| GO:0045095 | CC | keratin filament | 2.47E-07 | 0.00421 | 2 | 29 | 29 | 12504 |
| GO:0005882 | CC | intermediate filament | 3.49E-07 | 0.005941 | 3 | 28 | 80 | 12453 |
| GO:0021871 | BP | forebrain regionalization | 7.79E-07 | 0.013273 | 1 | 30 | 3 | 12530 |
| GO:0060633 | BP | negative regulation of transcription initiation from RNA polymerase II promoter | 7.79E-07 | 0.013273 | 1 | 30 | 3 | 12530 |
| GO:0044464 | CC | cell part | 1.92E-06 | 0.032627 | 13 | 18 | 9861 | 2672 |

**Supplementary Table 13 |** GO enrichment analysis of 24 rumen specifically expressed genes co-expressed with liver. BP denotes Biological Process, MF denotes Molecular Function, and CC denotes Cellular Component.

| ID | Type | Description | *p* value | adjusted *p* value | list1InGO | list1OutGO | list2InGO | list2OutGO |
|---|---|---|---|---|---|---|---|---|
| GO:0009064 | BP | glutamine family amino acid metabolic process | 2.57E-25 | 4.37E-21 | 3 | 14 | 38 | 12495 |
| GO:0019752 | BP | carboxylic acid metabolic process | 3.35E-24 | 5.70E-20 | 8 | 9 | 355 | 12178 |
| GO:0043436 | BP | oxoacid metabolic process | 3.35E-24 | 5.70E-20 | 8 | 9 | 355 | 12178 |
| GO:0006082 | BP | organic acid metabolic process | 1.35E-23 | 2.31E-19 | 8 | 9 | 364 | 12169 |
| GO:0032787 | BP | monocarboxylic acid metabolic process | 3.41E-23 | 5.80E-19 | 6 | 11 | 202 | 12331 |
| GO:0042180 | BP | cellular ketone metabolic process | 5.12E-23 | 8.73E-19 | 8 | 9 | 373 | 12160 |
| GO:0004008 | MF | copper-exporting ATPase activity | 1.16E-21 | 1.98E-17 | 1 | 16 | 1 | 12532 |
| GO:0004397 | MF | histidine ammonia-lyase activity | 1.16E-21 | 1.98E-17 | 1 | 16 | 1 | 12532 |
| GO:0008887 | MF | glycerate kinase activity | 1.16E-21 | 1.98E-17 | 1 | 16 | 1 | 12532 |
| GO:0009812 | BP | flavonoid metabolic process | 1.16E-21 | 1.98E-17 | 1 | 16 | 1 | 12532 |
| GO:0015680 | BP | intracellular copper ion transport | 1.16E-21 | 1.98E-17 | 1 | 16 | 1 | 12532 |
| GO:0043682 | MF | copper-transporting ATPase activity | 1.16E-21 | 1.98E-17 | 1 | 16 | 1 | 12532 |
| GO:0051552 | BP | flavone metabolic process | 1.16E-21 | 1.98E-17 | 1 | 16 | 1 | 12532 |
| GO:0052696 | BP | flavonoid glucuronidation | 1.16E-21 | 1.98E-17 | 1 | 16 | 1 | 12532 |
| GO:0052697 | BP | xenobiotic glucuronidation | 1.16E-21 | 1.98E-17 | 1 | 16 | 1 | 12532 |
| GO:0060003 | BP | copper ion export | 1.16E-21 | 1.98E-17 | 1 | 16 | 1 | 12532 |
| GO:1990349 | BP | gap junction-mediated intercellular transport | 1.16E-21 | 1.98E-17 | 1 | 16 | 1 | 12532 |
| GO:0006637 | BP | acyl-CoA metabolic process | 2.37E-18 | 4.03E-14 | 2 | 15 | 19 | 12514 |
| GO:0035383 | BP | thioester metabolic process | 2.37E-18 | 4.03E-14 | 2 | 15 | 19 | 12514 |
| GO:0072562 | CC | blood microparticle | 4.52E-18 | 7.71E-14 | 3 | 14 | 55 | 12478 |

| | | | | | | | | |
|---|---|---|---|---|---|---|---|---|
| GO:0004053 | MF | arginase activity | 6.92E-15 | 1.18E-10 | 1 | 16 | 2 | 12531 |
| GO:0004067 | MF | asparaginase activity | 6.92E-15 | 1.18E-10 | 1 | 16 | 2 | 12531 |
| GO:0006530 | BP | asparagine catabolic process | 6.92E-15 | 1.18E-10 | 1 | 16 | 2 | 12531 |
| GO:0009698 | BP | phenylpropanoid metabolic process | 6.92E-15 | 1.18E-10 | 1 | 16 | 2 | 12531 |
| GO:0015942 | BP | formate metabolic process | 6.92E-15 | 1.18E-10 | 1 | 16 | 2 | 12531 |
| GO:0019556 | BP | histidine catabolic process to glutamate and formamide | 6.92E-15 | 1.18E-10 | 1 | 16 | 2 | 12531 |
| GO:0019557 | BP | histidine catabolic process to glutamate and formate | 6.92E-15 | 1.18E-10 | 1 | 16 | 2 | 12531 |
| GO:0043152 | BP | induction of bacterial agglutination | 6.92E-15 | 1.18E-10 | 1 | 16 | 2 | 12531 |
| GO:0051208 | BP | sequestering of calcium ion | 6.92E-15 | 1.18E-10 | 1 | 16 | 2 | 12531 |
| GO:0052695 | BP | cellular glucuronidation | 6.92E-15 | 1.18E-10 | 1 | 16 | 2 | 12531 |
| GO:0060481 | BP | lobar bronchus epithelium development | 6.92E-15 | 1.18E-10 | 1 | 16 | 2 | 12531 |
| GO:0000050 | BP | urea cycle | 1.76E-11 | 3.00E-07 | 1 | 16 | 3 | 12530 |
| GO:0005375 | MF | copper ion transmembrane transporter activity | 1.76E-11 | 3.00E-07 | 1 | 16 | 3 | 12530 |
| GO:0016841 | MF | ammonia-lyase activity | 1.76E-11 | 3.00E-07 | 1 | 16 | 3 | 12530 |
| GO:0019627 | BP | urea metabolic process | 1.76E-11 | 3.00E-07 | 1 | 16 | 3 | 12530 |
| GO:0050436 | MF | microfibril binding | 1.76E-11 | 3.00E-07 | 1 | 16 | 3 | 12530 |
| GO:0044281 | BP | small molecule metabolic process | 8.41E-11 | 1.43E-06 | 8 | 9 | 769 | 11764 |
| GO:0003996 | MF | acyl-CoA ligase activity | 1.99E-09 | 3.39E-05 | 1 | 16 | 4 | 12529 |
| GO:0004022 | MF | alcohol dehydrogenase (NAD) activity | 1.99E-09 | 3.39E-05 | 1 | 16 | 4 | 12529 |
| GO:0004321 | MF | fatty-acyl-CoA synthase activity | 1.99E-09 | 3.39E-05 | 1 | 16 | 4 | 12529 |
| GO:0006063 | BP | uronic acid metabolic process | 1.99E-09 | 3.39E-05 | 1 | 16 | 4 | 12529 |
| GO:0006068 | BP | ethanol catabolic process | 1.99E-09 | 3.39E-05 | 1 | 16 | 4 | 12529 |
| GO:0006528 | BP | asparagine metabolic process | 1.99E-09 | 3.39E-05 | 1 | 16 | 4 | 12529 |
| GO:0006547 | BP | histidine metabolic process | 1.99E-09 | 3.39E-05 | 1 | 16 | 4 | 12529 |
| GO:0006548 | BP | histidine catabolic process | 1.99E-09 | 3.39E-05 | 1 | 16 | 4 | 12529 |

| | | | | | | | | |
|---|---|---|---|---|---|---|---|---|
| GO:0009075 | BP | obsolete histidine family amino acid metabolic process | 1.99E-09 | 3.39E-05 | 1 | 16 | 4 | 12529 |
| GO:0009077 | BP | obsolete histidine family amino acid catabolic process | 1.99E-09 | 3.39E-05 | 1 | 16 | 4 | 12529 |
| GO:0015677 | BP | copper ion import | 1.99E-09 | 3.39E-05 | 1 | 16 | 4 | 12529 |
| GO:0019585 | BP | glucuronate metabolic process | 1.99E-09 | 3.39E-05 | 1 | 16 | 4 | 12529 |
| GO:0034310 | BP | primary alcohol catabolic process | 1.99E-09 | 3.39E-05 | 1 | 16 | 4 | 12529 |
| GO:0071941 | BP | nitrogen cycle metabolic process | 1.99E-09 | 3.39E-05 | 1 | 16 | 4 | 12529 |
| GO:0072378 | BP | blood coagulation, fibrin clot formation | 1.99E-09 | 3.39E-05 | 1 | 16 | 4 | 12529 |
| GO:0034754 | BP | cellular hormone metabolic process | 2.12E-09 | 3.61E-05 | 2 | 15 | 41 | 12492 |
| GO:0044282 | BP | small molecule catabolic process | 6.54E-09 | 0.000111 | 3 | 14 | 118 | 12415 |
| GO:0044712 | BP | single-organism catabolic process | 6.54E-09 | 0.000111 | 3 | 14 | 118 | 12415 |
| GO:0009063 | BP | cellular amino acid catabolic process | 2.42E-08 | 0.000411 | 2 | 15 | 47 | 12486 |
| GO:0006067 | BP | ethanol metabolic process | 4.73E-08 | 0.000806 | 1 | 16 | 5 | 12528 |
| GO:0006699 | BP | bile acid biosynthetic process | 4.73E-08 | 0.000806 | 1 | 16 | 5 | 12528 |
| GO:0016840 | MF | carbon-nitrogen lyase activity | 4.73E-08 | 0.000806 | 1 | 16 | 5 | 12528 |
| GO:0034308 | BP | primary alcohol metabolic process | 4.73E-08 | 0.000806 | 1 | 16 | 5 | 12528 |
| GO:0035434 | BP | copper ion transmembrane transport | 4.73E-08 | 0.000806 | 1 | 16 | 5 | 12528 |
| GO:0043604 | BP | amide biosynthetic process | 4.73E-08 | 0.000806 | 1 | 16 | 5 | 12528 |
| GO:0051238 | BP | sequestering of metal ion | 4.73E-08 | 0.000806 | 1 | 16 | 5 | 12528 |
| GO:0006520 | BP | cellular amino acid metabolic process | 9.36E-08 | 0.001594 | 3 | 14 | 137 | 12396 |
| GO:0006707 | BP | cholesterol catabolic process | 4.60E-07 | 0.007839 | 1 | 16 | 6 | 12527 |
| GO:0016127 | BP | sterol catabolic process | 4.60E-07 | 0.007839 | 1 | 16 | 6 | 12527 |
| GO:0005577 | CC | fibrinogen complex | 2.55E-06 | 0.043519 | 1 | 16 | 7 | 12526 |
| GO:0006525 | BP | arginine metabolic process | 2.55E-06 | 0.043519 | 1 | 16 | 7 | 12526 |
| GO:0006878 | BP | cellular copper ion homeostasis | 2.55E-06 | 0.043519 | 1 | 16 | 7 | 12526 |
| GO:0009068 | BP | aspartate family amino acid catabolic process | 2.55E-06 | 0.043519 | 1 | 16 | 7 | 12526 |

| | | | | | | | | |
|---|---|---|---|---|---|---|---|---|
| GO:0019731 | BP | antibacterial humoral response | 2.55E-06 | 0.043519 | 1 | 16 | 7 | 12526 |
| GO:0033574 | BP | response to testosterone | 2.55E-06 | 0.043519 | 1 | 16 | 7 | 12526 |

**Supplementary Table 14 |** GO enrichment analysis of 61 rumen specifically expressed genes co-expressed with intestine. The intestine includes duodenum, jejunum, ileum, caecum, colon, and rectum. BP denotes Biological Process, MF denotes Molecular Function, and CC denotes Cellular Component.

| ID | Type | Description | *p* value | adjusted *p* value | list1InGO | list1OutGO | list2InGO | list2OutGO |
|---|---|---|---|---|---|---|---|---|
| GO:0015386 | MF | potassium:proton antiporter activity | 2.13E-22 | 3.62E-18 | 2 | 27 | 8 | 12525 |
| GO:0098719 | BP | sodium ion import across plasma membrane | 1.89E-20 | 3.22E-16 | 2 | 27 | 9 | 12524 |
| GO:0005451 | MF | monovalent cation:proton antiporter activity | 7.99E-19 | 1.36E-14 | 2 | 27 | 10 | 12523 |
| GO:0015385 | MF | sodium:proton antiporter activity | 7.99E-19 | 1.36E-14 | 2 | 27 | 10 | 12523 |
| GO:0015299 | MF | solute:proton antiporter activity | 2.89E-16 | 4.92E-12 | 2 | 27 | 12 | 12521 |
| GO:0022821 | MF | potassium ion antiporter activity | 2.89E-16 | 4.92E-12 | 2 | 27 | 12 | 12521 |
| GO:0051453 | BP | regulation of intracellular pH | 1.00E-14 | 1.70E-10 | 3 | 26 | 39 | 12494 |
| GO:0030641 | BP | regulation of cellular pH | 4.44E-14 | 7.57E-10 | 3 | 26 | 41 | 12492 |
| GO:0032002 | CC | interleukin-28 receptor complex | 2.89E-13 | 4.92E-09 | 1 | 28 | 1 | 12532 |
| GO:0033842 | MF | N-acetyl-beta-glucosaminyl-glycoprotein 4-beta-N-acetylgalactosaminyltransferase activity | 2.89E-13 | 4.92E-09 | 1 | 28 | 1 | 12532 |
| GO:0043163 | BP | cell envelope organization | 2.89E-13 | 4.92E-09 | 1 | 28 | 1 | 12532 |
| GO:0045229 | BP | external encapsulating structure organization | 2.89E-13 | 4.92E-09 | 1 | 28 | 1 | 12532 |
| GO:0047498 | MF | calcium-dependent phospholipase A2 activity | 2.89E-13 | 4.92E-09 | 1 | 28 | 1 | 12532 |
| GO:1904220 | BP | regulation of serine C-palmitoyltransferase activity | 2.89E-13 | 4.92E-09 | 1 | 28 | 1 | 12532 |
| GO:0006885 | BP | regulation of pH | 1.91E-12 | 3.25E-08 | 3 | 26 | 47 | 12486 |
| GO:0030004 | BP | cellular monovalent inorganic cation homeostasis | 9.13E-12 | 1.55E-07 | 3 | 26 | 50 | 12483 |
| GO:0015297 | MF | antiporter activity | 1.30E-10 | 2.21E-06 | 3 | 26 | 56 | 12477 |
| GO:0015491 | MF | cation:cation antiporter activity | 3.13E-10 | 5.33E-06 | 2 | 27 | 21 | 12512 |
| GO:0055067 | BP | monovalent inorganic cation homeostasis | 1.57E-09 | 2.67E-05 | 3 | 26 | 63 | 12470 |
| GO:0015298 | MF | solute:cation antiporter activity | 1.78E-09 | 3.04E-05 | 2 | 27 | 23 | 12510 |

| | | | | | | | | |
|---|---|---|---|---|---|---|---|---|
| GO:0001951 | BP | intestinal D-glucose absorption | 2.98E-09 | 5.08E-05 | 1 | 28 | 2 | 12531 |
| GO:0004528 | MF | phosphodiesterase I activity | 2.98E-09 | 5.08E-05 | 1 | 28 | 2 | 12531 |
| GO:0018146 | BP | keratan sulfate biosynthetic process | 2.98E-09 | 5.08E-05 | 1 | 28 | 2 | 12531 |
| GO:0018149 | BP | peptide cross-linking | 7.86E-09 | 0.000134 | 2 | 27 | 25 | 12508 |
| GO:0015291 | MF | secondary active transmembrane transporter activity | 1.60E-08 | 0.000273 | 4 | 25 | 135 | 12398 |
| GO:0006814 | BP | sodium ion transport | 3.18E-08 | 0.000541 | 3 | 26 | 74 | 12459 |
| GO:0030216 | BP | keratinocyte differentiation | 1.44E-07 | 0.002456 | 2 | 27 | 30 | 12503 |
| GO:0042339 | BP | keratan sulfate metabolic process | 3.16E-07 | 0.005377 | 1 | 28 | 3 | 12530 |
| GO:0045852 | BP | pH elevation | 3.16E-07 | 0.005377 | 1 | 28 | 3 | 12530 |
| GO:0047035 | MF | testosterone dehydrogenase (NAD+) activity | 3.16E-07 | 0.005377 | 1 | 28 | 3 | 12530 |
| GO:0051454 | BP | intracellular pH elevation | 3.16E-07 | 0.005377 | 1 | 28 | 3 | 12530 |
| GO:0015081 | MF | sodium ion transmembrane transporter activity | 9.87E-07 | 0.01681 | 3 | 26 | 92 | 12441 |
| GO:0005044 | MF | scavenger receptor activity | 1.74E-06 | 0.029712 | 2 | 27 | 36 | 12497 |
| GO:0009913 | BP | epidermal cell differentiation | 2.46E-06 | 0.041829 | 2 | 27 | 37 | 12496 |
| GO:1902600 | BP | hydrogen ion transmembrane transport | 2.46E-06 | 0.041829 | 2 | 27 | 37 | 12496 |

**Supplementary Table 15 |** GO enrichment analysis of 23 rumen specifically expressed genes co-expressed with muscle. BP denotes Biological Process, MF denotes Molecular Function, and CC denotes Cellular Component.

| ID | Type | Description | *p* value | adjusted *p* value | list1InGO | list1OutGO | list2InGO | list2OutGO |
|---|---|---|---|---|---|---|---|---|
| GO:0000272 | BP | polysaccharide catabolic process | 2.55E-06 | 0.043519 | 1 | 16 | 7 | 12526 |
| GO:0001957 | BP | intramembranous ossification | 1.99E-09 | 3.39E-05 | 1 | 16 | 4 | 12529 |
| GO:0003334 | BP | keratinocyte development | 4.73E-08 | 0.000806 | 1 | 16 | 5 | 12528 |
| GO:0003989 | MF | acetyl-CoA carboxylase activity | 6.92E-15 | 1.18E-10 | 1 | 16 | 2 | 12531 |
| GO:0004075 | MF | biotin carboxylase activity | 4.73E-08 | 0.000806 | 1 | 16 | 5 | 12528 |
| GO:0005546 | MF | phosphatidylinositol-4,5-bisphosphate binding | 2.38E-12 | 4.05E-08 | 2 | 15 | 30 | 12503 |
| GO:0005861 | CC | troponin complex | 4.73E-08 | 0.000806 | 1 | 16 | 5 | 12528 |
| GO:0005865 | CC | striated muscle thin filament | 6.92E-15 | 1.18E-10 | 1 | 16 | 2 | 12531 |
| GO:0005884 | CC | actin filament | 1.13E-11 | 1.93E-07 | 2 | 15 | 32 | 12501 |
| GO:0005980 | BP | glycogen catabolic process | 2.55E-06 | 0.043519 | 1 | 16 | 7 | 12526 |
| GO:0006937 | BP | regulation of muscle contraction | 1.19E-06 | 0.020187 | 2 | 15 | 61 | 12472 |
| GO:0009251 | BP | glucan catabolic process | 2.55E-06 | 0.043519 | 1 | 16 | 7 | 12526 |
| GO:0010314 | MF | phosphatidylinositol-5-phosphate binding | 2.55E-06 | 0.043519 | 1 | 16 | 7 | 12526 |
| GO:0010614 | BP | negative regulation of cardiac muscle hypertrophy | 4.60E-07 | 0.007839 | 1 | 16 | 6 | 12527 |
| GO:0010830 | BP | regulation of myotube differentiation | 1.47E-17 | 2.50E-13 | 2 | 15 | 20 | 12513 |
| GO:0010831 | BP | positive regulation of myotube differentiation | 4.60E-07 | 0.007839 | 1 | 16 | 6 | 12527 |
| GO:0010861 | MF | thyroid hormone receptor activator activity | 6.92E-15 | 1.18E-10 | 1 | 16 | 2 | 12531 |
| GO:0010959 | BP | regulation of metal ion transport | 1.88E-07 | 0.003203 | 3 | 14 | 143 | 12390 |
| GO:0014741 | BP | negative regulation of muscle hypertrophy | 4.60E-07 | 0.007839 | 1 | 16 | 6 | 12527 |
| GO:0016202 | BP | regulation of striated muscle tissue development | 2.65E-18 | 4.52E-14 | 4 | 13 | 106 | 12427 |

| | | | | | | | | |
|---|---|---|---|---|---|---|---|---|
| GO:0016421 | MF | CoA carboxylase activity | 4.60E-07 | 0.007839 | 1 | 16 | 6 | 12527 |
| GO:0022898 | BP | regulation of transmembrane transporter activity | 3.02E-13 | 5.15E-09 | 3 | 14 | 77 | 12456 |
| GO:0030018 | CC | Z disc | 9.51E-07 | 0.01619 | 2 | 15 | 60 | 12473 |
| GO:0030029 | BP | actin filament-based process | 4.20E-12 | 7.15E-08 | 5 | 12 | 264 | 12269 |
| GO:0030035 | BP | microspike assembly | 6.92E-15 | 1.18E-10 | 1 | 16 | 2 | 12531 |
| GO:0030036 | BP | actin cytoskeleton organization | 1.58E-12 | 2.69E-08 | 5 | 12 | 255 | 12278 |
| GO:0030175 | CC | filopodium | 4.08E-13 | 6.95E-09 | 2 | 15 | 28 | 12505 |
| GO:0030240 | BP | skeletal muscle thin filament assembly | 1.76E-11 | 3.00E-07 | 1 | 16 | 3 | 12530 |
| GO:0030274 | MF | LIM domain binding | 4.73E-08 | 0.000806 | 1 | 16 | 5 | 12528 |
| GO:0030314 | CC | junctional membrane complex | 4.60E-07 | 0.007839 | 1 | 16 | 6 | 12527 |
| GO:0030315 | CC | T-tubule | 2.65E-25 | 4.51E-21 | 2 | 15 | 13 | 12520 |
| GO:0030375 | MF | thyroid hormone receptor coactivator activity | 6.92E-15 | 1.18E-10 | 1 | 16 | 2 | 12531 |
| GO:0031032 | BP | actomyosin structure organization | 4.69E-07 | 0.007992 | 2 | 15 | 57 | 12476 |
| GO:0031432 | MF | titin binding | 4.73E-08 | 0.000806 | 1 | 16 | 5 | 12528 |
| GO:0031998 | BP | regulation of fatty acid beta-oxidation | 1.99E-09 | 3.39E-05 | 1 | 16 | 4 | 12529 |
| GO:0031999 | BP | negative regulation of fatty acid beta-oxidation | 6.92E-15 | 1.18E-10 | 1 | 16 | 2 | 12531 |
| GO:0032409 | BP | regulation of transporter activity | 4.37E-12 | 7.44E-08 | 3 | 14 | 85 | 12448 |
| GO:0032411 | BP | positive regulation of transporter activity | 1.01E-12 | 1.73E-08 | 2 | 15 | 29 | 12504 |
| GO:0032412 | BP | regulation of ion transmembrane transporter activity | 4.27E-14 | 7.28E-10 | 3 | 14 | 72 | 12461 |
| GO:0032414 | BP | positive regulation of ion transmembrane transporter activity | 1.46E-15 | 2.48E-11 | 2 | 15 | 23 | 12510 |
| GO:0032769 | BP | negative regulation of monooxygenase activity | 4.60E-07 | 0.007839 | 1 | 16 | 6 | 12527 |
| GO:0035914 | BP | skeletal muscle cell differentiation | 2.12E-09 | 3.61E-05 | 2 | 15 | 41 | 12492 |
| GO:0035995 | BP | detection of muscle stretch | 1.76E-11 | 3.00E-07 | 1 | 16 | 3 | 12530 |
| GO:0036072 | BP | direct ossification | 1.99E-09 | 3.39E-05 | 1 | 16 | 4 | 12529 |
| GO:0038009 | BP | regulation of signal transduction by receptor internalization | 1.16E-21 | 1.98E-17 | 1 | 16 | 1 | 12532 |

| | | | | | | | | |
|---|---|---|---|---|---|---|---|---|
| GO:0043268 | BP | positive regulation of potassium ion transport | 2.55E-06 | 0.043519 | 1 | 16 | 7 | 12526 |
| GO:0043271 | BP | negative regulation of ion transport | 1.01E-12 | 1.73E-08 | 2 | 15 | 29 | 12504 |
| GO:0043502 | BP | regulation of muscle adaptation | 4.08E-13 | 6.95E-09 | 2 | 15 | 28 | 12505 |
| GO:0044247 | BP | cellular polysaccharide catabolic process | 2.55E-06 | 0.043519 | 1 | 16 | 7 | 12526 |
| GO:0044325 | MF | ion channel binding | 1.19E-06 | 0.020187 | 2 | 15 | 61 | 12472 |
| GO:0044449 | CC | contractile fiber part | 5.02E-30 | 8.55E-26 | 5 | 12 | 104 | 12429 |
| GO:0045214 | BP | sarcomere organization | 1.54E-13 | 2.63E-09 | 2 | 15 | 27 | 12506 |
| GO:0045792 | BP | negative regulation of cell size | 4.60E-07 | 0.007839 | 1 | 16 | 6 | 12527 |
| GO:0046322 | BP | negative regulation of fatty acid oxidation | 1.99E-09 | 3.39E-05 | 1 | 16 | 4 | 12529 |
| GO:0048468 | BP | cell development | 5.23E-08 | 0.00089 | 5 | 12 | 398 | 12135 |
| GO:0048634 | BP | regulation of muscle organ development | 5.58E-18 | 9.51E-14 | 4 | 13 | 108 | 12425 |
| GO:0048641 | BP | regulation of skeletal muscle tissue development | 7.34E-15 | 1.25E-10 | 3 | 14 | 68 | 12465 |
| GO:0048741 | BP | skeletal muscle fiber development | 3.19E-19 | 5.44E-15 | 2 | 15 | 18 | 12515 |
| GO:0048742 | BP | regulation of skeletal muscle fiber development | 2.23E-18 | 3.80E-14 | 3 | 14 | 54 | 12479 |
| GO:0048747 | BP | muscle fiber development | 4.53E-11 | 7.71E-07 | 2 | 15 | 34 | 12499 |
| GO:0051001 | BP | negative regulation of nitric-oxide synthase activity | 1.76E-11 | 3.00E-07 | 1 | 16 | 3 | 12530 |
| GO:0051147 | BP | regulation of muscle cell differentiation | 1.37E-10 | 2.33E-06 | 3 | 14 | 98 | 12435 |
| GO:0051153 | BP | regulation of striated muscle cell differentiation | 6.19E-13 | 1.05E-08 | 3 | 14 | 79 | 12454 |
| GO:0051289 | BP | protein homotetramerization | 2.12E-09 | 3.61E-05 | 2 | 15 | 41 | 12492 |
| GO:0051373 | MF | FATZ binding | 1.76E-11 | 3.00E-07 | 1 | 16 | 3 | 12530 |
| GO:0051394 | BP | regulation of nerve growth factor receptor activity | 1.16E-21 | 1.98E-17 | 1 | 16 | 1 | 12532 |
| GO:0051647 | BP | nucleus localization | 4.73E-08 | 0.000806 | 1 | 16 | 5 | 12528 |
| GO:0051695 | BP | actin filament uncapping | 6.92E-15 | 1.18E-10 | 1 | 16 | 2 | 12531 |
| GO:0055001 | BP | muscle cell development | 2.51E-33 | 4.27E-29 | 4 | 13 | 56 | 12477 |
| GO:0055002 | BP | striated muscle cell development | 8.57E-11 | 1.46E-06 | 2 | 15 | 35 | 12498 |

| | | | | | | | | |
|---|---|---|---|---|---|---|---|---|
| GO:0055006 | BP | cardiac cell development | 8.74E-24 | 1.49E-19 | 2 | 15 | 14 | 12519 |
| GO:0055013 | BP | cardiac muscle cell development | 8.74E-24 | 1.49E-19 | 2 | 15 | 14 | 12519 |
| GO:0060297 | BP | regulation of sarcomere organization | 4.73E-08 | 0.000806 | 1 | 16 | 5 | 12528 |
| GO:0060299 | BP | negative regulation of sarcomere organization | 1.16E-21 | 1.98E-17 | 1 | 16 | 1 | 12532 |
| GO:0060316 | BP | positive regulation of ryanodine-sensitive calcium-release channel activity | 6.92E-15 | 1.18E-10 | 1 | 16 | 2 | 12531 |
| GO:0060373 | BP | regulation of ventricular cardiac muscle cell membrane depolarization | 1.99E-09 | 3.39E-05 | 1 | 16 | 4 | 12529 |
| GO:0060421 | BP | positive regulation of heart growth | 1.99E-09 | 3.39E-05 | 1 | 16 | 4 | 12529 |
| GO:0060762 | BP | regulation of branching involved in mammary gland duct morphogenesis | 1.99E-09 | 3.39E-05 | 1 | 16 | 4 | 12529 |
| GO:0061723 | BP | glycophagy | 1.16E-21 | 1.98E-17 | 1 | 16 | 1 | 12532 |
| GO:0070080 | MF | titin Z domain binding | 6.92E-15 | 1.18E-10 | 1 | 16 | 2 | 12531 |
| GO:0070836 | BP | caveola assembly | 1.99E-09 | 3.39E-05 | 1 | 16 | 4 | 12529 |
| GO:0071253 | MF | connexin binding | 4.73E-08 | 0.000806 | 1 | 16 | 5 | 12528 |
| GO:0086097 | BP | phospholipase C-activating angiotensin-activated signaling pathway | 1.16E-21 | 1.98E-17 | 1 | 16 | 1 | 12532 |
| GO:0090130 | BP | tissue migration | 1.99E-09 | 3.39E-05 | 1 | 16 | 4 | 12529 |
| GO:0090131 | BP | mesenchyme migration | 1.16E-21 | 1.98E-17 | 1 | 16 | 1 | 12532 |
| GO:0090150 | BP | establishment of protein localization to membrane | 3.42E-08 | 0.000583 | 2 | 15 | 48 | 12485 |
| GO:0098909 | BP | regulation of cardiac muscle cell action potential involved in regulation of contraction | 1.76E-11 | 3.00E-07 | 1 | 16 | 3 | 12530 |
| GO:1900825 | BP | regulation of membrane depolarization during cardiac muscle cell action potential | 1.99E-09 | 3.39E-05 | 1 | 16 | 4 | 12529 |
| GO:1900826 | BP | negative regulation of membrane depolarization during cardiac muscle cell action potential | 1.16E-21 | 1.98E-17 | 1 | 16 | 1 | 12532 |
| GO:1901017 | BP | negative regulation of potassium ion transmembrane transporter activity | 1.99E-09 | 3.39E-05 | 1 | 16 | 4 | 12529 |
| GO:1901018 | BP | positive regulation of potassium ion transmembrane transporter activity | 4.73E-08 | 0.000806 | 1 | 16 | 5 | 12528 |
| GO:1901019 | BP | regulation of calcium ion transmembrane transporter activity | 3.57E-16 | 6.08E-12 | 2 | 15 | 22 | 12511 |
| GO:1901021 | BP | positive regulation of calcium ion transmembrane transporter activity | 2.55E-06 | 0.043519 | 1 | 16 | 7 | 12526 |
| GO:2000009 | BP | negative regulation of protein localization to cell surface | 4.60E-07 | 0.007839 | 1 | 16 | 6 | 12527 |
| GO:2001014 | BP | regulation of skeletal muscle cell differentiation | 2.35E-34 | 4.00E-30 | 3 | 14 | 27 | 12506 |

| | | | | | | | | |
|---|---|---|---|---|---|---|---|---|
| GO:2001069 | MF | glycogen binding | 1.76E-11 | 3.00E-07 | 1 | 16 | 3 | 12530 |
| GO:2001070 | MF | starch binding | 6.92E-15 | 1.18E-10 | 1 | 16 | 2 | 12531 |
| GO:2001135 | BP | regulation of endocytic recycling | 2.55E-06 | 0.043519 | 1 | 16 | 7 | 12526 |
| GO:2001137 | BP | positive regulation of endocytic recycling | 1.76E-11 | 3.00E-07 | 1 | 16 | 3 | 12530 |
| GO:2001257 | BP | regulation of cation channel activity | 1.54E-13 | 2.63E-09 | 2 | 15 | 27 | 12506 |
| GO:2001259 | BP | positive regulation of cation channel activity | 2.26E-31 | 3.86E-27 | 2 | 15 | 10 | 12523 |
| GO:2001293 | BP | malonyl-CoA metabolic process | 6.92E-15 | 1.18E-10 | 1 | 16 | 2 | 12531 |
| GO:2001295 | BP | malonyl-CoA biosynthetic process | 1.16E-21 | 1.98E-17 | 1 | 16 | 1 | 12532 |

**Supplementary Table 16 |** GO enrichment analysis of 19 rumen specifically expressed genes co-expressed with kidney. BP denotes Biological Process, MF denotes Molecular Function, and CC denotes Cellular Component.

| ID | Type | Description | *p* value | adjusted *p* value | list1InGO | list1OutGO | list2InGO | list2OutGO |
|---|---|---|---|---|---|---|---|---|
| GO:0008336 | MF | gamma-butyrobetaine dioxygenase activity | 7.07E-28 | 1.20E-23 | 1 | 12 | 1 | 12532 |
| GO:0015265 | MF | urea channel activity | 7.07E-28 | 1.20E-23 | 1 | 12 | 1 | 12532 |
| GO:0015204 | MF | urea transmembrane transporter activity | 4.76E-19 | 8.10E-15 | 1 | 12 | 2 | 12531 |
| GO:0042887 | MF | amide transmembrane transporter activity | 4.76E-19 | 8.10E-15 | 1 | 12 | 2 | 12531 |
| GO:0045329 | BP | carnitine biosynthetic process | 4.76E-19 | 8.10E-15 | 1 | 12 | 2 | 12531 |
| GO:0071918 | BP | urea transmembrane transport | 4.76E-19 | 8.10E-15 | 1 | 12 | 2 | 12531 |
| GO:0006578 | BP | amino-acid betaine biosynthetic process | 1.29E-14 | 2.19E-10 | 1 | 12 | 3 | 12530 |
| GO:0015321 | MF | sodium-dependent phosphate transmembrane transporter activity | 1.29E-14 | 2.19E-10 | 1 | 12 | 3 | 12530 |
| GO:0044341 | BP | sodium-dependent phosphate transport | 1.29E-14 | 2.19E-10 | 1 | 12 | 3 | 12530 |
| GO:0015840 | BP | urea transport | 6.01E-12 | 1.02E-07 | 1 | 12 | 4 | 12529 |
| GO:0016338 | BP | calcium-independent cell-cell adhesion via plasma membrane cell-adhesion molecules | 3.68E-10 | 6.26E-06 | 1 | 12 | 5 | 12528 |
| GO:0005436 | MF | sodium:phosphate symporter activity | 7.03E-09 | 0.0001197 | 1 | 12 | 6 | 12527 |
| GO:0009437 | BP | carnitine metabolic process | 7.03E-09 | 0.0001197 | 1 | 12 | 6 | 12527 |
| GO:0015238 | MF | drug transmembrane transporter activity | 7.03E-09 | 0.0001197 | 1 | 12 | 6 | 12527 |
| GO:0030643 | BP | cellular phosphate ion homeostasis | 7.03E-09 | 0.0001197 | 1 | 12 | 6 | 12527 |
| GO:0042886 | BP | amide transport | 7.03E-09 | 0.0001197 | 1 | 12 | 6 | 12527 |
| GO:0048012 | BP | hepatocyte growth factor receptor signaling pathway | 7.03E-09 | 0.0001197 | 1 | 12 | 6 | 12527 |
| GO:0072502 | BP | cellular trivalent inorganic anion homeostasis | 7.03E-09 | 0.0001197 | 1 | 12 | 6 | 12527 |
| GO:0006577 | BP | amino-acid betaine metabolic process | 6.47E-08 | 0.0011029 | 1 | 12 | 7 | 12526 |
| GO:0030002 | BP | cellular anion homeostasis | 6.47E-08 | 0.0011029 | 1 | 12 | 7 | 12526 |

| | | | | | | | | |
|---|---|---|---|---|---|---|---|---|
| GO:0055062 | BP | phosphate ion homeostasis | 3.67E-07 | 0.0062452 | 1 | 12 | 8 | 12525 |
| GO:0072506 | BP | trivalent inorganic anion homeostasis | 3.67E-07 | 0.0062452 | 1 | 12 | 8 | 12525 |
| GO:0015698 | BP | inorganic anion transport | 1.10E-06 | 0.0187293 | 2 | 11 | 80 | 12453 |
| GO:0006855 | BP | drug transmembrane transport | 1.48E-06 | 0.0251308 | 1 | 12 | 9 | 12524 |

**Supplementary Table 17 |** GO enrichment analysis of 10 rumen specifically expressed genes co-expressed with salivary gland. BP denotes Biological Process, MF denotes Molecular Function, and CC denotes Cellular Component.

| ID | Type | Description | *p* value | adjusted *p* value | list1InGO | list1OutGO | list2InGO | list2OutGO |
|---|---|---|---|---|---|---|---|---|
| GO:0021589 | BP | cerebellum structural organization | 5.85E-70 | 9.97E-66 | 1 | 4 | 1 | 12532 |
| GO:0021813 | BP | cell-cell adhesion involved in neuronal-glial interactions involved in cerebral cortex radial glia guided migration | 5.85E-70 | 9.97E-66 | 1 | 4 | 1 | 12532 |
| GO:0030157 | BP | pancreatic juice secretion | 9.18E-36 | 1.56E-31 | 1 | 4 | 3 | 12530 |
| GO:0097477 | BP | lateral motor column neuron migration | 9.18E-36 | 1.56E-31 | 1 | 4 | 3 | 12530 |
| GO:0015670 | BP | carbon dioxide transport | 6.61E-29 | 1.13E-24 | 1 | 4 | 4 | 12529 |
| GO:0021942 | BP | radial glia guided migration of Purkinje cell | 6.61E-29 | 1.13E-24 | 1 | 4 | 4 | 12529 |
| GO:0051645 | BP | Golgi localization | 6.61E-29 | 1.13E-24 | 1 | 4 | 4 | 12529 |
| GO:0002279 | BP | mast cell activation involved in immune response | 2.51E-24 | 4.27E-20 | 1 | 4 | 5 | 12528 |
| GO:0021932 | BP | hindbrain radial glia guided cell migration | 2.51E-24 | 4.27E-20 | 1 | 4 | 5 | 12528 |
| GO:0043303 | BP | mast cell degranulation | 2.51E-24 | 4.27E-20 | 1 | 4 | 5 | 12528 |
| GO:0046541 | BP | saliva secretion | 2.51E-24 | 4.27E-20 | 1 | 4 | 5 | 12528 |
| GO:0021517 | BP | ventral spinal cord development | 4.72E-21 | 8.05E-17 | 1 | 4 | 6 | 12527 |
| GO:0007288 | BP | sperm axoneme assembly | 1.36E-18 | 2.32E-14 | 1 | 4 | 7 | 12526 |
| GO:0019372 | BP | lipoxygenase pathway | 1.36E-18 | 2.32E-14 | 1 | 4 | 7 | 12526 |
| GO:0021535 | BP | cell migration in hindbrain | 1.36E-18 | 2.32E-14 | 1 | 4 | 7 | 12526 |
| GO:0042629 | CC | mast cell granule | 1.36E-18 | 2.32E-14 | 1 | 4 | 7 | 12526 |
| GO:0048712 | BP | negative regulation of astrocyte differentiation | 1.36E-18 | 2.32E-14 | 1 | 4 | 7 | 12526 |
| GO:0001675 | BP | acrosome assembly | 1.13E-16 | 1.92E-12 | 1 | 4 | 8 | 12525 |
| GO:0005372 | MF | water transmembrane transporter activity | 1.13E-16 | 1.92E-12 | 1 | 4 | 8 | 12525 |

| | | | | | | | | |
|---|---|---|---|---|---|---|---|---|
| GO:0015250 | MF | water channel activity | 1.13E-16 | 1.92E-12 | 1 | 4 | 8 | 12525 |
| GO:0045576 | BP | mast cell activation | 1.13E-16 | 1.92E-12 | 1 | 4 | 8 | 12525 |
| GO:0021799 | BP | cerebral cortex radially oriented cell migration | 3.86E-15 | 6.58E-11 | 1 | 4 | 9 | 12524 |
| GO:0043299 | BP | leukocyte degranulation | 7.00E-14 | 1.19E-09 | 1 | 4 | 10 | 12523 |
| GO:0045686 | BP | negative regulation of glial cell differentiation | 7.86E-13 | 1.34E-08 | 1 | 4 | 11 | 12522 |
| GO:0006833 | BP | water transport | 6.10E-12 | 1.04E-07 | 1 | 4 | 12 | 12521 |
| GO:0014014 | BP | negative regulation of gliogenesis | 6.10E-12 | 1.04E-07 | 1 | 4 | 12 | 12521 |
| GO:0015669 | BP | gas transport | 6.10E-12 | 1.04E-07 | 1 | 4 | 12 | 12521 |
| GO:0048532 | BP | anatomical structure arrangement | 6.10E-12 | 1.04E-07 | 1 | 4 | 12 | 12521 |
| GO:0035082 | BP | axoneme assembly | 3.54E-11 | 6.04E-07 | 1 | 4 | 13 | 12520 |
| GO:0042044 | BP | fluid transport | 1.63E-10 | 2.78E-06 | 1 | 4 | 14 | 12519 |
| GO:0048710 | BP | regulation of astrocyte differentiation | 1.63E-10 | 2.78E-06 | 1 | 4 | 14 | 12519 |
| GO:0048593 | BP | camera-type eye morphogenesis | 6.22E-10 | 1.06E-05 | 1 | 4 | 15 | 12518 |
| GO:0009925 | CC | basal plasma membrane | 5.81E-09 | 9.90E-05 | 1 | 4 | 17 | 12516 |
| GO:0022412 | BP | cellular process involved in reproduction in multicellular organism | 5.81E-09 | 9.90E-05 | 1 | 4 | 17 | 12516 |
| GO:0032309 | BP | icosanoid secretion | 1.49E-08 | 0.0002541 | 1 | 4 | 18 | 12515 |
| GO:0050482 | BP | arachidonic acid secretion | 1.49E-08 | 0.0002541 | 1 | 4 | 18 | 12515 |
| GO:0071715 | BP | icosanoid transport | 1.49E-08 | 0.0002541 | 1 | 4 | 18 | 12515 |
| GO:0004623 | MF | phospholipase A2 activity | 7.54E-08 | 0.0012843 | 1 | 4 | 20 | 12513 |
| GO:0021795 | BP | cerebral cortex cell migration | 7.54E-08 | 0.0012843 | 1 | 4 | 20 | 12513 |
| GO:0002275 | BP | myeloid cell activation involved in immune response | 1.52E-07 | 0.0025893 | 1 | 4 | 21 | 12512 |
| GO:0016358 | BP | dendrite development | 1.52E-07 | 0.0025893 | 1 | 4 | 21 | 12512 |
| GO:0045055 | BP | regulated exocytosis | 1.52E-07 | 0.0025893 | 1 | 4 | 21 | 12512 |
| GO:0046470 | BP | phosphatidylcholine metabolic process | 1.52E-07 | 0.0025893 | 1 | 4 | 21 | 12512 |
| GO:0045685 | BP | regulation of glial cell differentiation | 8.94E-07 | 0.0152287 | 1 | 4 | 24 | 12509 |

| | | | | | | | | |
|---|---|---|---|---|---|---|---|---|
| GO:0046717 | BP | acid secretion | 1.48E-06 | 0.0251382 | 1 | 4 | 25 | 12508 |

**Supplementary Table 18 |** DEGs between five RNA-seq from the ruminal and esophageal epithelium cells of four sheep 60 days embryos.

[see the excel file]

**Supplementary Table 19 |** GO enrichment analysis of 285 highly expressed genes in the rumen compared to the esophagus. BP denotes Biological Process, MF denotes Molecular Function, and CC denotes Cellular Component.

| ID | Type | Description | *p* value | adjusted *p* value | list1InGO | list1OutGO | list2InGO | list2OutGO |
|---|---|---|---|---|---|---|---|---|
| GO:0061436 | BP | establishment of skin barrier | 1.74E-31 | 2.97E-27 | 8 | 193 | 16 | 12517 |
| GO:0033561 | BP | regulation of water loss via skin | 3.62E-30 | 6.16E-26 | 8 | 193 | 17 | 12516 |
| GO:0050891 | BP | multicellular organismal water homeostasis | 8.48E-22 | 1.44E-17 | 8 | 193 | 26 | 12507 |
| GO:0030104 | BP | water homeostasis | 3.92E-21 | 6.67E-17 | 8 | 193 | 27 | 12506 |
| GO:0004867 | MF | serine-type endopeptidase inhibitor activity | 2.81E-17 | 4.78E-13 | 10 | 191 | 55 | 12478 |
| GO:0045682 | BP | regulation of epidermis development | 5.89E-17 | 1.00E-12 | 9 | 192 | 45 | 12488 |
| GO:0044092 | BP | negative regulation of molecular function | 1.27E-16 | 2.16E-12 | 36 | 165 | 603 | 11930 |
| GO:0009913 | BP | epidermal cell differentiation | 4.12E-16 | 7.02E-12 | 8 | 193 | 37 | 12496 |
| GO:0016327 | CC | apicolateral plasma membrane | 1.49E-15 | 2.53E-11 | 5 | 196 | 13 | 12520 |
| GO:0030216 | BP | keratinocyte differentiation | 5.52E-15 | 9.41E-11 | 7 | 194 | 30 | 12503 |
| GO:0043086 | BP | negative regulation of catalytic activity | 3.66E-14 | 6.23E-10 | 28 | 173 | 442 | 12091 |
| GO:0045683 | BP | negative regulation of epidermis development | 2.21E-13 | 3.77E-09 | 4 | 197 | 9 | 12524 |
| GO:0000982 | MF | transcription factor activity, RNA polymerase II core promoter proximal region sequence-specific binding | 3.58E-13 | 6.09E-09 | 20 | 181 | 262 | 12271 |
| GO:0010838 | BP | positive regulation of keratinocyte proliferation | 4.23E-13 | 7.20E-09 | 3 | 198 | 4 | 12529 |
| GO:0035976 | CC | transcription factor AP-1 complex | 4.23E-13 | 7.20E-09 | 3 | 198 | 4 | 12529 |
| GO:0045617 | BP | negative regulation of keratinocyte differentiation | 4.23E-13 | 7.20E-09 | 3 | 198 | 4 | 12529 |
| GO:0030855 | BP | epithelial cell differentiation | 2.94E-12 | 5.00E-08 | 13 | 188 | 128 | 12405 |
| GO:0052548 | BP | regulation of endopeptidase activity | 3.31E-12 | 5.63E-08 | 17 | 184 | 209 | 12324 |
| GO:0010837 | BP | regulation of keratinocyte proliferation | 4.30E-12 | 7.32E-08 | 5 | 196 | 18 | 12515 |

| | | | | | | | | |
|---|---|---|---|---|---|---|---|---|
| GO:0005882 | CC | intermediate filament | 7.03E-12 | 1.20E-07 | 10 | 191 | 80 | 12453 |
| GO:0045604 | BP | regulation of epidermal cell differentiation | 8.01E-12 | 1.36E-07 | 6 | 195 | 28 | 12505 |
| GO:0001228 | MF | transcriptional activator activity, RNA polymerase II transcription regulatory region sequence-specific binding | 1.07E-11 | 1.82E-07 | 18 | 183 | 239 | 12294 |
| GO:0000981 | MF | RNA polymerase II transcription factor activity, sequence-specific DNA binding | 1.14E-11 | 1.95E-07 | 27 | 174 | 475 | 12058 |
| GO:0010951 | BP | negative regulation of endopeptidase activity | 1.24E-11 | 2.11E-07 | 13 | 188 | 134 | 12399 |
| GO:0016338 | BP | calcium-independent cell-cell adhesion via plasma membrane cell-adhesion molecules | 1.64E-11 | 2.79E-07 | 3 | 198 | 5 | 12528 |
| GO:0035914 | BP | skeletal muscle cell differentiation | 2.69E-11 | 4.59E-07 | 7 | 194 | 41 | 12492 |
| GO:0050878 | BP | regulation of body fluid levels | 5.12E-11 | 8.73E-07 | 11 | 190 | 103 | 12430 |
| GO:0052547 | BP | regulation of peptidase activity | 6.22E-11 | 1.06E-06 | 17 | 184 | 228 | 12305 |
| GO:0004866 | MF | endopeptidase inhibitor activity | 6.71E-11 | 1.14E-06 | 11 | 190 | 104 | 12429 |
| GO:0001533 | CC | cornified envelope | 7.14E-11 | 1.22E-06 | 4 | 197 | 12 | 12521 |
| GO:0010466 | BP | negative regulation of peptidase activity | 1.30E-10 | 2.22E-06 | 13 | 188 | 145 | 12388 |
| GO:0043565 | MF | sequence-specific DNA binding | 1.40E-10 | 2.39E-06 | 33 | 168 | 697 | 11836 |
| GO:0061135 | MF | endopeptidase regulator activity | 1.46E-10 | 2.49E-06 | 11 | 190 | 107 | 12426 |
| GO:0045605 | BP | negative regulation of epidermal cell differentiation | 2.82E-10 | 4.81E-06 | 3 | 198 | 6 | 12527 |
| GO:0098641 | MF | cadherin binding involved in cell-cell adhesion | 3.12E-10 | 5.32E-06 | 4 | 197 | 13 | 12520 |
| GO:0001071 | MF | nucleic acid binding transcription factor activity | 3.71E-10 | 6.32E-06 | 33 | 168 | 714 | 11819 |
| GO:0003700 | MF | transcription factor activity, sequence-specific DNA binding | 3.71E-10 | 6.32E-06 | 33 | 168 | 714 | 11819 |
| GO:0016328 | CC | lateral plasma membrane | 6.18E-10 | 1.05E-05 | 6 | 195 | 34 | 12499 |
| GO:0030414 | MF | peptidase inhibitor activity | 6.20E-10 | 1.06E-05 | 11 | 190 | 113 | 12420 |
| GO:0004857 | MF | enzyme inhibitor activity | 7.05E-10 | 1.20E-05 | 16 | 185 | 222 | 12311 |
| GO:0005913 | CC | cell-cell adherens junction | 7.78E-10 | 1.32E-05 | 8 | 193 | 62 | 12471 |
| GO:0006366 | BP | transcription from RNA polymerase II promoter | 1.48E-09 | 2.51E-05 | 20 | 181 | 332 | 12201 |
| GO:0051146 | BP | striated muscle cell differentiation | 3.17E-09 | 5.39E-05 | 8 | 193 | 66 | 12467 |

| | | | | | | | | |
|---|---|---|---|---|---|---|---|---|
| GO:0032502 | BP | developmental process | 3.24E-09 | 5.53E-05 | 78 | 123 | 2664 | 9869 |
| GO:0048729 | BP | tissue morphogenesis | 3.45E-09 | 5.88E-05 | 15 | 186 | 210 | 12323 |
| GO:0018149 | BP | peptide cross-linking | 3.53E-09 | 6.01E-05 | 5 | 196 | 25 | 12508 |
| GO:0098609 | BP | cell-cell adhesion | 3.76E-09 | 6.40E-05 | 4 | 197 | 15 | 12518 |
| GO:0065009 | BP | regulation of molecular function | 4.38E-09 | 7.46E-05 | 55 | 146 | 1627 | 10906 |
| GO:0009605 | BP | response to external stimulus | 5.11E-09 | 8.71E-05 | 27 | 174 | 560 | 11973 |
| GO:0001712 | BP | ectodermal cell fate commitment | 8.17E-09 | 0.000139162 | 2 | 199 | 2 | 12531 |
| GO:0010482 | BP | regulation of epidermal cell division | 8.17E-09 | 0.000139162 | 2 | 199 | 2 | 12531 |
| GO:0050790 | BP | regulation of catalytic activity | 8.49E-09 | 0.000144537 | 46 | 155 | 1273 | 11260 |
| GO:0044212 | MF | transcription regulatory region DNA binding | 9.27E-09 | 0.000157904 | 27 | 174 | 570 | 11963 |
| GO:0000975 | MF | regulatory region DNA binding | 9.83E-09 | 0.00016738 | 27 | 174 | 571 | 11962 |
| GO:0001067 | MF | regulatory region nucleic acid binding | 9.83E-09 | 0.00016738 | 27 | 174 | 571 | 11962 |
| GO:0044421 | CC | extracellular region part | 9.87E-09 | 0.000168041 | 69 | 132 | 2287 | 10246 |
| GO:0045104 | BP | intermediate filament cytoskeleton organization | 1.40E-08 | 0.000238943 | 5 | 196 | 27 | 12506 |
| GO:0045216 | BP | cell-cell junction organization | 1.50E-08 | 0.000254898 | 8 | 193 | 71 | 12462 |
| GO:0031424 | BP | keratinization | 1.80E-08 | 0.000306926 | 3 | 198 | 8 | 12525 |
| GO:0061134 | MF | peptidase regulator activity | 2.25E-08 | 0.000383206 | 11 | 190 | 131 | 12402 |
| GO:0045103 | BP | intermediate filament-based process | 2.63E-08 | 0.000447619 | 5 | 196 | 28 | 12505 |
| GO:0070830 | BP | bicellular tight junction assembly | 2.83E-08 | 0.000481195 | 4 | 197 | 17 | 12516 |
| GO:0045095 | CC | keratin filament | 4.75E-08 | 0.00080827 | 5 | 196 | 29 | 12504 |
| GO:0051346 | BP | negative regulation of hydrolase activity | 5.72E-08 | 0.00097418 | 15 | 186 | 234 | 12299 |
| GO:0002934 | BP | desmosome organization | 8.60E-08 | 0.001465649 | 3 | 198 | 9 | 12524 |
| GO:0008544 | BP | epidermis development | 9.70E-08 | 0.001652427 | 8 | 193 | 78 | 12455 |
| GO:0009888 | BP | tissue development | 1.52E-07 | 0.00258743 | 18 | 183 | 329 | 12204 |
| GO:0042127 | BP | regulation of cell proliferation | 1.75E-07 | 0.002986486 | 33 | 168 | 846 | 11687 |

| | | | | | | | | |
|---|---|---|---|---|---|---|---|---|
| GO:0006351 | BP | transcription, DNA-templated | 2.29E-07 | 0.003900569 | 30 | 171 | 740 | 11793 |
| GO:0009653 | BP | anatomical structure morphogenesis | 3.12E-07 | 0.005305914 | 35 | 166 | 939 | 11594 |
| GO:0048730 | BP | epidermis morphogenesis | 3.24E-07 | 0.005516929 | 3 | 198 | 10 | 12523 |
| GO:0061029 | BP | eyelid development in camera-type eye | 3.24E-07 | 0.005516929 | 3 | 198 | 10 | 12523 |
| GO:0031077 | BP | post-embryonic camera-type eye development | 3.40E-07 | 0.005789935 | 2 | 199 | 3 | 12530 |
| GO:0001078 | MF | transcriptional repressor activity, RNA polymerase II core promoter proximal region sequence-specific binding | 3.83E-07 | 0.006526352 | 8 | 193 | 84 | 12449 |
| GO:0005911 | CC | cell-cell junction | 4.89E-07 | 0.008331672 | 16 | 185 | 285 | 12248 |
| GO:0001077 | MF | transcriptional activator activity, RNA polymerase II core promoter proximal region sequence-specific binding | 4.91E-07 | 0.008354825 | 12 | 189 | 176 | 12357 |
| GO:0048523 | BP | negative regulation of cellular process | 5.50E-07 | 0.009364706 | 65 | 136 | 2289 | 10244 |
| GO:0045616 | BP | regulation of keratinocyte differentiation | 6.11E-07 | 0.010400482 | 4 | 197 | 21 | 12512 |
| GO:0048646 | BP | anatomical structure formation involved in morphogenesis | 6.40E-07 | 0.01089856 | 35 | 166 | 960 | 11573 |
| GO:0042692 | BP | muscle cell differentiation | 8.69E-07 | 0.014806701 | 8 | 193 | 88 | 12445 |
| GO:0032774 | BP | RNA biosynthetic process | 9.28E-07 | 0.015801132 | 30 | 171 | 775 | 11758 |
| GO:0051336 | BP | regulation of hydrolase activity | 1.03E-06 | 0.017480497 | 28 | 173 | 702 | 11831 |
| GO:0005198 | MF | structural molecule activity | 1.11E-06 | 0.01898234 | 18 | 183 | 357 | 12176 |
| GO:0045684 | BP | positive regulation of epidermis development | 1.14E-06 | 0.019367483 | 4 | 197 | 22 | 12511 |
| GO:0030154 | BP | cell differentiation | 1.24E-06 | 0.021111449 | 37 | 164 | 1062 | 11471 |
| GO:0016337 | BP | single organismal cell-cell adhesion | 1.28E-06 | 0.021730588 | 14 | 187 | 239 | 12294 |
| GO:0002009 | BP | morphogenesis of an epithelium | 1.28E-06 | 0.021859368 | 11 | 190 | 159 | 12374 |
| GO:0034330 | BP | cell junction organization | 1.85E-06 | 0.031433647 | 8 | 193 | 92 | 12441 |
| GO:0097110 | MF | scaffold protein binding | 2.02E-06 | 0.034455658 | 4 | 197 | 23 | 12510 |
| GO:1900745 | BP | positive regulation of p38MAPK cascade | 2.72E-06 | 0.046275604 | 3 | 198 | 12 | 12521 |

603

**Supplementary Table 20 |** GO enrichment analysis of 1840 highly expressed genes in the esophagus compared to the rumen. BP denotes Biological Process, MF denotes Molecular Function, and CC denotes Cellular Component.

| ID | Type | Description | *p* value | adjusted *p* value | list1InGO | list1OutGO | list2InGO | list2OutGO |
|---|---|---|---|---|---|---|---|---|
| GO:0031012 | CC | extracellular matrix | 8.52E-48 | 1.45E-43 | 123 | 1181 | 283 | 12250 |
| GO:0005578 | CC | proteinaceous extracellular matrix | 2.35E-35 | 4.00E-31 | 79 | 1225 | 162 | 12371 |
| GO:0005615 | CC | extracellular space | 5.52E-35 | 9.40E-31 | 198 | 1106 | 757 | 11776 |
| GO:0044420 | CC | extracellular matrix component | 2.01E-31 | 3.42E-27 | 60 | 1244 | 107 | 12426 |
| GO:0044449 | CC | contractile fiber part | 2.77E-31 | 4.72E-27 | 59 | 1245 | 104 | 12429 |
| GO:0051239 | BP | regulation of multicellular organismal process | 5.30E-31 | 9.02E-27 | 319 | 985 | 1601 | 10932 |
| GO:0048646 | BP | anatomical structure formation involved in morphogenesis | 1.22E-28 | 2.08E-24 | 218 | 1086 | 960 | 11573 |
| GO:2000026 | BP | regulation of multicellular organismal development | 1.17E-27 | 1.99E-23 | 227 | 1077 | 1033 | 11500 |
| GO:0032502 | BP | developmental process | 1.57E-27 | 2.67E-23 | 450 | 854 | 2664 | 9869 |
| GO:0050793 | BP | regulation of developmental process | 1.30E-26 | 2.21E-22 | 268 | 1036 | 1327 | 11206 |
| GO:0051270 | BP | regulation of cellular component movement | 3.64E-25 | 6.20E-21 | 125 | 1179 | 445 | 12088 |
| GO:0022603 | BP | regulation of anatomical structure morphogenesis | 5.64E-25 | 9.61E-21 | 142 | 1162 | 543 | 11990 |
| GO:0044421 | CC | extracellular region part | 1.34E-24 | 2.27E-20 | 392 | 912 | 2287 | 10246 |
| GO:0048856 | BP | anatomical structure development | 2.17E-24 | 3.70E-20 | 291 | 1013 | 1534 | 10999 |
| GO:0040012 | BP | regulation of locomotion | 8.41E-24 | 1.43E-19 | 127 | 1177 | 470 | 12063 |
| GO:0030334 | BP | regulation of cell migration | 3.19E-23 | 5.44E-19 | 115 | 1189 | 409 | 12124 |
| GO:2000145 | BP | regulation of cell motility | 4.43E-23 | 7.55E-19 | 118 | 1186 | 427 | 12106 |
| GO:0005576 | CC | extracellular region | 5.43E-23 | 9.25E-19 | 150 | 1154 | 614 | 11919 |
| GO:0003012 | BP | muscle system process | 5.46E-22 | 9.30E-18 | 55 | 1249 | 125 | 12408 |
| GO:0030198 | BP | extracellular matrix organization | 1.78E-20 | 3.03E-16 | 53 | 1251 | 124 | 12409 |

| | | | | | | | | |
|---|---|---|---|---|---|---|---|---|
| GO:0043062 | BP | extracellular structure organization | 1.78E-20 | 3.03E-16 | 53 | 1251 | 124 | 12409 |
| GO:0032879 | BP | regulation of localization | 6.12E-20 | 1.04E-15 | 252 | 1052 | 1350 | 11183 |
| GO:0009653 | BP | anatomical structure morphogenesis | 8.15E-20 | 1.39E-15 | 193 | 1111 | 939 | 11594 |
| GO:0022610 | BP | biological adhesion | 8.25E-20 | 1.41E-15 | 108 | 1196 | 405 | 12128 |
| GO:0003779 | MF | actin binding | 2.20E-19 | 3.75E-15 | 74 | 1230 | 227 | 12306 |
| GO:0007155 | BP | cell adhesion | 2.22E-19 | 3.78E-15 | 107 | 1197 | 404 | 12129 |
| GO:0006936 | BP | muscle contraction | 7.43E-19 | 1.27E-14 | 45 | 1259 | 99 | 12434 |
| GO:0044428 | CC | nuclear part | 8.19E-19 | 1.40E-14 | 132 | 1172 | 2551 | 9982 |
| GO:0001525 | BP | angiogenesis | 1.62E-18 | 2.75E-14 | 49 | 1255 | 117 | 12416 |
| GO:0048869 | BP | cellular developmental process | 8.88E-18 | 1.51E-13 | 283 | 1021 | 1633 | 10900 |
| GO:0048729 | BP | tissue morphogenesis | 1.08E-17 | 1.84E-13 | 68 | 1236 | 210 | 12323 |
| GO:0030018 | CC | Z disc | 2.84E-17 | 4.84E-13 | 33 | 1271 | 60 | 12473 |
| GO:1901342 | BP | regulation of vasculature development | 4.30E-17 | 7.33E-13 | 54 | 1250 | 147 | 12386 |
| GO:0048513 | BP | animal organ development | 4.61E-17 | 7.86E-13 | 138 | 1166 | 623 | 11910 |
| GO:0040011 | BP | locomotion | 2.13E-16 | 3.63E-12 | 132 | 1172 | 595 | 11938 |
| GO:0045765 | BP | regulation of angiogenesis | 2.51E-16 | 4.28E-12 | 51 | 1253 | 138 | 12395 |
| GO:0005581 | CC | collagen trimer | 2.63E-16 | 4.48E-12 | 28 | 1276 | 46 | 12487 |
| GO:0008092 | MF | cytoskeletal protein binding | 4.73E-16 | 8.05E-12 | 119 | 1185 | 518 | 12015 |
| GO:0060284 | BP | regulation of cell development | 1.54E-15 | 2.62E-11 | 117 | 1187 | 513 | 12020 |
| GO:0009888 | BP | tissue development | 2.49E-15 | 4.25E-11 | 86 | 1218 | 329 | 12204 |
| GO:0042383 | CC | sarcolemma | 3.41E-15 | 5.81E-11 | 29 | 1275 | 53 | 12480 |
| GO:0045595 | BP | regulation of cell differentiation | 3.77E-15 | 6.43E-11 | 179 | 1125 | 935 | 11598 |
| GO:0060537 | BP | muscle tissue development | 4.46E-15 | 7.60E-11 | 30 | 1274 | 57 | 12476 |
| GO:0044446 | CC | intracellular organelle part | 6.23E-15 | 1.06E-10 | 333 | 971 | 4566 | 7967 |
| GO:0005604 | CC | basement membrane | 6.59E-15 | 1.12E-10 | 29 | 1275 | 54 | 12479 |

| | | | | | | | | |
|---|---|---|---|---|---|---|---|---|
| GO:0043292 | CC | contractile fiber | 7.55E-15 | 1.29E-10 | 22 | 1282 | 31 | 12502 |
| GO:0003676 | MF | nucleic acid binding | 8.19E-15 | 1.39E-10 | 138 | 1166 | 2433 | 10100 |
| GO:0005509 | MF | calcium ion binding | 9.38E-15 | 1.60E-10 | 100 | 1204 | 419 | 12114 |
| GO:0040017 | BP | positive regulation of locomotion | 1.29E-14 | 2.20E-10 | 70 | 1234 | 247 | 12286 |
| GO:0005539 | MF | glycosaminoglycan binding | 1.33E-14 | 2.26E-10 | 40 | 1264 | 99 | 12434 |
| GO:0014706 | BP | striated muscle tissue development | 1.35E-14 | 2.30E-10 | 27 | 1277 | 48 | 12485 |
| GO:0097367 | MF | carbohydrate derivative binding | 1.48E-14 | 2.52E-10 | 42 | 1262 | 108 | 12425 |
| GO:0051094 | BP | positive regulation of developmental process | 1.62E-14 | 2.76E-10 | 130 | 1174 | 613 | 11920 |
| GO:2000147 | BP | positive regulation of cell motility | 1.73E-14 | 2.95E-10 | 67 | 1237 | 232 | 12301 |
| GO:0030335 | BP | positive regulation of cell migration | 1.90E-14 | 3.23E-10 | 66 | 1238 | 227 | 12306 |
| GO:0051272 | BP | positive regulation of cellular component movement | 2.21E-14 | 3.77E-10 | 67 | 1237 | 233 | 12300 |
| GO:0016525 | BP | negative regulation of angiogenesis | 1.85E-13 | 3.16E-09 | 27 | 1277 | 52 | 12481 |
| GO:0055001 | BP | muscle cell development | 2.20E-13 | 3.75E-09 | 28 | 1276 | 56 | 12477 |
| GO:0030029 | BP | actin filament-based process | 5.64E-13 | 9.61E-09 | 70 | 1234 | 264 | 12269 |
| GO:0061061 | BP | muscle structure development | 6.38E-13 | 1.09E-08 | 21 | 1283 | 33 | 12500 |
| GO:0003723 | MF | RNA binding | 7.79E-13 | 1.33E-08 | 32 | 1272 | 999 | 11534 |
| GO:0016477 | BP | cell migration | 1.01E-12 | 1.72E-08 | 95 | 1209 | 417 | 12116 |
| GO:0007517 | BP | muscle organ development | 1.72E-12 | 2.93E-08 | 20 | 1284 | 31 | 12502 |
| GO:0048519 | BP | negative regulation of biological process | 2.07E-12 | 3.53E-08 | 367 | 937 | 2485 | 10048 |
| GO:0030036 | BP | actin cytoskeleton organization | 3.00E-12 | 5.10E-08 | 67 | 1237 | 255 | 12278 |
| GO:0030016 | CC | myofibril | 3.48E-12 | 5.93E-08 | 15 | 1289 | 17 | 12516 |
| GO:0005518 | MF | collagen binding | 4.58E-12 | 7.80E-08 | 22 | 1282 | 39 | 12494 |
| GO:0030154 | BP | cell differentiation | 5.35E-12 | 9.12E-08 | 186 | 1118 | 1062 | 11471 |
| GO:0007275 | BP | multicellular organism development | 5.65E-12 | 9.62E-08 | 64 | 1240 | 241 | 12292 |
| GO:0006941 | BP | striated muscle contraction | 7.62E-12 | 1.30E-07 | 24 | 1280 | 47 | 12486 |

| | | | | | | | | |
|---|---|---|---|---|---|---|---|---|
| GO:0044422 | CC | organelle part | 8.39E-12 | 1.43E-07 | 358 | 946 | 4643 | 7890 |
| GO:0005515 | MF | protein binding | 1.08E-11 | 1.85E-07 | 523 | 781 | 3868 | 8665 |
| GO:0050907 | BP | detection of chemical stimulus involved in sensory perception | 1.30E-11 | 2.21E-07 | 0 | 1304 | 437 | 12096 |
| GO:0006807 | BP | nitrogen compound metabolic process | 1.65E-11 | 2.81E-07 | 109 | 1195 | 1922 | 10611 |
| GO:0048870 | BP | cell motility | 1.67E-11 | 2.85E-07 | 99 | 1205 | 462 | 12071 |
| GO:0090304 | BP | nucleic acid metabolic process | 2.03E-11 | 3.45E-07 | 81 | 1223 | 1578 | 10955 |
| GO:0046483 | BP | heterocycle metabolic process | 2.27E-11 | 3.87E-07 | 105 | 1199 | 1867 | 10666 |
| GO:0007519 | BP | skeletal muscle tissue development | 2.29E-11 | 3.90E-07 | 19 | 1285 | 31 | 12502 |
| GO:0048518 | BP | positive regulation of biological process | 2.55E-11 | 4.35E-07 | 422 | 882 | 3001 | 9532 |
| GO:0034641 | BP | cellular nitrogen compound metabolic process | 2.80E-11 | 4.77E-07 | 108 | 1196 | 1898 | 10635 |
| GO:0005102 | MF | receptor binding | 3.08E-11 | 5.25E-07 | 148 | 1156 | 804 | 11729 |
| GO:0048583 | BP | regulation of response to stimulus | 3.21E-11 | 5.46E-07 | 292 | 1012 | 1915 | 10618 |
| GO:0004984 | MF | olfactory receptor activity | 3.80E-11 | 6.47E-07 | 0 | 1304 | 418 | 12115 |
| GO:0050911 | BP | detection of chemical stimulus involved in sensory perception of smell | 3.80E-11 | 6.47E-07 | 0 | 1304 | 418 | 12115 |
| GO:0050906 | BP | detection of stimulus involved in sensory perception | 4.27E-11 | 7.26E-07 | 3 | 1301 | 472 | 12061 |
| GO:1901360 | BP | organic cyclic compound metabolic process | 6.57E-11 | 1.12E-06 | 115 | 1189 | 1961 | 10572 |
| GO:0006935 | BP | chemotaxis | 7.24E-11 | 1.23E-06 | 63 | 1241 | 248 | 12285 |
| GO:0042330 | BP | taxis | 7.24E-11 | 1.23E-06 | 63 | 1241 | 248 | 12285 |
| GO:0010594 | BP | regulation of endothelial cell migration | 7.45E-11 | 1.27E-06 | 30 | 1274 | 76 | 12457 |
| GO:0006725 | BP | cellular aromatic compound metabolic process | 7.96E-11 | 1.36E-06 | 107 | 1197 | 1862 | 10671 |
| GO:0097159 | MF | organic cyclic compound binding | 8.49E-11 | 1.45E-06 | 262 | 1042 | 3584 | 8949 |
| GO:0051271 | BP | negative regulation of cellular component movement | 9.15E-11 | 1.56E-06 | 44 | 1260 | 144 | 12389 |
| GO:1901363 | MF | heterocyclic compound binding | 9.14E-11 | 1.56E-06 | 262 | 1042 | 3582 | 8951 |
| GO:0006139 | BP | nucleobase-containing compound metabolic process | 9.27E-11 | 1.58E-06 | 101 | 1203 | 1787 | 10746 |
| GO:0008201 | MF | heparin binding | 9.88E-11 | 1.68E-06 | 28 | 1276 | 68 | 12465 |

| | | | | | | | | |
|---|---|---|---|---|---|---|---|---|
| GO:0030336 | BP | negative regulation of cell migration | 9.86E-11 | 1.68E-06 | 42 | 1262 | 134 | 12399 |
| GO:0023051 | BP | regulation of signaling | 1.03E-10 | 1.75E-06 | 261 | 1043 | 1684 | 10849 |
| GO:0009966 | BP | regulation of signal transduction | 1.11E-10 | 1.89E-06 | 239 | 1065 | 1510 | 11023 |
| GO:0051093 | BP | negative regulation of developmental process | 1.13E-10 | 1.92E-06 | 98 | 1206 | 470 | 12063 |
| GO:0005634 | CC | nucleus | 1.13E-10 | 1.93E-06 | 224 | 1080 | 3170 | 9363 |
| GO:0007608 | BP | sensory perception of smell | 1.16E-10 | 1.97E-06 | 2 | 1302 | 436 | 12097 |
| GO:2000146 | BP | negative regulation of cell motility | 1.26E-10 | 2.14E-06 | 43 | 1261 | 140 | 12393 |
| GO:0007606 | BP | sensory perception of chemical stimulus | 1.46E-10 | 2.49E-06 | 4 | 1300 | 468 | 12065 |
| GO:0009593 | BP | detection of chemical stimulus | 1.65E-10 | 2.81E-06 | 3 | 1301 | 448 | 12085 |
| GO:0060415 | BP | muscle tissue morphogenesis | 1.71E-10 | 2.91E-06 | 21 | 1283 | 41 | 12492 |
| GO:0040013 | BP | negative regulation of locomotion | 1.75E-10 | 2.98E-06 | 46 | 1258 | 157 | 12376 |
| GO:0010646 | BP | regulation of cell communication | 1.77E-10 | 3.01E-06 | 262 | 1042 | 1701 | 10832 |
| GO:0032970 | BP | regulation of actin filament-based process | 1.90E-10 | 3.24E-06 | 52 | 1252 | 190 | 12343 |
| GO:0050840 | MF | extracellular matrix binding | 2.00E-10 | 3.40E-06 | 19 | 1285 | 34 | 12499 |
| GO:0048731 | BP | system development | 2.32E-10 | 3.96E-06 | 74 | 1230 | 321 | 12212 |
| GO:0010810 | BP | regulation of cell-substrate adhesion | 2.43E-10 | 4.13E-06 | 37 | 1267 | 112 | 12421 |
| GO:0002009 | BP | morphogenesis of an epithelium | 2.87E-10 | 4.89E-06 | 46 | 1258 | 159 | 12374 |
| GO:0044459 | CC | plasma membrane part | 3.02E-10 | 5.15E-06 | 200 | 1104 | 1220 | 11313 |
| GO:0005901 | CC | caveola | 3.88E-10 | 6.61E-06 | 19 | 1285 | 35 | 12498 |
| GO:0030155 | BP | regulation of cell adhesion | 5.29E-10 | 9.02E-06 | 60 | 1244 | 241 | 12292 |
| GO:0048523 | BP | negative regulation of cellular process | 5.36E-10 | 9.13E-06 | 331 | 973 | 2289 | 10244 |
| GO:0048641 | BP | regulation of skeletal muscle tissue development | 6.28E-10 | 1.07E-05 | 27 | 1277 | 68 | 12465 |
| GO:0048468 | BP | cell development | 6.41E-10 | 1.09E-05 | 85 | 1219 | 398 | 12135 |
| GO:0005654 | CC | nucleoplasm | 7.43E-10 | 1.27E-05 | 68 | 1236 | 1337 | 11196 |
| GO:0019838 | MF | growth factor binding | 7.91E-10 | 1.35E-05 | 28 | 1276 | 73 | 12460 |

| | | | | | | | | |
|---|---|---|---|---|---|---|---|---|
| GO:0023056 | BP | positive regulation of signaling | 8.55E-10 | 1.46E-05 | 135 | 1169 | 746 | 11787 |
| GO:0034762 | BP | regulation of transmembrane transport | 8.96E-10 | 1.53E-05 | 71 | 1233 | 311 | 12222 |
| GO:0045597 | BP | positive regulation of cell differentiation | 9.18E-10 | 1.56E-05 | 89 | 1215 | 427 | 12106 |
| GO:0030529 | CC | intracellular ribonucleoprotein complex | 1.07E-09 | 1.83E-05 | 7 | 1297 | 484 | 12049 |
| GO:0048522 | BP | positive regulation of cellular process | 1.11E-09 | 1.88E-05 | 381 | 923 | 2729 | 9804 |
| GO:0010647 | BP | positive regulation of cell communication | 1.12E-09 | 1.91E-05 | 135 | 1169 | 749 | 11784 |
| GO:0022604 | BP | regulation of cell morphogenesis | 1.19E-09 | 2.03E-05 | 67 | 1237 | 288 | 12245 |
| GO:0043269 | BP | regulation of ion transport | 1.22E-09 | 2.08E-05 | 66 | 1238 | 282 | 12251 |
| GO:0031032 | BP | actomyosin structure organization | 1.43E-09 | 2.44E-05 | 24 | 1280 | 57 | 12476 |
| GO:0006396 | BP | RNA processing | 1.47E-09 | 2.50E-05 | 3 | 1301 | 409 | 12124 |
| GO:0043231 | CC | intracellular membrane-bounded organelle | 1.65E-09 | 2.81E-05 | 457 | 847 | 5486 | 7047 |
| GO:0044427 | CC | chromosomal part | 2.33E-09 | 3.96E-05 | 9 | 1295 | 503 | 12030 |
| GO:0045121 | CC | membrane raft | 2.48E-09 | 4.22E-05 | 37 | 1267 | 120 | 12413 |
| GO:0034765 | BP | regulation of ion transmembrane transport | 2.68E-09 | 4.57E-05 | 51 | 1253 | 197 | 12336 |
| GO:0032412 | BP | regulation of ion transmembrane transporter activity | 3.07E-09 | 5.23E-05 | 27 | 1277 | 72 | 12461 |
| GO:0000904 | BP | cell morphogenesis involved in differentiation | 3.08E-09 | 5.25E-05 | 38 | 1266 | 126 | 12407 |
| GO:0005201 | MF | extracellular matrix structural constituent | 3.08E-09 | 5.25E-05 | 17 | 1287 | 31 | 12502 |
| GO:0009967 | BP | positive regulation of signal transduction | 3.22E-09 | 5.49E-05 | 128 | 1176 | 710 | 11823 |
| GO:0044057 | BP | regulation of system process | 3.36E-09 | 5.73E-05 | 74 | 1230 | 339 | 12194 |
| GO:0007507 | BP | heart development | 3.45E-09 | 5.87E-05 | 40 | 1264 | 137 | 12396 |
| GO:0016202 | BP | regulation of striated muscle tissue development | 3.55E-09 | 6.04E-05 | 34 | 1270 | 106 | 12427 |
| GO:0022898 | BP | regulation of transmembrane transporter activity | 3.57E-09 | 6.08E-05 | 28 | 1276 | 77 | 12456 |
| GO:0048584 | BP | positive regulation of response to stimulus | 4.15E-09 | 7.07E-05 | 156 | 1148 | 920 | 11613 |
| GO:0005912 | CC | adherens junction | 4.81E-09 | 8.19E-05 | 67 | 1237 | 297 | 12236 |
| GO:0005924 | CC | cell-substrate adherens junction | 6.26E-09 | 0.000107 | 57 | 1247 | 237 | 12296 |

| | | | | | | | | |
|---|---|---|---|---|---|---|---|---|
| GO:0032956 | BP | regulation of actin cytoskeleton organization | 6.26E-09 | 0.000107 | 48 | 1256 | 184 | 12349 |
| GO:0048634 | BP | regulation of muscle organ development | 6.31E-09 | 0.000107 | 34 | 1270 | 108 | 12425 |
| GO:0051147 | BP | regulation of muscle cell differentiation | 6.41E-09 | 0.000109 | 32 | 1272 | 98 | 12435 |
| GO:0030055 | CC | cell-substrate junction | 8.79E-09 | 0.00015 | 57 | 1247 | 239 | 12294 |
| GO:0050896 | BP | response to stimulus | 9.40E-09 | 0.00016 | 544 | 760 | 4228 | 8305 |
| GO:0051240 | BP | positive regulation of multicellular organismal process | 9.93E-09 | 0.000169 | 81 | 1223 | 393 | 12140 |
| GO:0051606 | BP | detection of stimulus | 9.93E-09 | 0.000169 | 12 | 1292 | 524 | 12009 |
| GO:0044237 | BP | cellular metabolic process | 1.05E-08 | 0.00018 | 357 | 947 | 4429 | 8104 |
| GO:0070887 | BP | cellular response to chemical stimulus | 1.09E-08 | 0.000185 | 166 | 1138 | 1009 | 11524 |
| GO:0051015 | MF | actin filament binding | 1.09E-08 | 0.000186 | 29 | 1275 | 85 | 12448 |
| GO:0071363 | BP | cellular response to growth factor stimulus | 1.15E-08 | 0.000195 | 40 | 1264 | 142 | 12391 |
| GO:0042127 | BP | regulation of cell proliferation | 1.45E-08 | 0.000247 | 144 | 1160 | 846 | 11687 |
| GO:0070848 | BP | response to growth factor | 1.61E-08 | 0.000274 | 41 | 1263 | 149 | 12384 |
| GO:0048742 | BP | regulation of skeletal muscle fiber development | 1.65E-08 | 0.000282 | 22 | 1282 | 54 | 12479 |
| GO:0065007 | BP | biological regulation | 1.69E-08 | 0.000288 | 838 | 466 | 7030 | 5503 |
| GO:0048747 | BP | muscle fiber development | 1.97E-08 | 0.000335 | 17 | 1287 | 34 | 12499 |
| GO:0070161 | CC | anchoring junction | 2.05E-08 | 0.000349 | 67 | 1237 | 307 | 12226 |
| GO:0043537 | BP | negative regulation of blood vessel endothelial cell migration | 2.43E-08 | 0.000414 | 12 | 1292 | 17 | 12516 |
| GO:2001014 | BP | regulation of skeletal muscle cell differentiation | 2.46E-08 | 0.000419 | 15 | 1289 | 27 | 12506 |
| GO:0051960 | BP | regulation of nervous system development | 2.68E-08 | 0.000457 | 82 | 1222 | 408 | 12125 |
| GO:0006928 | BP | movement of cell or subcellular component | 2.95E-08 | 0.000503 | 107 | 1197 | 583 | 11950 |
| GO:0007010 | BP | cytoskeleton organization | 3.18E-08 | 0.000542 | 99 | 1205 | 527 | 12006 |
| GO:0005925 | CC | focal adhesion | 3.41E-08 | 0.00058 | 55 | 1249 | 235 | 12298 |
| GO:0055002 | BP | striated muscle cell development | 3.47E-08 | 0.00059 | 17 | 1287 | 35 | 12498 |
| GO:0051153 | BP | regulation of striated muscle cell differentiation | 3.59E-08 | 0.000611 | 27 | 1277 | 79 | 12454 |

| | | | | | | | | |
|---|---|---|---|---|---|---|---|---|
| GO:0001936 | BP | regulation of endothelial cell proliferation | 3.82E-08 | 0.00065 | 22 | 1282 | 56 | 12477 |
| GO:0009986 | CC | cell surface | 4.26E-08 | 0.000726 | 64 | 1240 | 293 | 12240 |
| GO:0030278 | BP | regulation of ossification | 4.42E-08 | 0.000753 | 36 | 1268 | 126 | 12407 |
| GO:0007600 | BP | sensory perception | 4.46E-08 | 0.00076 | 23 | 1281 | 658 | 11875 |
| GO:0032409 | BP | regulation of transporter activity | 5.09E-08 | 0.000868 | 28 | 1276 | 85 | 12448 |
| GO:0031589 | BP | cell-substrate adhesion | 5.12E-08 | 0.000873 | 29 | 1275 | 90 | 12443 |
| GO:0050678 | BP | regulation of epithelial cell proliferation | 5.42E-08 | 0.000923 | 40 | 1264 | 149 | 12384 |
| GO:0030017 | CC | sarcomere | 5.78E-08 | 0.000984 | 12 | 1292 | 18 | 12515 |
| GO:0043535 | BP | regulation of blood vessel endothelial cell migration | 5.97E-08 | 0.001017 | 17 | 1287 | 36 | 12497 |
| GO:0032501 | BP | multicellular organismal process | 6.42E-08 | 0.001094 | 281 | 1023 | 1968 | 10565 |
| GO:0044707 | BP | single-multicellular organism process | 6.85E-08 | 0.001167 | 280 | 1024 | 1961 | 10572 |
| GO:0071704 | BP | organic substance metabolic process | 7.69E-08 | 0.00131 | 147 | 1157 | 2147 | 10386 |
| GO:0006974 | BP | cellular response to DNA damage stimulus | 8.00E-08 | 0.001363 | 5 | 1299 | 372 | 12161 |
| GO:0010596 | BP | negative regulation of endothelial cell migration | 8.75E-08 | 0.001491 | 15 | 1289 | 29 | 12504 |
| GO:0050789 | BP | regulation of biological process | 9.14E-08 | 0.001557 | 804 | 500 | 6752 | 5781 |
| GO:0042325 | BP | regulation of phosphorylation | 9.36E-08 | 0.001594 | 123 | 1181 | 713 | 11820 |
| GO:0055008 | BP | cardiac muscle tissue morphogenesis | 1.01E-07 | 0.001716 | 17 | 1287 | 37 | 12496 |
| GO:0010033 | BP | response to organic substance | 1.02E-07 | 0.001741 | 167 | 1137 | 1050 | 11483 |
| GO:0030054 | CC | cell junction | 1.07E-07 | 0.001828 | 110 | 1194 | 619 | 11914 |
| GO:0050767 | BP | regulation of neurogenesis | 1.28E-07 | 0.002183 | 72 | 1232 | 354 | 12179 |
| GO:0000902 | BP | cell morphogenesis | 1.30E-07 | 0.002217 | 41 | 1263 | 159 | 12374 |
| GO:0048585 | BP | negative regulation of response to stimulus | 1.36E-07 | 0.002318 | 119 | 1185 | 688 | 11845 |
| GO:0043230 | CC | extracellular organelle | 1.38E-07 | 0.002348 | 245 | 1059 | 1684 | 10849 |
| GO:0065010 | CC | extracellular membrane-bounded organelle | 1.38E-07 | 0.002348 | 245 | 1059 | 1684 | 10849 |
| GO:0009887 | BP | animal organ morphogenesis | 1.43E-07 | 0.002429 | 60 | 1244 | 276 | 12257 |

| | | | | | | | | |
|---|---|---|---|---|---|---|---|---|
| GO:0007167 | BP | enzyme linked receptor protein signaling pathway | 1.57E-07 | 0.002674 | 68 | 1236 | 329 | 12204 |
| GO:0009605 | BP | response to external stimulus | 1.86E-07 | 0.003173 | 101 | 1203 | 560 | 11973 |
| GO:0051049 | BP | regulation of transport | 1.92E-07 | 0.00327 | 152 | 1152 | 943 | 11590 |
| GO:0070062 | CC | extracellular exosome | 1.97E-07 | 0.003355 | 244 | 1060 | 1683 | 10850 |
| GO:0043229 | CC | intracellular organelle | 2.15E-07 | 0.003668 | 538 | 766 | 6121 | 6412 |
| GO:0051276 | BP | chromosome organization | 2.18E-07 | 0.003706 | 13 | 1291 | 481 | 12052 |
| GO:0001568 | BP | blood vessel development | 2.24E-07 | 0.003821 | 27 | 1277 | 85 | 12448 |
| GO:0010975 | BP | regulation of neuron projection development | 2.40E-07 | 0.004089 | 47 | 1257 | 198 | 12335 |
| GO:0030199 | BP | collagen fibril organization | 2.49E-07 | 0.004238 | 14 | 1290 | 27 | 12506 |
| GO:0045214 | BP | sarcomere organization | 2.49E-07 | 0.004238 | 14 | 1290 | 27 | 12506 |
| GO:0001570 | BP | vasculogenesis | 2.70E-07 | 0.004595 | 17 | 1287 | 39 | 12494 |
| GO:0005178 | MF | integrin binding | 2.72E-07 | 0.004625 | 20 | 1284 | 52 | 12481 |
| GO:0008307 | MF | structural constituent of muscle | 2.75E-07 | 0.004692 | 12 | 1292 | 20 | 12513 |
| GO:0060688 | BP | regulation of morphogenesis of a branching structure | 2.82E-07 | 0.004811 | 16 | 1288 | 35 | 12498 |
| GO:0090257 | BP | regulation of muscle system process | 3.04E-07 | 0.005178 | 26 | 1278 | 81 | 12452 |
| GO:0044451 | CC | nucleoplasm part | 3.07E-07 | 0.005232 | 40 | 1264 | 847 | 11686 |
| GO:0001932 | BP | regulation of protein phosphorylation | 3.15E-07 | 0.005366 | 113 | 1191 | 654 | 11879 |
| GO:0006898 | BP | receptor-mediated endocytosis | 3.27E-07 | 0.005578 | 30 | 1274 | 102 | 12431 |
| GO:2000027 | BP | regulation of organ morphogenesis | 3.27E-07 | 0.005578 | 30 | 1274 | 102 | 12431 |
| GO:0008152 | BP | metabolic process | 3.48E-07 | 0.005932 | 439 | 865 | 5136 | 7397 |
| GO:0051493 | BP | regulation of cytoskeleton organization | 3.66E-07 | 0.006237 | 58 | 1246 | 270 | 12263 |
| GO:0044238 | BP | primary metabolic process | 3.80E-07 | 0.00647 | 375 | 929 | 4494 | 8039 |
| GO:0010811 | BP | positive regulation of cell-substrate adhesion | 3.89E-07 | 0.006627 | 23 | 1281 | 67 | 12466 |
| GO:0005739 | CC | mitochondrion | 4.39E-07 | 0.007473 | 53 | 1251 | 1004 | 11529 |
| GO:0042692 | BP | muscle cell differentiation | 5.17E-07 | 0.008805 | 27 | 1277 | 88 | 12445 |

| | | | | | | | | |
|---|---|---|---|---|---|---|---|---|
| GO:0016070 | BP | RNA metabolic process | 5.21E-07 | 0.00887 | 73 | 1231 | 1244 | 11289 |
| GO:0043408 | BP | regulation of MAPK cascade | 5.40E-07 | 0.009203 | 77 | 1227 | 401 | 12132 |
| GO:0090092 | BP | regulation of transmembrane receptor protein serine/threonine kinase signaling pathway | 6.10E-07 | 0.010387 | 36 | 1268 | 138 | 12395 |
| GO:0045664 | BP | regulation of neuron differentiation | 6.13E-07 | 0.010445 | 60 | 1244 | 287 | 12246 |
| GO:0070374 | BP | positive regulation of ERK1 and ERK2 cascade | 6.14E-07 | 0.010454 | 31 | 1273 | 110 | 12423 |
| GO:0044260 | BP | cellular macromolecule metabolic process | 6.19E-07 | 0.010551 | 265 | 1039 | 3351 | 9182 |
| GO:0070372 | BP | regulation of ERK1 and ERK2 cascade | 6.62E-07 | 0.01127 | 38 | 1266 | 150 | 12383 |
| GO:0035914 | BP | skeletal muscle cell differentiation | 6.71E-07 | 0.011436 | 17 | 1287 | 41 | 12492 |
| GO:0060048 | BP | cardiac muscle contraction | 7.09E-07 | 0.012071 | 13 | 1291 | 25 | 12508 |
| GO:0003013 | BP | circulatory system process | 7.44E-07 | 0.012673 | 24 | 1280 | 74 | 12459 |
| GO:0019220 | BP | regulation of phosphate metabolic process | 8.72E-07 | 0.014859 | 127 | 1177 | 773 | 11760 |
| GO:0051174 | BP | regulation of phosphorus metabolic process | 9.35E-07 | 0.015931 | 127 | 1177 | 774 | 11759 |
| GO:0055082 | BP | cellular chemical homeostasis | 9.59E-07 | 0.01634 | 84 | 1220 | 456 | 12077 |
| GO:0044424 | CC | intracellular part | 1.02E-06 | 0.017349 | 756 | 548 | 8126 | 4407 |
| GO:0031674 | CC | I band | 1.09E-06 | 0.018605 | 9 | 1295 | 12 | 12521 |
| GO:0010648 | BP | negative regulation of cell communication | 1.33E-06 | 0.022729 | 103 | 1201 | 598 | 11935 |
| GO:0006259 | BP | DNA metabolic process | 1.45E-06 | 0.024691 | 10 | 1294 | 399 | 12134 |
| GO:0010959 | BP | regulation of metal ion transport | 1.61E-06 | 0.027367 | 36 | 1268 | 143 | 12390 |
| GO:0045785 | BP | positive regulation of cell adhesion | 1.63E-06 | 0.027749 | 32 | 1272 | 120 | 12413 |
| GO:0031982 | CC | vesicle | 1.77E-06 | 0.030083 | 283 | 1021 | 2061 | 10472 |
| GO:0007165 | BP | signal transduction | 1.83E-06 | 0.031099 | 360 | 944 | 2730 | 9803 |
| GO:0050920 | BP | regulation of chemotaxis | 1.87E-06 | 0.03181 | 27 | 1277 | 93 | 12440 |
| GO:0061138 | BP | morphogenesis of a branching epithelium | 1.87E-06 | 0.03181 | 27 | 1277 | 93 | 12440 |
| GO:0032414 | BP | positive regulation of ion transmembrane transporter activity | 2.02E-06 | 0.034468 | 12 | 1292 | 23 | 12510 |
| GO:0044699 | BP | single-organism process | 2.05E-06 | 0.034927 | 319 | 985 | 2375 | 10158 |

| | | | | | | | | |
|---|---|---|---|---|---|---|---|---|
| GO:0009968 | BP | negative regulation of signal transduction | 2.09E-06 | 0.035564 | 99 | 1205 | 574 | 11959 |
| GO:0065008 | BP | regulation of biological quality | 2.11E-06 | 0.035899 | 227 | 1077 | 1592 | 10941 |
| GO:0006873 | BP | cellular ion homeostasis | 2.15E-06 | 0.036642 | 79 | 1225 | 429 | 12104 |
| GO:0070167 | BP | regulation of biomineral tissue development | 2.16E-06 | 0.036805 | 19 | 1285 | 53 | 12480 |
| GO:0050794 | BP | regulation of cellular process | 2.17E-06 | 0.037037 | 752 | 552 | 6359 | 6174 |
| GO:0004930 | MF | G-protein coupled receptor activity | 2.22E-06 | 0.037895 | 35 | 1269 | 738 | 11795 |
| GO:2001257 | BP | regulation of cation channel activity | 2.23E-06 | 0.037942 | 13 | 1291 | 27 | 12506 |
| GO:0051051 | BP | negative regulation of transport | 2.30E-06 | 0.039213 | 48 | 1256 | 219 | 12314 |
| GO:0050801 | BP | ion homeostasis | 2.40E-06 | 0.040864 | 84 | 1220 | 466 | 12067 |
| GO:0048514 | BP | blood vessel morphogenesis | 2.43E-06 | 0.041414 | 21 | 1283 | 63 | 12470 |
| GO:0032991 | CC | macromolecular complex | 2.45E-06 | 0.041766 | 236 | 1068 | 3001 | 9532 |
| GO:0023057 | BP | negative regulation of signaling | 2.67E-06 | 0.045483 | 101 | 1203 | 592 | 11941 |
| GO:0030239 | BP | myofibril assembly | 2.69E-06 | 0.04574 | 9 | 1295 | 13 | 12520 |
| GO:0030315 | CC | T-tubule | 2.69E-06 | 0.04574 | 9 | 1295 | 13 | 12520 |
| GO:0085029 | BP | extracellular matrix assembly | 2.80E-06 | 0.047646 | 8 | 1296 | 10 | 12523 |
| GO:0014068 | BP | positive regulation of phosphatidylinositol 3-kinase signaling | 2.80E-06 | 0.047655 | 16 | 1288 | 40 | 12493 |
| GO:0034660 | BP | ncRNA metabolic process | 2.82E-06 | 0.047993 | 0 | 1304 | 218 | 12315 |
| GO:0044710 | BP | single-organism metabolic process | 2.91E-06 | 0.049622 | 230 | 1074 | 2932 | 9601 |

606

**Supplementary Table 21 |** List of 657 rumen key genes have nearby RSCNEs.

[see the excel file]

**Supplementary Table 22 |** List of rumen key genes under positive selection. The BEB sites showing the detailed selected positions in cattle.

| Ensembl ID | Official name | Description | *P*-value | BEB sites |
|---|---|---|---|---|
| ENSBTAT00000036662 | *PDE6A* | phosphodiesterase 6A [Source:HGNC Symbol;Acc:HGNC:8785] | 0.026308 | 70:E:0.612;202:R:0.966*;427:E:0.555 |
| ENSBTAT00000005088 | *HMGCS2* | 3-hydroxy-3-methylglutaryl-CoAsynthase 2 [Source:HGNC Symbol;Acc:HGNC:5008] | 0.034 | 19:M:0.626;32:E:0.560;227:K:0.896;313:N:0.979* |
| ENSBTAT00000037934 | *CRYBG2* | crystallin beta-gamma domain containing 2 [Source:HGNC Symbol;Acc:HGNC:17295] | 0.005069 | 10:R:0.673;51:Q:0.839;1021:C:0.681 |
| ENSBTAT00000025418 | *COL7A1* | collagen type VII alpha 1 chain [Source:HGNC Symbol;Acc:HGNC:2214] | 0.007 | 455:H:0.523;724:D:0.512;767:I:0.714 |
| ENSBTAT00000026918 | *NIPAL4* | NIPA like domain containing 4 [Source:HGNC Symbol;Acc:HGNC:28018] | 0.035706 | 22:Q:0.835;261:I:0.690 |
| ENSBTAT00000002834 | *C16orf89* | chromosome 16 open reading frame 89 [Source:HGNC Symbol;Acc:HGNC:28687] | 0.007229 | 31:D:0.961*;190:Q:0.566 |
| ENSBTAT00000009269 | *CLDN23* | claudin 23 [Source:HGNC Symbol;Acc:HGNC:17591] | 0.016565 | 102:R:0.556;259:E:0.952* |
| ENSBTAT00000005993 | *CD207* | CD207 molecule [Source:HGNC Symbol;Acc:HGNC:17935] | 0.029101 | 95:R:0.878;122:K:0.597;199:I:0.590;225:I:0.533 |
| ENSBTAT00000010989 | *TMPRSS13* | transmembrane protease, serine 13 [Source:HGNC Symbol;Acc:HGNC:29808] | 0.03394 | 88:V:0.945;93:K:0.618;132:K:0.609;173:I:0.535;176:Y:0.624;187:E:0.578;245:V:0.604;281:I:0.557 |
| ENSBTAT00000036393 | *KRT14* | keratin 14 [Source:HGNC Symbol;Acc:HGNC:6416] | 0.005 | 5:Y:0.712;14:G:0.893 |
| ENSBTAT00000027421 | *PRDM6* | PR/SET domain 6 [Source:HGNC Symbol;Acc:HGNC:9350] | 0.042559 | 30:G:0.631;264:Q:0.769 |
| ENSBTAT00000023514 | *RIN1* | Ras and Rab interactor 1 [Source:HGNC Symbol;Acc:HGNC:18749] | 0.018395 | 46:E:0.765;68:S:0.537;277:A:0.542 |
| ENSBTAT00000015892 | *CYP4B1* | cytochrome P450 family 4 subfamily B member 1 [Source:HGNC Symbol;Acc:HGNC:2644] | 0.000153 | 36:D:0.681;92:Q:0.539;163:K:0.942;186:E:0.680;206:A:0.766;229:Q:0.865;238:L:0.948 |
| ENSBTAT00000034040 | *CTRC* | Chymotrypsin-C [Source:UniProtKB/Swiss-Prot;Acc:Q7M3E1] | 0.029915 | 129:T:0.519;163:Y:0.982*;227:N:0.586;229:D:0.591 |

| | | | | |
|---|---|---|---|---|
| ENSBTAT00000004659 | *PSD* | Pleckstrin and Sec7 domain containing [Source:NCBI gene;Acc:523124]. | 0.003226 | 145:A:0.832;148:G:0.878;198:L:0.915;233:P:0.665 |
| ENSBTAT00000029024 | *SLC4A9* | anion exchange protein 4 [Source:RefSeq peptide;Acc:NP_001179135] | 0.045573 | 39:P:0.532;96:Q:0.575;541:L:0.582;568:I:0.564 |
| ENSBTAT00000063014 | *SH2D7* | SH2 domain containing 7 [Source:HGNC Symbol;Acc:HGNC:34549] | 0.005587 | 89:M:0.531;120:S:0.922;121:N:0.794;200:S:0.697;204:G:0.590;231:M:0.593 |
| ENSBTAT00000053674 | *LYPD5* | LY6/PLAUR domain containing 5 [Source:HGNC Symbol;Acc:HGNC:26397] | 0.035067 | 29:S:0.992** |
| ENSBTAT00000018189 | *WSCD2* | WSC domain containing 2 [Source:HGNC Symbol;Acc:HGNC:29117] | 0.008574 | 447:R:0.738 |
| ENSBTAT00000001458 | *SERPINB10* | serpin family B member 10 [Source:HGNC Symbol;Acc:HGNC:8942] | 0.007288 | 63:P:0.712;138:K:0.917;161:S:0.760;248:S:0.597 |
| ENSBTAT00000027887 | *NOD2* | Nucleotide-binding oligomerization domain-containing protein 2 [Source:UniProtKB/Swiss-Prot;Acc:Q6E804] | 0.012411 | 224:I:0.800;333:Q:0.842;443:I:0.559;502:D:0.714;647:A:0.575;708:M:0.534;712:I:0.558 |
| ENSBTAT00000013898 | *EVPL* | envoplakin [Source:HGNC Symbol;Acc:HGNC:3503] | 0.049555 | 79:R:0.543;226:H:0.594;480:H:0.571;532:E:0.558;654:Q:0.596;687:K:0.555;702:K:0.590;718:L:0.580;740:E:0.797;750:E:0.501;754:R:0.988*;810:T:0.568;1033:L:0.659;1057:I:0.591;1194:Q:0.550;1250:A:0.617;1318:G:0.594;1325:E:0.586 |
| ENSBTAT00000005477 | *WDR66* | WD repeat domain 66 [Source:HGNC Symbol;Acc:HGNC:28506] | 0.02947 | 79:S:0.598;177:P:0.749 |
| ENSBTAT00000027052 | *TMPRSS11A* | transmembrane protease, serine 11A [Source:HGNC Symbol;Acc:HGNC:27954] | 0.011 | 28:K:0.844;29:D:0.911;33:M:0.654;60:R:0.811;65:A:0.892;81:R:0.911;104:A:0.896;134:V:0.967*;169:H:0.608;195:V:0.902;300:K:0.904;310:T:0.780 |
| ENSBTAT00000010648 | *ACER1* | alkaline ceramidase 1 [Source:HGNC Symbol;Acc:HGNC:18356] | 0.03952 | 17:P:0.992**;129:L:0.664;158:F:0.717;167:L:0.719 |
| ENSBTAT00000020102 | *SLC16A1* | solute carrier family 16 member 1 [Source:HGNC Symbol;Acc:HGNC:10922] | 0.02755078 | 21:I:0.947;41:F:0.639;139:G:0.665;271:T:0.695 |
| ENSBTAT00000049319 | *F2RL1* | F2R like trypsin receptor 1 [Source:HGNC Symbol;Acc:HGNC:3538] | 0.042478702 | 164:L:0.654;295:G:0.579;309:G:0.981* |
| ENSBTAT00000064532 | *GPX2* | glutathione peroxidase 2 [Source:HGNC Symbol;Acc:HGNC:4554] | 0.030667882 | 130:Q:0.941 |

**Supplementary Table 23 |** Data statistics of eight samples from the ruminal and esophageal epithelium cells of four 60-day sheep embryos.

| Type | Sample | Age | Total reads | Fastq1 Adapter reads rate (%) | Fastq2 Adapter reads rate (%) | Organism | Sample ID |
|---|---|---|---|---|---|---|---|
| ATAC-seq | Rumen-1 | 60-day Embryo | 180,331,144 | 69.90% | 71.90% | *Ovis Aries* | SRR10426652 |
| ATAC-seq | Rumen-2 | 60-day Embryo | 208,760,182 | 77.10% | 75.80% | *Ovis Aries* | SRR10426651 |
| ATAC-seq | Rumen-3 | 60-day Embryo | 215,723,738 | 68.30% | 70.40% | *Ovis Aries* | SRR10426647 |
| ATAC-seq | Rumen-4 | 60-day Embryo | 169,531,318 | 78.30% | 76.90% | *Ovis Aries* | SRR10426646 |
| ATAC-seq | Esophagus-1 | 60-day Embryo | 162,012,032 | 70.60% | 70.50% | *Ovis Aries* | SRR10426645 |
| ATAC-seq | Esophagus-2 | 60-day Embryo | 182,267,094 | 74.50% | 76.30% | *Ovis Aries* | SRR10426644 |
| ATAC-seq | Esophagus-3 | 60-day Embryo | 147,260,046 | 77.40% | 77.00% | *Ovis Aries* | SRR10426643 |
| ATAC-seq | Esophagus-4 | 60-day Embryo | 166,895,578 | 77.00% | 76.60% | *Ovis Aries* | SRR10426642 |

**Supplementary Table 24 |** Gene list of 243 rumen key genes which have RSCNEs nearby overlapping with open accessible peaks.

[see the excel file]

**Supplementary Table 25 |** GO enrichment analysis of 243 rumen key genes which have RSCNEs nearby overlapping with open accessible peaks. BP denotes Biological Process, MF denotes Molecular Function, and CC denotes Cellular Component.

| ID | Type | Description | *p* value | adjusted *p* value | list1InGO | list1OutGO | list2InGO | list2OutGO |
|---|---|---|---|---|---|---|---|---|
| GO:0005198 | MF | structural molecule activity | 2.60E-09 | 4.43E-05 | 19 | 157 | 357 | 12176 |
| GO:0005882 | CC | intermediate filament | 5.38E-20 | 9.16E-16 | 12 | 164 | 80 | 12453 |
| GO:0005911 | CC | cell-cell junction | 3.60E-11 | 6.14E-07 | 18 | 158 | 285 | 12248 |
| GO:0009913 | BP | epidermal cell differentiation | 2.83E-23 | 4.82E-19 | 9 | 167 | 37 | 12496 |
| GO:0030216 | BP | keratinocyte differentiation | 3.30E-17 | 5.62E-13 | 7 | 169 | 30 | 12503 |
| GO:0043392 | BP | negative regulation of DNA binding | 3.75E-13 | 6.39E-09 | 6 | 170 | 29 | 12504 |
| GO:0044459 | CC | plasma membrane part | 4.94E-09 | 8.42E-05 | 41 | 135 | 1220 | 11313 |
| GO:0045095 | CC | keratin filament | 3.75E-13 | 6.39E-09 | 6 | 170 | 29 | 12504 |
| GO:0045604 | BP | regulation of epidermal cell differentiation | 1.46E-13 | 2.49E-09 | 6 | 170 | 28 | 12505 |
| GO:0045682 | BP | regulation of epidermis development | 1.59E-15 | 2.71E-11 | 8 | 168 | 45 | 12488 |
| GO:0098609 | BP | cell-cell adhesion | 6.12E-16 | 1.04E-11 | 5 | 171 | 15 | 12518 |

**Supplementary Table 26 |** Differentially accessible peaks (DAPs) between eight ATAC-seq from the ruminal and esophageal epithelium cells of four sheep 60 days embryos.

[see the excel file]

**Supplementary Table 27 |** Gene list of 22 rumen key genes have nearby Rumen-specific DAP-associated RSCNEs.

| ***Gene Name*** | | | | | | |
|---|---|---|---|---|---|---|
| *HPN* | *SLC28A3* | *PROM2* | *LOC101107795* | *HES1* | *DAB1* | *SLC26A3* |
| *EPHA4* | *SLC9A3* | *MSX2* | *LOC101104661* | *PFKFB3* | *TNFAIP3* | *CPA6* |
| *LOC105606881* | *LOC105604781* | *LOC101121219* | *TMEM177* | *DMRT2* | *FXYD3* | *WDR66* |
| *HPSE2* | | | | | | |
